# Bringing Attomolar Detection to the Point-of-Care with Nanopatterned DNA Origami Nanoantennas

**DOI:** 10.1101/2024.10.14.618183

**Authors:** Renukka Yaadav, Kateryna Trofymchuk, Mihir Dass, Vivien Behrendt, Benedikt Hauer, Jan Schütz, Cindy Close, Michael Scheckenbach, Giovanni Ferrari, Leoni Maeurer, Sophia Sebina, Viktorija Glembockyte, Tim Liedl, Philip Tinnefeld

## Abstract

Creating increasingly sensitive and cost-effective nucleic acid detection methods is critical for enhancing point-of-care (POC) applications. This involves capturing all desired biomarkers in a sample with high specificity and transducing the capture events to a detector. However, the signal from biomarkers present at extremely low amounts often falls below the detection limit of typical fluorescence-based methods, making molecular amplification a necessary step. Here, we present a nucleic acid detection assay of a 151-nucleotide sequence specific to antibiotics-resistant *Klebsiella pneumoniae*, based on single-molecule fluorescence detection of non-amplified DNA down to the attomolar level, using Trident NanoAntennas with Cleared HOtSpots (NACHOS). Our NACHOS-diagnostics assay leverages a compact microscope with a large field-of-view and cost-efficient components, including microfluidic flow to enhance capturing efficiency. Fluorescence enhancement is provided by DNA origami NanoAntennas, arranged in a dense array using a combination of nanosphere lithography and site-specific DNA origami placement. Our method can detect 200 ± 50 out of 600 molecules in a 100 µL sample volume within an hour. This represents typical number of pathogens in clinical samples commonly detected by Polymerase Chain Reaction but without the need for molecular amplification. We achieve similar sensitivity in untreated human blood plasma, enhancing the practical applicability of the system. Our platform can be adapted to detect shorter nucleic acid fragments that are not compatible with traditional amplification-based technologies. This broadens its potential for diverse diagnostic and healthcare applications, providing a robust and scalable solution for sensitive nucleic acid detection in various clinical settings.

## Introduction

Single-molecule methods are gaining ground in bioanalytical applications like nucleic acid sequencing and sensitive analyte detection.^1–4^ For Point-of-Care (POC) detection in low-technology environments, however, single-molecule approaches, are still considered prohibitively expensive due to their reliance on sophisticated setups. For instance, single-molecule detection using fluorescence often requires either molecular multiplication to detect the target’s signal against background noise or expensive instrumentation to detect single fluorescent molecules directly.^1,5,6^ Advances in plasmonic fluorescence enhancement,^7–9^ utilizing DNA origami nanostructures,^10–12^ have facilitated signal amplification of fluorophores captured in the hotspot, improving the distinction of the real signal from background impurities and enabling single-molecule detection using a portable, battery-driven smartphone microscope.^13^ But their use for target detection at clinically relevant nucleic acid concentrations below the picomolar range^14^ has remained elusive.

Arguably, analytical methods cannot get more sensitive than detecting a single molecule and concentration determination cannot become more direct than digitally counting all molecules in a sample.^2,15–17^ Single-molecule detection is thus swiftly moving towards the development of affordable, portable devices, making it accessible outside of specialized laboratories and into POC and field settings.^13,18–23^ The challenge here is not to detect the single molecule present in a (typically very small) detection volume, but to find all molecules in the patient sample. At 1 aM (10^−18^ mol/L) concentration, 100 µL of blood serum contains ∼60 molecules that need to be detected. It is impractical to rely solely on Brownian motion to transport these molecules through the minuscule detection volumes of e.g. a focused laser beam or a nanopore.^1^ To address this, most detection methods rely on a pre-concentration step^2,24,25^ or utilize molecular amplification strategies^26–28^. For nucleic acids, dPCR (digital polymerase chain reaction) has been a game changer. dPCR works by partitioning the sample into many reaction chambers or droplets and diluting until each chamber contains one or zero target molecule. The molecules are then amplified by thermal cycling, and the positive droplets are detected using fluorescence-based methods, determining an absolute number of target nucleic acids.^20^ However, practical use of commercially available dPCR devices, especially in low-resource settings is still limited by complex workflow, need for trained personnel and advanced equipment.^29^

Building on addressable NanoAntennas with Cleared HOtSpots (NACHOS)^13^ we present NACHOS-diagnostics, a POC-compatible amplification-free detection approach for nucleic acids that addresses these challenges. Specifically, we target a synthetic DNA sequence specific to carbapenem-resistant *Klebsiella pneumoniae*, a bacterium which has been directly linked to 1.27 million deaths and contributing to 4.95 million deaths annually.^30^ The fluorescence enhancement by NanoAntennas not only facilitates single-molecule detection with simple optics but it also creates a contrast against an unavoidable background of single-molecule impurities, thus minimizing false positive signals.^13^ NACHOS diagnostics utilizes a sandwich assay^13^ with a capturing sequence and a dye-labeled imager strand to detect DNA target strands over a broad concentration range from attomolar to nanomolar.

## Results

At nucleic acid concentrations where only a few molecules are present in the sample volume, detection systems face significant challenges−background noise from impurities, sensitivity of the assay and long response times.^24^ Employing NanoAntennas allows us to tackle the first challenge by physically amplifying the fluorescence signal of a fluorophore captured in the hotspot of plasmonic nanoparticles (NPs). Resolving the latter two challenges requires maximizing the probability of capturing the target molecules in the shortest possible time. We address this with an integrated NACHOS-diagnostic approach involving several steps of development (Figure 1). These include optimization of the DNA origami NanoAntenna design; integrating nanopatterning and microfluidics; and engineering of a fluorescence reader with single-molecule detection software.

**Figure 1:**
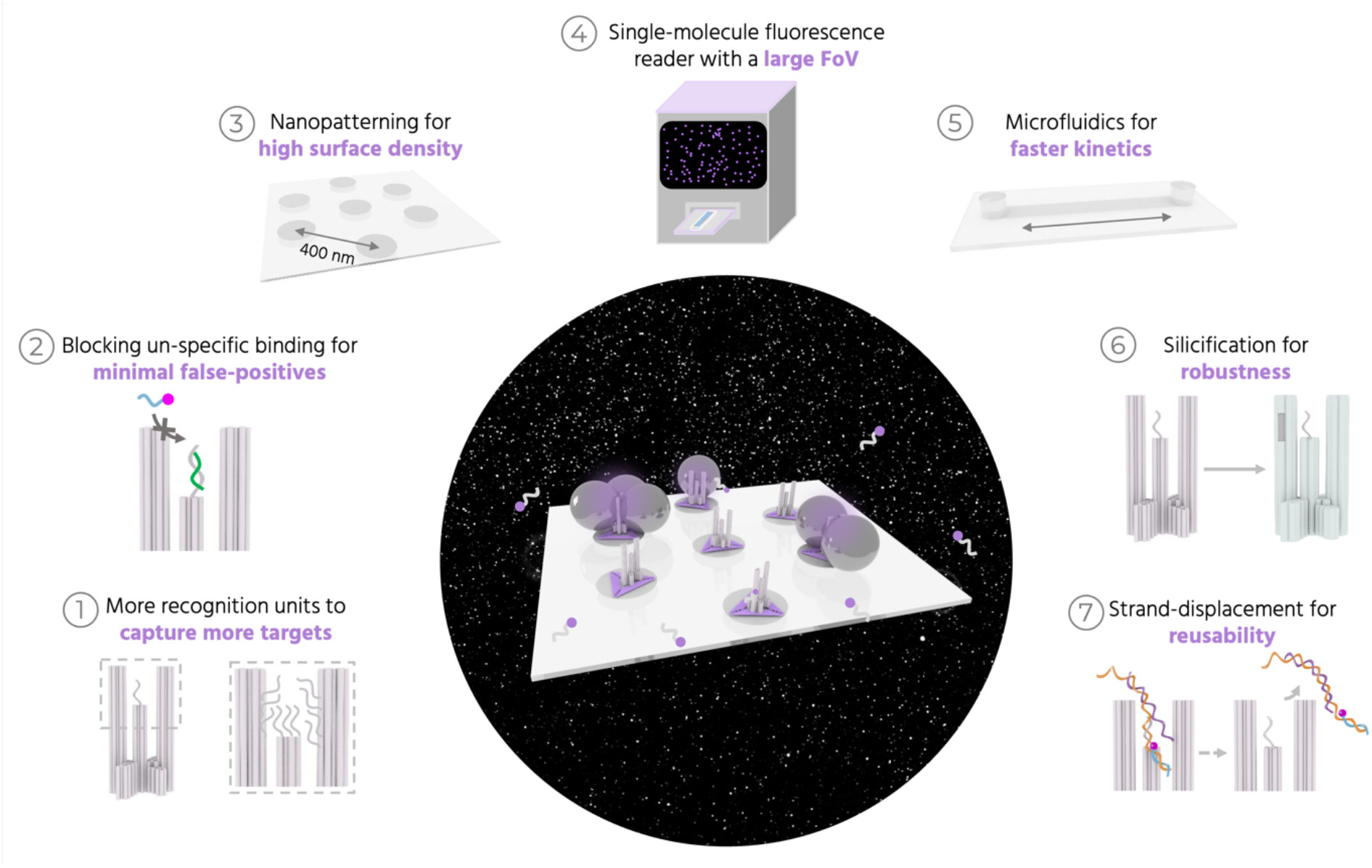
Overview of the different components for NACHOS diagnostics. Different components involved in our approach to develop fluorescence enhanced single-molecule detection into a POC-compatible method for detecting low target concentrations of nucleic acids. An abstract image in the center showing a purple glow upon capturing DNA in the NanoAntennas with a snapshot of thousands of single molecules detected on our fluorescence reader as the background. Panels 1-7 show the different steps we employed, to address the challenges in achieving high sensitivity.

### NanoAntenna design and nucleic acid assay optimization

We use the second generation of NACHOS^31^ for the NanoAntenna assembly as its larger gap (19 nm) between side pillars compared to the first generation (6.5 nm^13^), allows more room for integrating multiple capture strands (Figure 1, panel 1). The DNA origami structure, called *Trident*, consists of a central pillar (51 nm in height) flanked on two sides by longer pillars (74 nm in height). The pillars emerge from a 44 nm wide cross-shaped base (Figure 2a and Figure S1). We modify the base of the Trident to extend twelve biotinylated single-stranded DNA (ssDNA) extensions from the bottom face, which are used to immobilize the structure on glass coverslips using Biotin-NeutrAvidin linkages (Figure 2a). The sandwich-type hybridization assay^13^ involves four strands – capture, target, imager, and blocker (Figure 2b). The ‘capture’ strands are extended staple strands that protrude from the Trident and hybridize to the ‘target’ strand (which is 151 nt long) via a 17 nucleotide (nt) complementary sequence. The ‘imager’ strand, labelled with Alexa Fluor 647, has a 17 nt sequence complementary to another part of the target. The ‘blocker’ strand is shorter (10 nt) and is complementary to a portion of the capture strand. It prevents unspecific interaction between the imager and capture strands in the absence of the target (Figure 2b, Figure S2 and Figure 1, panel 2). The target, when added, can displace the blocker (Figure 2b). We utilize a synthetic target sequence specific to the OXA-48 gene, used for diagnosing carbapenem-resistant *Klebsiella pneumoniae* infection (Supplementary Note 1.1).^32^ The gram-negative bacterium can cause various infections, including pneumonia and others of the bloodstream and urinary tract.^33^

**Figure 2.**
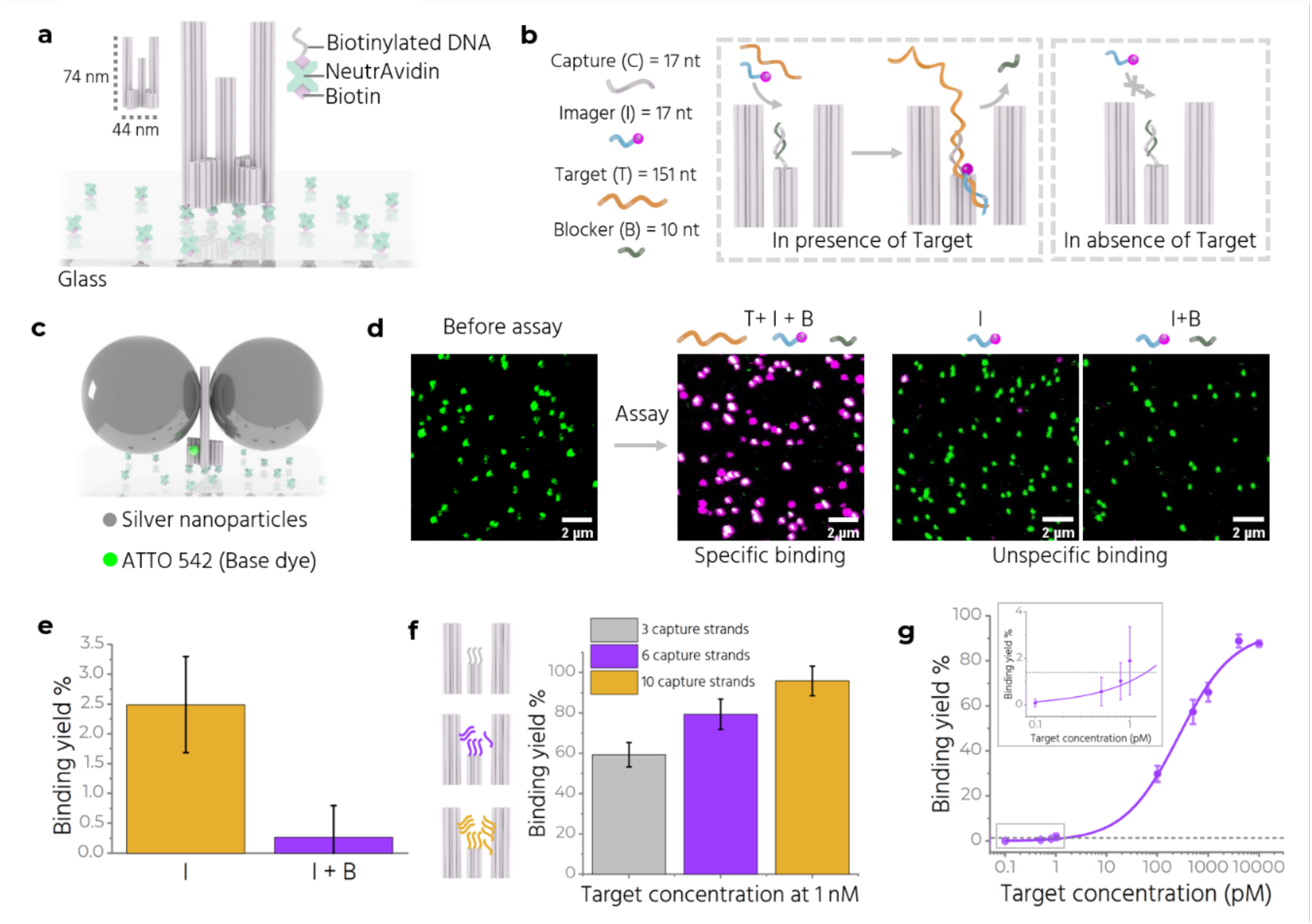
Optimization of nucleic acid assay detection with NACHOS. **a** Sketch of the Trident nanostructure immobilized on a BSA-Biotin-NeutrAvidin modified glass coverslip. **b** Working principle of the assay in presence and absence of target. A single capture strand is shown for simplified visualization. **c** Sketch of a fully assembled Trident NACHOS with 80 nm silver nanoparticles and an ATTO 542 dye at the base. **d** Confocal fluorescence scans before and after the assay. Scale bars: 2 µm. **e** Binding yield or colocalization percentage comparison after incubation with imager only (in yellow) and with imager and blocker (in purple). **f** (Left, insets) Schematic showing the positions of 3, 6 and 10 capture strands on the Trident. (Right) Target binding yield in the three cases upon incubation with target, imager and blocker. **g** Target binding yield at varying target concentrations. Inset shows a magnified view between 0.1 - 1 pM target concentration. For each data point, at-least 3 (20 µm x 20 µm) confocal scans with at-least 300 molecules per scan are analyzed. The error bars represent standard deviation. The purple line represents logistic fit. The grey dashed line indicates the minimum detectable target concentration above the blank.

We use 80 nm silver nanoparticles (AgNPs) functionalized with thiolated T_20_ strands for plasmonic enhancement. We modify each side pillar to incorporate six A_20_ extensions to capture two AgNPs, one on each pillar, assembling the NanoAntenna (Figure 2c). An ATTO 542 at the base of the Trident acts as an internal reference to determine the position of the nanostructure during confocal measurements (Figure 2c). Detailed characterization of the NanoAntenna is included in the supporting information (Figure S3). We assess the specificity and efficiency of the assay by taking single-molecule confocal fluorescence scans before and after performing the assay. Initially, we observe only green spots corresponding to the reference ATTO 542 (Figure 2d). After incubation with target (4 nM), imager (12 nM), and blocker (12 nM) strands for one hour at 37°C and subsequent washing, we detect magenta (absence of reference dye due to limited labelling efficiency) and white spots (colocalization of two dyes on the structure) (Figure 2d). To quantify the assay, we calculate the specific ‘binding yield’ as

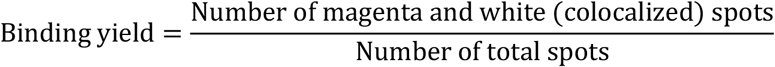

To quantify ‘unspecific binding’, we perform the assay without the target. We obtain fewer colocalized spots when both imager and blocker are present compared to imager alone (Figure 2d and 2e).

As predicted, the larger gap between the two side pillars in the Trident allows integration of more capture strands in the hotspot region. Increasing the number of capture strands increases the binding yield (Figure 2f). Bleaching step analysis confirms that a single Trident NACHOS with 10 capture strands can capture up to 8 target strands at higher concentrations (4 nM) (Figure S4). This supports the increased accessibility of the hotspot in the current design compared to the previous design, where more than 70% structures captured only a single target strand.^13^ We also examine whether including more capture strands affects nanoparticle binding due to steric hindrance. We compare Trident variants with zero and 10 capture strands, and do not observe a loss in enhancement with the introduction of more capture strands (Figure S5). Hence, we employ 10 capture strands for subsequent experiments. Next, we measure the target binding yield at varying target concentrations from 100 fM to 10 nM (Figure 2g). The imager and blocker concentrations were kept constant at 12 nM for each sample (See Figure S6 for example fluorescence scans). In this setting, the limit of detection (LoD) for the assay is determined to be ∼1 pM using the formula:

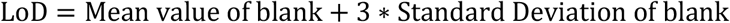

### Optimizing the cross-section for binding with DNA origami NanoAntenna placement

Besides increasing the cross-section for target binding by increasing the number of capture strands per NanoAntenna, we aim next at increasing the surface density of NanoAntennas. The challenge here is to achieve dense placement of NanoAntennas on a substrate without them interacting with each other and aggregating into clusters with uncontrolled properties.^34^ We adapt the combination of DNA origami placement (DOP)^35–37^ and nanosphere lithography^38^ to regularly arrange NanoAntennas at 400 nm distances (Figure 1, panel 3). The method involves drop-casting polystyrene (PS) nanospheres (400 nm diameter) onto a hydrophilic coverslip and then passivating the surface to render the unmasked areas hydrophobic. The spheres are then lifted off, revealing a hexagonal array of hydrophilic *placement sites* (Figure 3a). DNA origami nanostructures are selectively placed on these sites through electrostatic interactions. A high-quality DOP, defined as one origami per placement site, is achieved by optimizing Mg^2+^ concentration, pH, incubation time, and DNA origami concentration (Figure 3b). Our modified DOP protocol involves two placement steps. First, a two-dimensional *Triangle* origami is placed on the substrate. We use a modified version of the ‘Rothemund triangle’ with a side length of ∼127 nm (Figure S7),^39^ which corresponds well to the placement sites of ∼120 nm diameter.^35^ Second, the Trident is introduced with six ssDNA strands extending from its bottom face that are complementary to the six protruding sequences on the Triangle. The two-step placement prevents direct interaction of the Trident with placement sites, which otherwise results in the placement of multiple Tridents per placement site in random orientations (Figure S8). We use atomic force microscopy (AFM) (Figure S9) and DNA-PAINT (Point Accumulation In Nanoscale Topography)^40,41^ to characterize the quality of placement. DNA-PAINT, as a super-resolution microscopy technique, allows single-molecule localization of freely diffusing short, labelled DNA probes (imager) transiently binding to complementary ssDNA (docking strand) on the origami. We incorporate six docking strands (60 nm apart) (Figure S10) on the outer rim of the Triangle and perform DNA-PAINT imaging with an 8 nt imager labelled with ATTO 655. We observe a hexagonal placement pattern with Triangles placed ∼400 nm apart (Figure 3c). We also observe geometric defects consistent with those observed in scanning electron microscopy (SEM) images of self-assembled nanospheres on glass (Figure S11).

**Figure 3.**
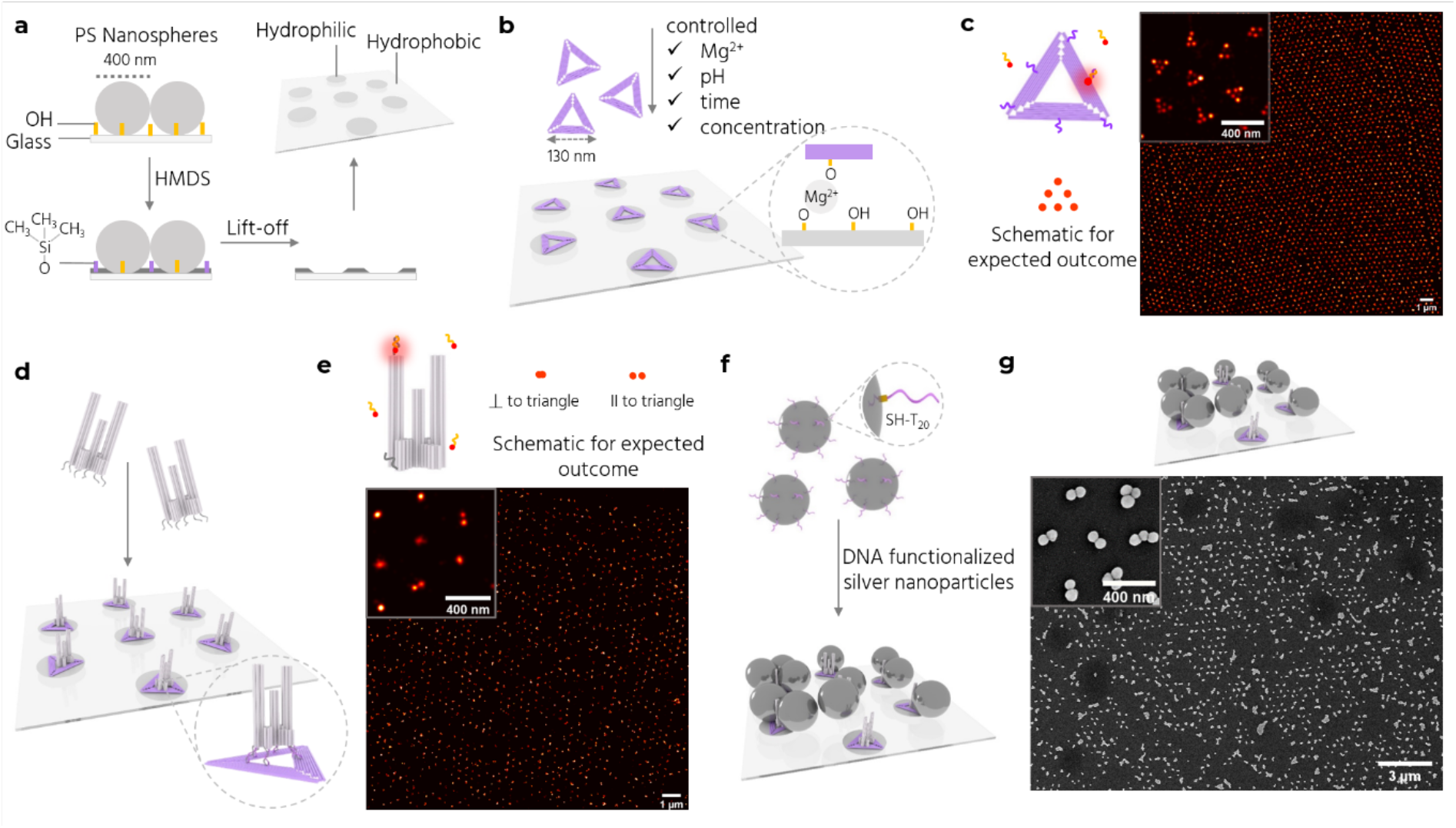
Site-specific nanopatterning of Trident NanoAntennas. **a** Fabrication of a nanopatterned surface via nanosphere lithography involving nanospheres deposition, vapor-phase passivation by HMDS, and lift-off. **b** DOP of Triangle DNA origami nanostructures on the hydrophilic placement sites through electrostatic interactions between the placement sites and the Triangle (inset). **c** (Left) Triangle with 6 docking sites (purple) for DNA-PAINT experiments and a schematic of the expected outcome. (Right) DNA-PAINT image of the triangle placed on a nanopatterned surface with a zoom-in (inset). **d** DNA hybridization between the Trident and the Triangle. **e** Trident with 2 docking sites (dark gray) for DNA-PAINT experiments and a schematic of the expected outcome (top) for upright Tridents (correct orientation) or fallen Tridents (undesired orientation) on the Triangle. A single strand per docking site is shown for simplicity. DNA-PAINT image of the Trident bound to the Triangle is shown with a zoom-in (inset). **f** Incubation with functionalized AgNPs (dark gray spheres) results in full NanoAntenna assembly. g SEM image of NanoAntennas on a patterned surface with a zoom-in (inset).

After the Triangles are placed, we add the Trident. The extensions at the base of the Trident hybridize with the protrusions on the Triangle (Figure 3d). The extensions are placed at the inner hole, protruding in the plane of the Triangle, to make sure they are accessible irrespective of which face of the Triangle lands on the placement site. We use DNA-PAINT to study the orientation of the Tridents after placement by incorporating six docking strands (three near the top and three near the bottom of one outer pillar). Tridents in the desired upright orientation are observed as one overlapping spot, while those lying parallel to the surface show up as two spots (Figure 3e). We achieve more than 80% upright Tridents after optimizations (Figure S12).

Next, we incubate the patterned Tridents with thiolated-DNA functionalized 80 nm AgNPs to complete the NanoAntenna assembly (Figure 3f). Under ideal conditions, each NanoAntenna would contain two AgNPs to create a ‘dimer’. To achieve this, it is imperative to refine the functionalization protocol to minimize NP aggregation (Figure S13). Although aggregation does not compromise the assay and can even yield elevated enhancement factors,^42^ it poses a challenge for achieving uniform, closely spaced assemblies. We employ the freeze and thaw functionalization method to increase the DNA loading on each NP, ^43^ followed by an agarose gel-based purification step to separate single AgNPs from aggregates and free thiol-DNA strands. SEM imaging reveals the hexagonal array of NanoAntennas on the coverslips (Figure 3g and S14). As a control, we perform DOP with Triangles and incubate the surface with functionalized NPs, observing minimal or no binding. This confirms the selective binding of NPs to the Trident (Figure S15). Adding Tridents directly to a surface with empty placement sites results in the placement of multiple Tridents per placement site, many of which are misoriented or have fallen. Adding AgNPs to this sample results in minimal binding or aggregation (Figure S16). This suggests that it is crucial to have at least one Trident bound to the Triangle in the desired orientation for proper assembly of the NanoAntenna. Additional Tridents bound to the Triangle in random orientations are unable to capture NPs to form the plasmonic hotspot due to steric hindrance and are rendered unfunctional.

### Single-molecule fluorescence reader

To enable POC detection of single molecules on a cost-efficient device that also provides a large field-of-view (FOV) to count all the single molecules captured by NanoAntennas, we developed a fluorescence reader (Figure 4a and Figure 1, panel 4). The reader uses two spectrally filtered LEDs to excite the Alexa Fluor 647 and a CMOS camera for detection. It is sensitive enough to detect single molecules when their fluorescence is enhanced by NanoAntennas (Figure 4b and S17). At our standard settings of 2 W/cm^2^ excitation power density and an integration time of 300 ms, single molecules not enhanced by NanoAntennas remain dark. This creates an intrinsic filter against false positive signals, as unspecific binding in the ultra-small volumes of the NanoAntenna hotspot is minimal, whereas an unavoidable background of not-enhanced single-molecule impurities is invisible.^13^ The reader also includes two white LEDs to collect scattering light from NPs. An integrated computer with a touchscreen serves as the user interface, running a custom software that can control all components and execute script-based processes. The software also performs image processing to detect single molecules based on their intensity and temporal blinking behavior. A detailed description of the reader is provided in Supplementary Note 2.

**Figure 4.**
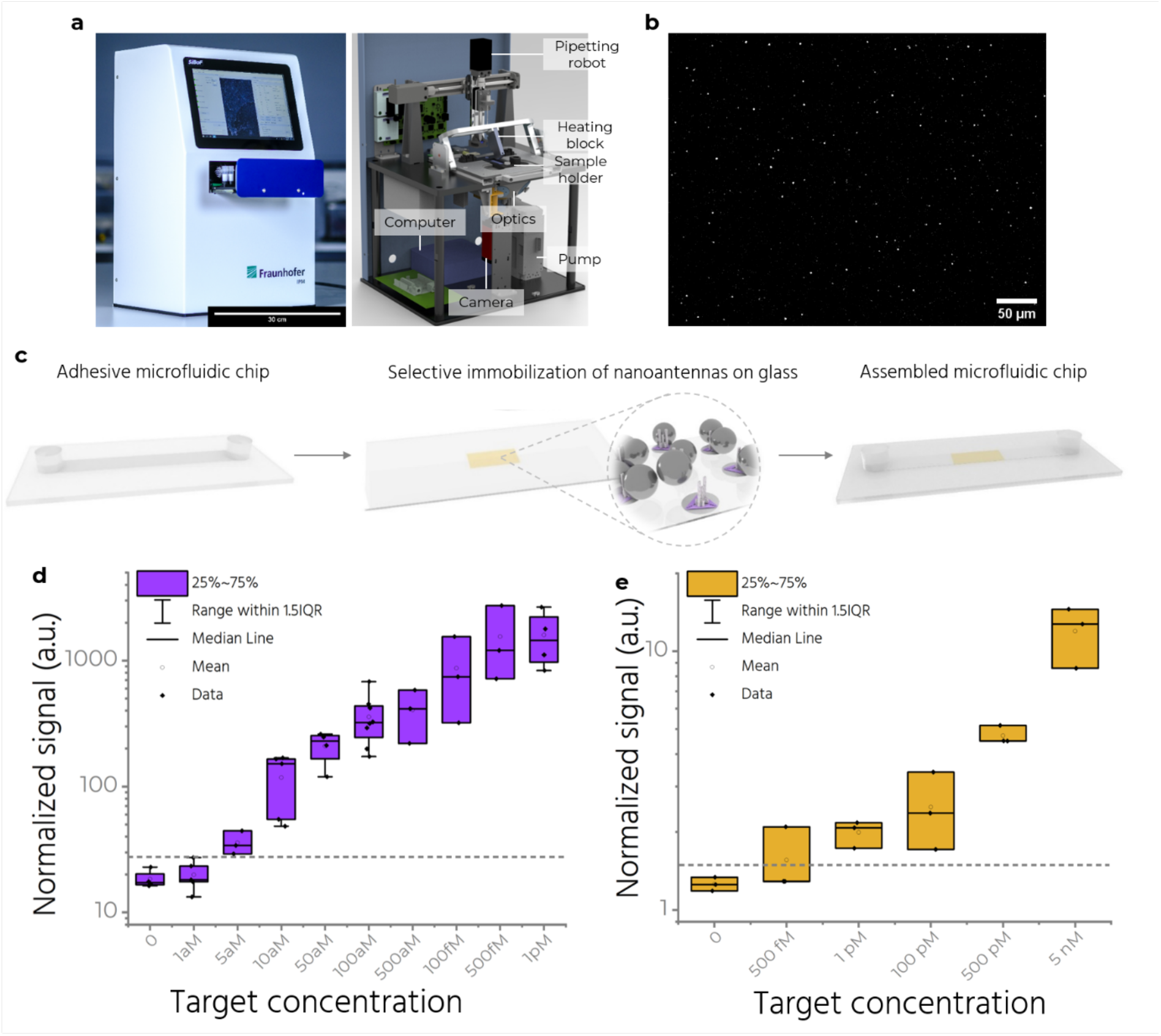
Microfluidics and a simple fluorescence reader. **a** (left) The single-molecule fluorescence reader and (right) a sketch showing its components. Scale bar: 30 cm. **b** A zoom-in view of an exemplary image captured on the reader after sandwich assay with target concentration of 1 pM. Scale bar: 50 µm. **c** A self-adhesive microfluidic chip is attached on top of the coverslip with patterned NanoAntennas. The yellow rectangle depicts the area where DOP of NanoAntennas is performed. **d** Measurements showing efficiency of the assay at target concentrations from 1 aM to 1 pM, performed on the reader and analyzed by spot picking. The grey dashed line indicates the minimum target concentration detectable above the blank. **e** Measurements on the reader analyzed by intensity averaging between target concentrations of 500 fM-5 nM. The box plots in d and e show the 25/75 percentiles and the whisker represents the 1.5*IQR (inter quartile range) length, the center lines represent the average values. The grey dashed line indicates the minimum target concentration detectable above the blank in d and e.

For finding all target molecules in our 100 µL sample solutions, the reader features a field-of-view (FOV) of 3 mm x 2.5 mm. This matches the self-adhesive microfluidic chip that features a channel (Width: 2.5 mm, Depth: 150 µm, Length: 58.5 mm), inlet and outlet ports for easy pipetting, and can attach to glass substrates (Figure 4c). This selective placement (Figure S18) allows us to detect most of what we capture, minimizing the loss of target strands outside the detection FOV. Besides the high density of capturing strands and the increased density of NanoAntennas, the large FOV further enhances the interactions of sample volume with the NanoAntennas. Next, we employ fluidics housed in the reader-a simple, automated repetitive back and forth flow within the narrow channel in the microfluidic chip, to further increase the binding kinetics beyond purely diffusive interactions (Figure 1, panel 5). Applying flow allows us to achieve a ∼10-fold increase in target capture. We demonstrate this by comparing assays on chips patterned with NanoAntennas and comparing the two cases—one performed with flow and the other without (Figure S19), both with an incubation time of one hour and a target concentration of 500 fM.

### Achieving attomolar sensitivity

Combining the various elements (Figure 1, panels 1-5) into the NACHOS chip, we use the reader to characterize assays performed at target concentrations ranging from 1 aM to 5 nM, achieving an LoD of ∼5 aM (Figure 4e and S20).

We determine this by counting the number of spots detected in each sample normalized to the Nanoantenna surface density (see Supplementary Note 2, Normalized signal). Measurements were repeated at least three times to obtain a standard deviation. In our reader, with an optical resolution of ∼ 2.7 µm (Supplementary Note 2, reader specifications), a single molecule is detected as a spot (Point Spread Function) with a diameter of 3 pixels, with each pixel corresponding to 0.8 µm. Within the aM-fM concentration range, the probability of spatially overlapping signals from two different target molecules remains low, so each detected spot is assumed to correspond to a single molecule. Specifically, at 1 pM, we estimate the average proportion of double molecules among the detected spot to be 0.5-1.5% (Supplementary Note 2, double-event estimate). Thus, throughout the aM–fM range, the proportion of unaccounted double molecules is safely below 1%. At concentrations above 1 pM, the spots are too closely spaced to accurately distinguish them individually (Figure E, Supplementary Note 2). We therefore use an intensity-based analysis - which calculates an average intensity value over the whole image – to analyze data from 500 fM-5nM. Using average intensity alone, we calculate an LoD of approximately 1 pM, further highlighting the advantage of single-molecule counting over averaging. It is worth noting that the combination of the two detection strategies (in their respective ranges of applicability) preserves accuracy in the full concentration range, from aM to nM regime.^44^

### Silicification for stability in clinically relevant fluids

Biological fluids like plasma contain many components in addition to target molecules. These components can interact unspecifically with substrates and add to the noise, reducing the sensitivity of our assay. We use buffers containing monovalent electrostatic ions like Na^+^ to reduce such interactions and wash away weakly-bound unspecific molecules (Figure S21). But the same Na^+^-containing buffers also lift-off the DNA origami structures placed through DOP (Figure S22). To overcome these issues, we employ silicification (Figure 1, panel 6), which involves the formation of a robust silica coating on the DNA origami nanostructure. The inorganic coating significantly enhances the stability of the structure and keeps them ‘glued’ to the surface (Figure S23) over a wide range of pH, temperature and salt concentrations.^45,46^ Importantly, silicification does not affect the accessibility of ssDNA extensions (capture strands), which is crucial for the assay’s functionality (Figure 5a).^47^

**Figure 5.**
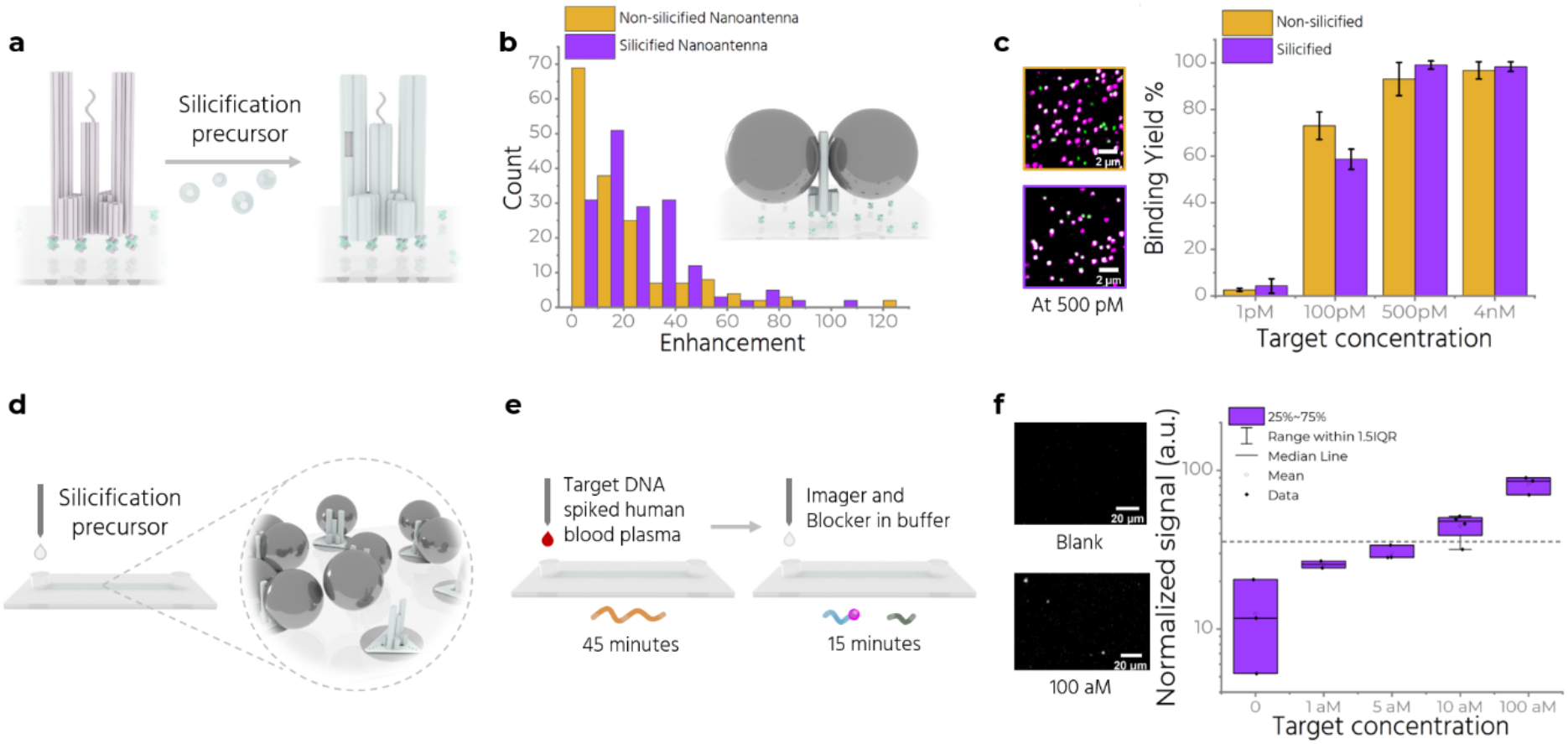
Silicification and measuring in human blood plasma. **a** Coating of the double stranded regions of the Trident with silica. **b** Enhancement values are compared for non-silicified and silicified NanoAntennas. (inset) Schematic of a silicified NanoAntenna. **c** (left) Exemplary confocal fluorescence scans for silicified and non-silicified NanoAntenna after incubating with 500 pM target concentration on the left. (right) Column plot comparing target binding yield at different target concentration in case of silicified and non-silicified NanoAntennas. **d** Silicification of nanopatterned NanoAntennas assembled in the microfluidic chip. **e** The microfluidic chip is incubated first with target spiked plasma for 45 minutes, washed with buffer and then incubated with imager and blocker for 15 minutes. **f** Measurements on the reader with target spiked plasma at target concentrations 0, 1 aM, 5 aM, 10 aM and 100 aM, with silicified NanoAntennas. Grey dashed line indicates the lowest detectable signal above blank. Zoom-in snapshots from the blank and 100 aM chip is shown on the left. Scale bar at 20 µm.

We introduce a pre-hydrolyzed precursor - prepared by mixing *N*-trimethoxylsilylpropyl-*N,N,N*-trimethylammonium chloride (TMAPS) and tetraethoxysilane (TEOS) solutions in Mg^2+^ buffer^48^ - to the surface with DNA origami, incubate for desired durations, wash with water and ethanol and air-dry to complete the process. We use AFM to compare the height of the base of the Trident prior to silicification, and after one and four days of silicification to determine optimal silicification parameters (Figure S24). We observe a thickness increase of ∼3 nm after one and ∼7 nm after four days of silicification, consistent with other studies.^47,48^ After 4 days of silicification, we could not reasonably resolve the structural features of the Trident with AFM, and observed an increase in formation of unidentified clusters. We chose the 1-day incubation period for further experiments. Next, we check the effect of silicification on fluorescence enhancement by silicifying NanoAntennas immobilized on glass using BSA-biotin-NeutrAvidin. These NanoAntennas are folded with an Alexa Fluor 647-modified staple strand which positions the dye in the hotspot, in addition to the base dye (ATTO 542). We record single-molecule confocal fluorescence scans and measure the intensity of the Alexa Fluor 647, computing a maximum enhancement factor of 126 and 102 for non-silicified and silicified NanoAntennas respectively (Figure 5b). We also check if silicifying the origami adversely affects the accessibility of the capture strand in the hotspot. Results show similar target-capture binding yields in silicified and non-silicified NanoAntennas at all tested target strand concentrations (Figure 5c). We then test the performance of silicified NanoAntennas in untreated human blood plasma. Plasma - a clear, slightly yellow liquid composed of blood without red blood cells, white blood cells and platelets – is ubiquitously used in diagnostics.

We start by silicifying NanoAntennas immobilized using BSA-biotin-NeutrAvidin chemistry. We then incubate the sample for one hour with plasma spiked with 5 nM of target and 12 nM of both imager and blocker and measure the imager and base dye colocalization (Figure S25). We observe only green spots, consistent with the base dye, indicating the presence of stable origami on the surface. Absence of other colored spots suggests degradation of the target and/or imager by enzymes in the plasma. Next, we perform an assay under the same buffer conditions by incubating with plasma spiked only with the target for an hour, wash the sample with buffer to remove the plasma, and then incubate with the imager and blocker for 15 minutes (Figure S25). We observe colocalized spots confirming proper functioning of the assay and suggesting that the imager gets quickly degraded in plasma but not the target. Additionally, we introduce 1 µM of random DNA sequence (sacrificial DNA) during the target incubation. This results in a 94.6% binding yield, an improvement over the 90.8% yield without sacrificial DNA (Figure S25). The target’s longer sequence (151 nt) and its tendency to form secondary structures^49^ make it less prone to enzymatic degradation (Figure S26). Shorter sacrificial DNA strands appear to further slow this degradation process. We adapt this strategy and incubate the NACHOS chip (Figure 5d) with target-spiked plasma for 45 minutes, put it in the reader with fluidics and wash the chip to remove any unspecifically bound residues originating from the plasma. We then re-incubate the chip with the imager and blocker for 15 minutes while performing fluidics, and wash with fresh buffer (Figure 5e). We measure target concentrations from 1 aM to 100 aM and observe the lowest detectable concentration above the blank to be ∼10 aM after silicification (Figure 5f and S27), compared to 5 aM in assays performed with unsilicified NanoAntennas. Silicification thus significantly increases the stability of the structures without impacting the sensing performance, enabling assays in clinically relevant fluids.

Finally, we use toehold-mediated strand-displacement^50^ to displace the bound target-imager duplex from the capture strand, making the capture strand accessible for the next round of use (Figure 1, panel 7 and Figure S28). As the cost of a diagnostic test often determines its accessibility to those most in need, making the chips reusable and offering users the ability to modify the same chip for detecting various target sequences can benefit POC applications.

## Conclusion

In this work, we have developed an amplification-free POC-compatible nucleic acid detection assay capable of attomolar sensitivity, targeting a 151-nucleotide sequence specific to antibiotics-resistant *Klebsiella pneumoniae*. DNA origami NanoAntennas, particularly NACHOS, excel in distinguishing signals from background noise. We merged the strengths of NACHOS with nanopatterning and microfluidics to offer an integrated approach that addressed key challenges in detecting low-abundance targets.

A nanopatterned surface densely populated with NanoAntennas combined with microfluidics to overcome the effects of slow kinetics at low target concentrations, increases the probability of target capture. We develop a single-molecule fluorescence reader boasting a larger FOV to efficiently detect captured targets. Silicification reduces the dependence of our system on ionic concentrations, pH, temperature as well as protects against degrading enzymes. Finally, a strand displacement strategy is optimized to be able to re-use our chips. We use a sandwich-type assay to detect target DNA bearing the sequence specific to a clinically relevant microbe and achieve a detection limit of 5 aM in buffer and 10 aM in untreated human blood plasma within one hour. It should be possible to modify our chip to target any DNA target by extending an ‘adapter’ sequence from the origami to which a part of a capture strand can bind, allowing the user to choose the sequence for the capture and thus the target.

Overall, our method offers a robust and scalable solution for sensitive nucleic acid detection in various clinical settings, with potential applications in diagnosing antibiotic-resistant infections, cancer biomarkers, and neurodegenerative diseases. By providing a simple, amplification-free approach with high sensitivity and specificity, our platform addresses critical needs in rapid, on-site diagnostics, potentially improving patient outcomes and aiding in the global fight against antimicrobial resistance. Finally, NACHOS-diagnostics has the potential to evolve into an alternative to digital PCR for quantifying nucleic acid molecules, as it may eliminate the need for separation into small reaction vessels.

## Methods

### DNA Origami

We design DNA origami structures using caDNAno2,^51^ assembling and purifying them with protocols adapted from Wagenbauer et al.^52^ We mix 30 nM of in-house produced p8064 scaffold for Trident and p7249 for Triangle with 300 nM unmodified staples (pooled from an original 100 µM concentration) and 750 nM modified staples (also pooled from 100 µM). Integrated DNA Technologies Europe GmbH, Germany; Eurofins Genomics GmbH, Germany and biomers.net GmbH, Germany supply all staples, and their exact sequences are in Supplementary Note 3. We add 10 x of folding buffer (FoB) (200 mM MgCl_2_, 50 mM Tris, 50 mM NaCl, 10 mM EDTA) to this mixture and subject it to a thermal annealing ramp for 16.5 hours (detailed in Supplementary Table S2). For purification, we use 100 kDa MWCO Amicon Ultra filters (Merck KGaA, Germany), washing the samples four times with a lower ionic strength buffer (5 mM MgCl_2_, 5 mM Tris, 5 mM NaCl, 1 mM EDTA) for 5 minutes at 10,000 x g, 4 °C. We invert the filter and collect the purified origami in a fresh tube after centrifuging at 1000 x g for 5 minutes.

### Nanoparticle functionalization

We adapt a freeze and thaw approach from the work of Liu et. al.^43^ Typically we take 100 µl of 1 mg/ml 80 nm BioPure AgNPs from nanoComposix, USA, into a low-bind 1.5 ml Eppendorf tube. We reconstitute two tubes of lyophilized thiol-modified ssDNA (5’-thiol-20T-3’) at 4 nmol from Ella Biotech GmbH in 670 µl of nuclease-free water each. Gradually, we add the thiolated-DNA strands to the NPs while mixing with a pipette. Next, we introduce 60 µl of 5 M NaCl and mix gently. We then freeze the mixture at -20 °C for a minimum of 1 hour. For purification, we thaw the mixture and centrifuge at 2800 x g, 4 °C for 15 minutes. After discarding the supernatant, we add 1x BlueJuice loading dye (Merck, Germany) and pipette to mix. We load a 1.2 % agarose gel and perform electrophoresis at 100 V for 45 minutes to isolate the monomer functionalized NPs from aggregates and excess ssDNA. We cut and squeeze the band to get the NPs. We mix this with an equal volume of nuclease-free water and centrifuge again under same conditions. We remove the supernatant, check the concentration at Nanodrop 2000 (Thermo Fisher, USA) and store at 4 °C for later use.

### Surface preparation for immobilizing NanoAntennas with Biotinylated strands

We rinse 24 mm x 60 mm, 170 µm thick microscope coverslips (Carl Roth GmbH, Germany) with Milli-Q water and isopropanol, dry them with an air stream and treat them in a UV-Ozone cleaner (PSD-UV4, Novascan Technologies, USA) for 30 min at 100 °C. After applying SecureSeal™ Hybridization Chambers (2.6 mm depth, Grace BioLabs, USA) to the cleaned coverslips, we wash the chambers thrice with Phosophate-buffered saline (PBS) and incubate them for at least 5 minutes with BSA-biotin (0.5 mg/ml, Sigma Aldrich, USA), followed by 5 minutes with NeutrAvidin (0.2 mg/ml, Thermo Fisher Scientific, USA). We add 100 pM of Trident DNA origami (in 5 mM Tris, 2M NaCl, 1 mM EDTA) with twelve biotinylated ssDNA extensions at the base to bind to NeutrAvidin and wash off excess origami after one and a half minutes. Next, we add functionalized NPs at 0.05 OD (measured at a path length of 1 mm) and incubate overnight in the dark at room temperature in the buffer (5 mM Tris, 2M NaCl, 1 mM EDTA). The following day, we wash 4-5 times with the same buffer to remove unbound nanoparticles.

### Nucleic acid assay

We fold Trident DNA origami with 10 capture strands protruding from the hotspot to detect a 151 nt synthetic DNA sequence specific to the OXA-48 gene carrying an antibiotic resistance (see Supplementary Note 1.1). We assemble the NanoAntennas as mentioned above and incubate with 4 nM of the target strand with 12 nM of an Alexa Fluor 647-labelled imager strand (17 nt) and 12 nM of a blocker strand (10 nt) in a buffer containing 5 mM Tris, 2M NaCl, 1 mM EDTA, at 37 °C for 1 hour. After incubation, we wash off the unbound strands with the same buffer.

### Surface preparation for patterned NanoAntennas

We adapt and modify the DNA origami placement method by Shetty et. al.^35^ We rinse 25 mm x 75 mm, 170 µm thick microscope coverslips (Electron Microscopy Sciences, Pennsylvania, USA) with Milli-Q water and isopropanol, dry them with an air stream. We mark a rectangle of ∼7 mm x 5 mm with a marker in the center of one side and prepare samples on the reverse. We cover the unmarked area with extra glass coverslips and treat the marked region in an UV-Ozone cleaner for 30 min at 100 °C to make it hydrophilic. We centrifuge 350 µl of 400 nm polystyrene (PS) nanospheres (Thermo Scientific™ Nanosphere™ Size standard 3400A) for 5 minutes at 10,000 x g at room temperature, discard the supernatant, re-suspend in 350 µl of Milli-Q water, and repeat the centrifugation twice. We re-suspend the final pellet in 100 µl of 25 % ethanol/water and drop-cast 10 µl onto the marked area at a ∼30° angle. After heating at 60 °C for 4-5 minutes, we expose the coverslips to hexamethyldisilazane (HMDS) vapors for 30 minutes in a sealed glass chamber. We then ultrasonicate in water to lift-off spheres, dry with nitrogen stream and annealed at 120 °C for 5 minutes. We dilute the Triangle DNA origami to 150 pM, add ∼200 µl to the marked area and incubate for 1 hour at room temperature in placement buffer (PB, 40 mM Tris, 40 mM MgCl_2_ at pH∼8.4). After washing with placement buffer and Tween20, we add 350 pM Trident DNA origami and incubate for another 1 hour, followed by repeated washing. Finally, we add 50 µl of 0.2 OD (measured at a path length of 1 mm) functionalized 80 nm AgNPs in placement buffer with Tween and incubate overnight in the dark. We wash the coverslips 5-6 times in placement buffer containing Tween.

### Sample preparation for measurements on the reader

We wash and keep the patterned area moist, place an adhesive microfluidic chip (Straight channel chip with adhesive tape, Fluidic 268, channel width: 2500 µm, channel depth: 150 µm, channel length: 58.5 mm, material: Zeonor™, from microfluidic ChipShop GmbH, Germany) over the coverslip, and flush the channel with placement buffer containing Tween before further measurements.

## Acknowledgements

R.Y. thanks Dr. Lorena C. Manzanares, Dr. Alan Szalai, Julian Bauer and Dr. Ece Büber for fruitful discussions. P.T. gratefully acknowledges financial support from the DFG (TI 329/9-2 (project number 267681426), INST 86/1904-1 FUGG, INST 86/2224-1 FUGG, excellence cluster e-conversion EXC 2089/1-390776260), Sino-German Center for Research Promotion (grant agreement C-0008), BMBF (Grants POCEMON, 13N14336, and SIBOF, 03VP03891). K.T. acknowledges the support by Humboldt Research Fellowships from the Alexander von Humboldt Foundation. P.T. and M.D. acknowledge funding by the Federal Ministry of Education and Research (BMBF, 13N16929) and the Free State of Bavaria under the Excellence Strategy of the Federal Government and the Länder through the ONE MUNICH Project Munich Multiscale Biofabrication, and by the Center for NanoScience (CeNS).

## Author contributions

The manuscript was written through contributions of all authors. All authors have given approval to the final version of the manuscript.

## REFERENCES

1. Holzmeister, P., Acuna, G. P., Grohmann, D. & Tinnefeld, P. Breaking the concentration limit of optical single-molecule detection. Chem. Soc. Rev. 43, 1014–1028 (2014).

2. Walt, D. R. Optical Methods for Single Molecule Detection and Analysis. Anal Chem 85, 1258–1263 (2013).

3. Bizuayehu, T. T. et al. Long-read single-molecule RNA structure sequencing using nanopore. Nucleic Acids Res 50, e120–e120 (2022).

4. Gooding, J. J. & Gaus, K. Single-Molecule Sensors: Challenges and Opportunities for Quantitative Analysis. Angewandte Chemie International Edition 55, 11354–11366 (2016).

5. Howorka, S. & Hesse, J. Microarrays and single molecules: an exciting combination. Soft Matter 10, 931 (2014).

6. Mayr, R. et al. A microfluidic platform for transcription- and amplification-free detection of zepto-mole amounts of nucleic acid molecules. Biosens Bioelectron 78, 1–6 (2016).

7. Punj, D. et al. A plasmonic ‘antenna-in-box’ platform for enhanced single-molecule analysis at micromolar concentrations. Nat Nanotechnol 8, 512–516 (2013).

8. Anger, P., Bharadwaj, P. & Novotny, L. Enhancement and Quenching of Single-Molecule Fluorescence. Phys Rev Lett 96, 113002 (2006).

9. Curto, A. G. et al. Unidirectional Emission of a Quantum Dot Coupled to a Nanoantenna. Science (1979) 329, 930–933 (2010).

10. Rothemund, P. W. K. Folding DNA to create nanoscale shapes and patterns. Nature 440, 297–302 (2006).

11. Acuna, G. P. et al. Fluorescence Enhancement at Docking Sites of DNA-Directed Self-Assembled Nanoantennas. Science (1979) 338, 506–510 (2012).

12. Glembockyte, V., Grabenhorst, L., Trofymchuk, K. & Tinnefeld, P. DNA Origami Nanoantennas for Fluorescence Enhancement. Acc Chem Res 54, 3338–3348 (2021).

13. Trofymchuk, K. et al. Addressable nanoantennas with cleared hotspots for single-molecule detection on a portable smartphone microscope. Nat Commun 12, 950 (2021).

14. Kelley, S. O. What Are Clinically Relevant Levels of Cellular and Biomolecular Analytes? ACS Sens 2, 193–197 (2017).

15. Punj, D. et al. Plasmonic antennas and zero-mode waveguides to enhance single molecule fluorescence detection and fluorescence correlation spectroscopy toward physiological concentrations. WIREs Nanomedicine and Nanobiotechnology 6, 268–282 (2014).

16. Peng, S., Wang, W. & Chen, C. Breaking the Concentration Barrier for Single-Molecule Fluorescence Measurements. Chemistry – A European Journal 24, 1002–1009 (2018).

17. White, D. S., Smith, M. A., Chanda, B. & Goldsmith, R. H. Strategies for Overcoming the Single-Molecule Concentration Barrier. ACS Measurement Science Au 3, 239–257 (2023).

18. Rissin, D. M. et al. Single-molecule enzyme-linked immunosorbent assay detects serum proteins at subfemtomolar concentrations. Nat Biotechnol 28, 595–599 (2010).

19. Bell, N. A. W. & Keyser, U. F. Digitally encoded DNA nanostructures for multiplexed, single-molecule protein sensing with nanopores. Nat Nanotechnol 11, 645–651 (2016).

20. Hindson, B. J. et al. High-Throughput Droplet Digital PCR System for Absolute Quantitation of DNA Copy Number. Anal Chem 83, 8604–8610 (2011).

21. Loretan, M. et al. Direct single-molecule detection and super-resolution imaging with a low-cost portable smartphone-based microscope. bioRxiv 2024.05.08.593103 (2024) doi:10.1101/2024.05.08.593103.

22. Brown, J. W. P. et al. Single-molecule detection on a portable 3D-printed microscope. Nat Commun 10, 5662 (2019).

23. Moya Muñoz, G. G. et al. Single-molecule detection and super-resolution imaging with a portable and adaptable 3D-printed microscopy platform (Brick-MIC). Sci Adv 10, (2024).

24. Wu, Y., Tilley, R. D. & Gooding, J. J. Challenges and Solutions in Developing Ultrasensitive Biosensors. J Am Chem Soc 141, 1162–1170 (2019).

25. Wanunu, M. et al. Rapid electronic detection of probe-specific microRNAs using thin nanopore sensors. Nat Nanotechnol 5, 807–814 (2010).

26. Giljohann, D. A. & Mirkin, C. A. Drivers of biodiagnostic development. Nature 462, 461–464 (2009).

27. Yan, L. et al. Isothermal amplified detection of DNA and RNA. Mol Biosyst 10, 970 (2014).

28. Kaminski, M. M., Abudayyeh, O. O., Gootenberg, J. S., Zhang, F. & Collins, J. J. CRISPR-based diagnostics. Nat Biomed Eng 5, 643–656 (2021).

29. Mirabile, A. et al. Advancing Pathogen Identification: The Role of Digital PCR in Enhancing Diagnostic Power in Different Settings. Diagnostics 14, 1598 (2024).

30. Murray, C. J. L. et al. Global burden of bacterial antimicrobial resistance in 2019: a systematic analysis. The Lancet 399, 629–655 (2022).

31. Close, C. et al. Maximizing the Accessibility in DNA Origami Nanoantenna Plasmonic Hotspots. Adv Mater Interfaces 9, (2022).

32. Poirel, L., Bonnin, R. A. & Nordmann, P. Genetic Features of the Widespread Plasmid Coding for the Carbapenemase OXA-48. Antimicrob Agents Chemother 56, 559–562 (2012).

33. Budia-Silva, M. et al. International and regional spread of carbapenem-resistant Klebsiella pneumoniae in Europe. Nat Commun 15, 5092 (2024).

34. Leake, M. C. The physics of life: one molecule at a time. Philosophical Transactions of the Royal Society B: Biological Sciences 368, 20120248 (2013).

35. Shetty, R. M., Brady, S. R., Rothemund, P. W. K., Hariadi, R. F. & Gopinath, A. Bench-Top Fabrication of Single-Molecule Nanoarrays by DNA Origami Placement. ACS Nano 15, 11441–11450 (2021).

36. Martynenko, I. V. et al. Site-directed placement of three-dimensional DNA origami. Nat Nanotechnol 18, 1456–1462 (2023).

37. Dass, M. R. L. ; P. C. ; S. C. ; H. L. ; T. J. ; M. I. V. ; R. U. ; P. G. ; L. T. Self-assembled physical unclonable function labels based on plasmonic coupling. ArXiv (2023).

38. Vogel, N., Weiss, C. K. & Landfester, K. From soft to hard: the generation of functional and complex colloidal monolayers for nanolithography. Soft Matter 8, 4044–4061 (2012).

39. Gopinath, A. & Rothemund, P. W. K. Optimized Assembly and Covalent Coupling of Single-Molecule DNA Origami Nanoarrays. ACS Nano 8, 12030–12040 (2014).

40. Jungmann, R. et al. Single-Molecule Kinetics and Super-Resolution Microscopy by Fluorescence Imaging of Transient Binding on DNA Origami. Nano Lett 10, 4756–4761 (2010).

41. Sharonov, A. & Hochstrasser, R. M. Wide-field subdiffraction imaging by accumulated binding of diffusing probes. Proceedings of the National Academy of Sciences 103, 18911–18916 (2006).

42. Li, S., He, J. & Xu, Q.-H. Aggregation of Metal-Nanoparticle-Induced Fluorescence Enhancement and Its Application in Sensing. ACS Omega 5, 41–48 (2020).

43. Liu, B. & Liu, J. Freezing Directed Construction of Bio/Nano Interfaces: Reagentless Conjugation, Denser Spherical Nucleic Acids, and Better Nanoflares. J Am Chem Soc 139, 9471–9474 (2017).

44. Zang, F. et al. Ultrasensitive Ebola Virus Antigen Sensing via 3D Nanoantenna Arrays. Advanced Materials 31, (2019).

45. Liu, X. et al. Complex silica composite nanomaterials templated with DNA origami. Nature 559, 593–598 (2018).

46. Nguyen, L., Döblinger, M., Liedl, T. & Heuer-Jungemann, A. DNA-Origami-Templated Silica Growth by Sol–Gel Chemistry. Angewandte Chemie International Edition 58, 912–916 (2019).

47. Wassermann, L. M., Scheckenbach, M., Baptist, A. V., Glembockyte, V. & Heuer-Jungemann, A. Full Site-Specific Addressability in DNA Origami-Templated Silica Nanostructures. Advanced Materials 35, (2023).

48. Jing, X. et al. Solidifying framework nucleic acids with silica. Nat Protoc 14, 2416–2436 (2019).

49. Chandrasekaran, A. R. Nuclease resistance of DNA nanostructures. Nat Rev Chem 5, 225–239 (2021).

50. Yurke, B., Turberfield, A. J., Mills, A. P., Simmel, F. C. & Neumann, J. L. A DNA-fuelled molecular machine made of DNA. Nature 406, 605–608 (2000).

51. Douglas, S. M. et al. Rapid prototyping of 3D DNA-origami shapes with caDNAno. Nucleic Acids Res 37, 5001–5006 (2009).

52. Wagenbauer, K. F. et al. How We Make DNA Origami_supplement. ChemBioChem 18, 1873–1885 (2017).

