## Supplementary information for "Bringing Attomolar Detection to the Point-of-Care with Nanopatterned DNA Origami Nanoantennas"

for

**Figure S1. Trident DNA Origami design**

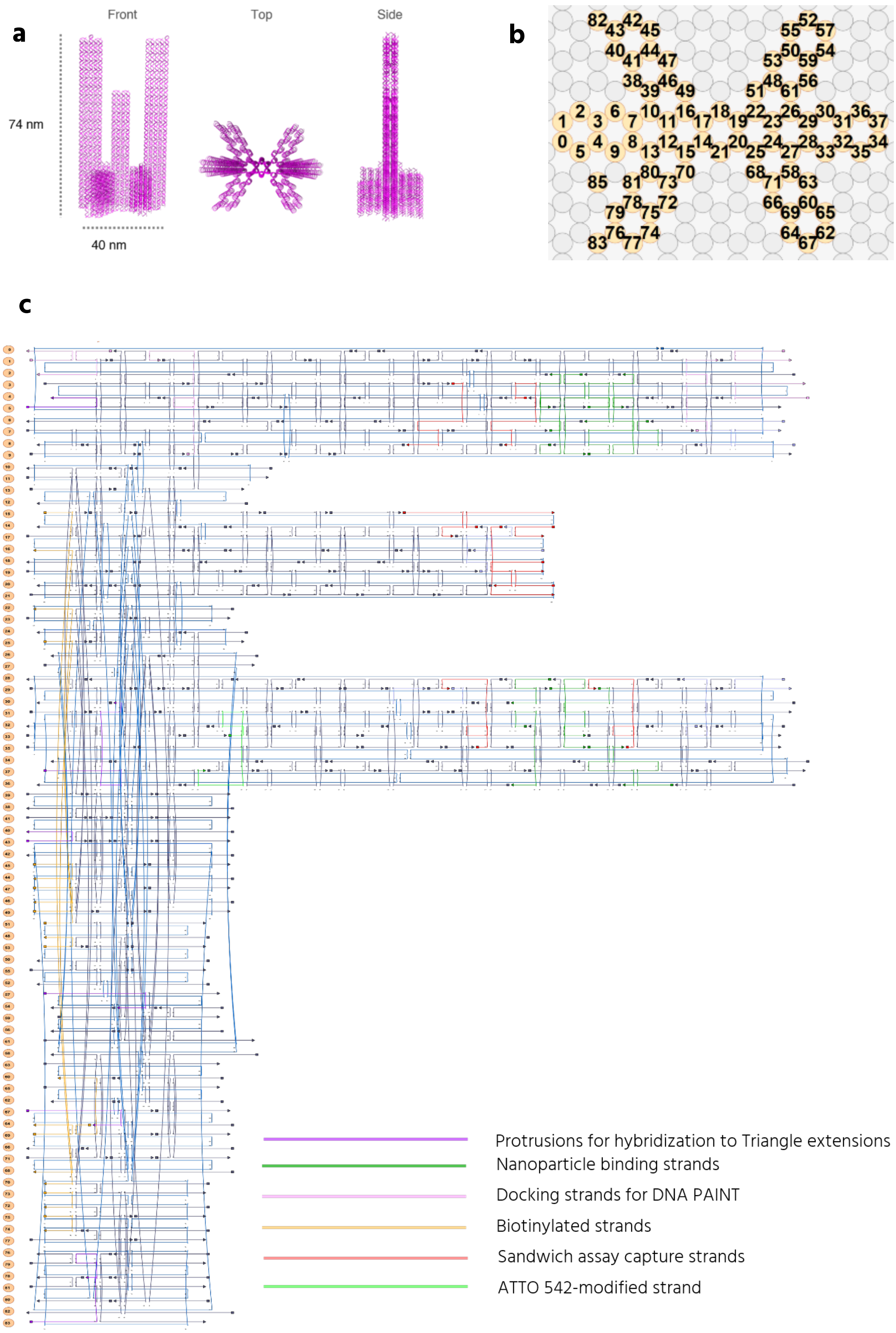

Figure S1. **a** 3D model of the Trident DNA origami. **b** Helical design snapshot from cadnano2[1]. **c** Scaffold routing and staple layout for Trident DNA origami. Scaffold in blue, biotin modified staples in orange, capture strands in red, nanoparticle binding strands in green, ATTO 542 labelled strand in neon green, PAINT staples in light pink, protrusions for hybridization to triangle in purple.

#### Figure S2. Blocker strand.

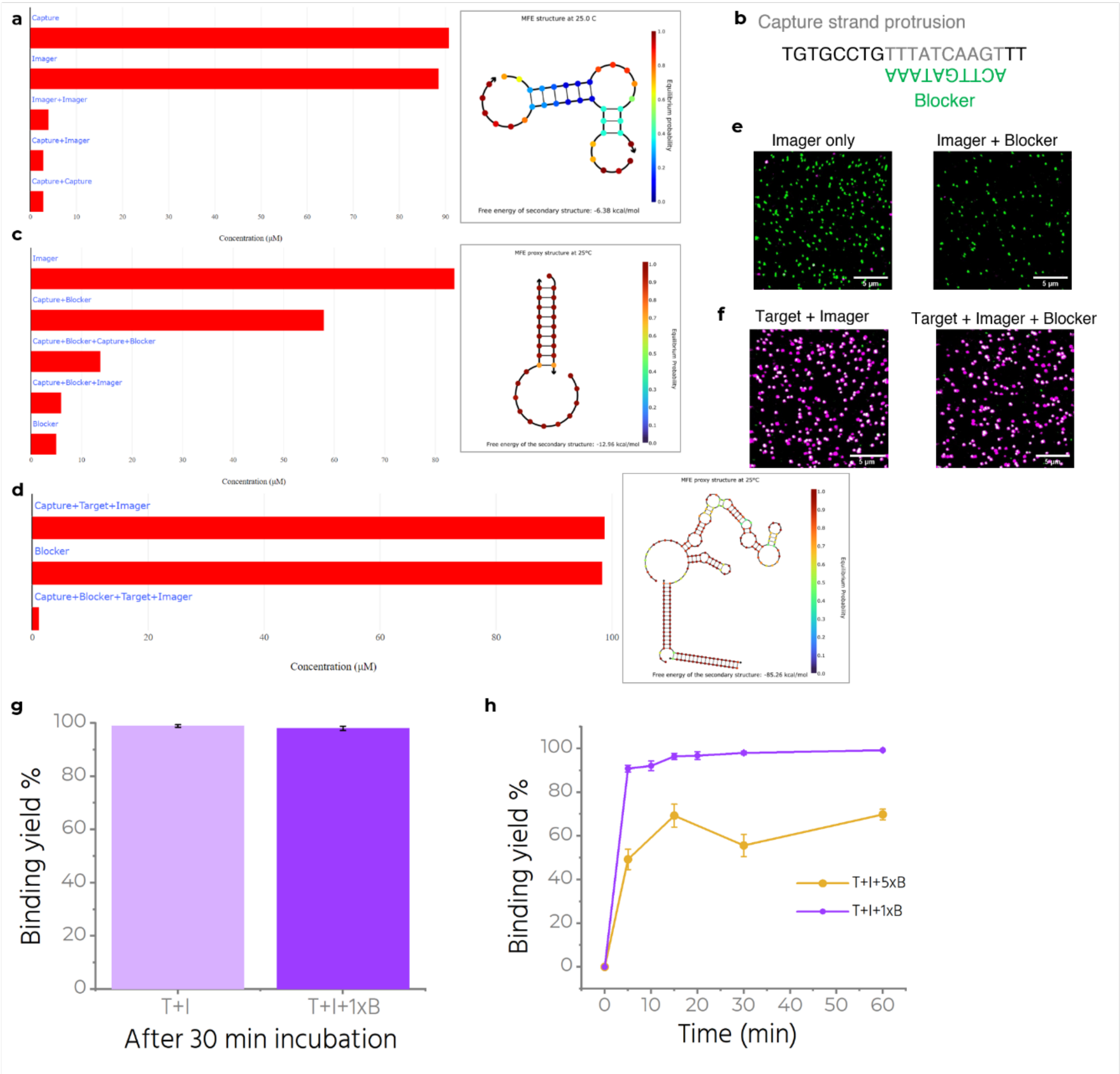


**Figure S2.** NUPACK analysis: **a** Interaction probability histogram and equilibrium probability chart for complexes formed in a solution with capture and imager strand. **b** Designed sequence for blocker strand to restrict interaction between capture and imager strand. **c** Interaction probability histogram for complexes formed in a solution with capture, imager and blocker strand. The equilibrium probability chart shows stable interaction between capture and blocker. **d** Interaction probability histogram for complexes formed in a solution with capture, imager, blocker and target strand, with equilibrium probability chart showing a stable interaction between capture, target and imager. **e** Confocal fluorescence scans for NanoAntennas incubated with imager only and with imager+blocker. **f** Confocal fluorescence scans for NanoAntennas incubated with target+imager and with target+imager+blocker. **g** Comparing binding yield of two samples after 30 minutes of incubation with 4nM T + 12 nM Imager and 4nM T + 12 nM Imager + 12 nM Blocker. **h** Effect of Blocker concentration on binding kinetics of the assay. We use 1x Blocker (12 nM) throughout the manuscript.

**Figure S3. Trident NanoAntenna characterization.**

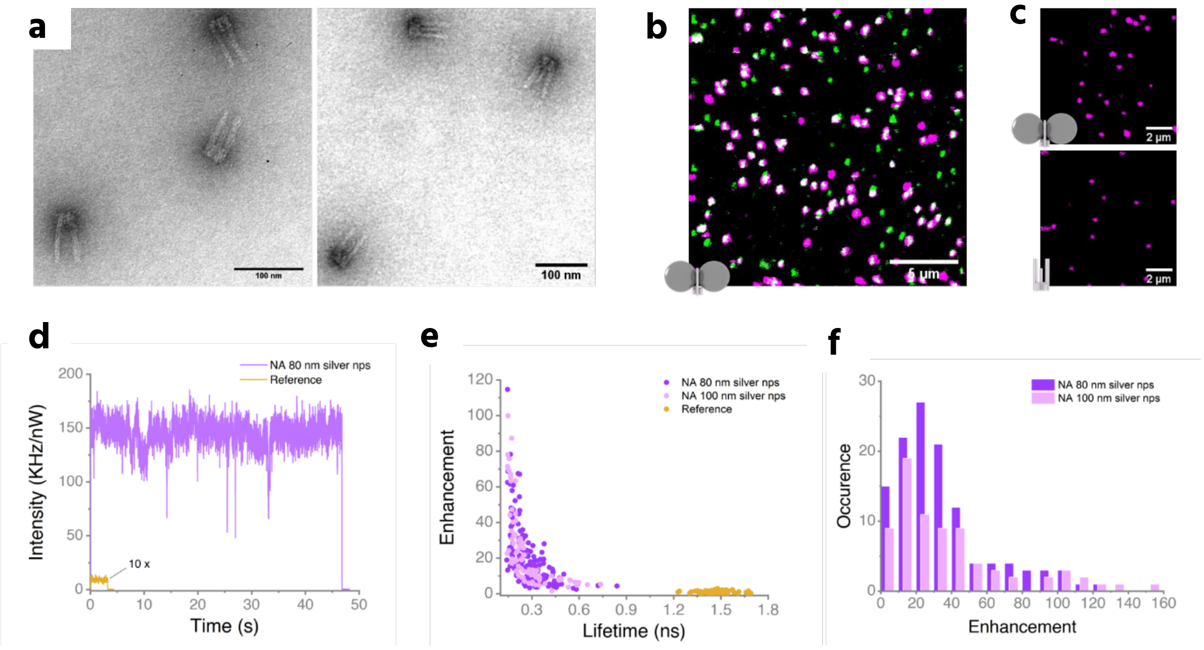


**Figure S3. a** TEM images showing successfully folded Trident. **b** Exemplary confocal fluorescence scan for antennas with ATTO 542 (green spots) and co-localized spots in white showing the presence of both ATTO542 and Alexa Fluor 647 dye on the same origami. **c** 10x10 µm exemplary confocal fluorescence scan for Trident with nanoparticles on the top and Trident without nanoparticles on the bottom, measured at 50 nW and 500 nW red laser power respectively. **d** Exemplary transient comparing intensity from a single Alexa F 647 bound to the Trident in presence (NanoAntenna (NA)) and absence of nanoparticles (reference). The reference intensity has been multiplied ten times for better visibility. **e** Enhancement and lifetime correlation for NA with 80 nm silver nanoparticles, with 100 nm silver nanoparticles and without nanoparticles. **f** Enhancement distribution in case of NA with 80 nm silver nanoparticles and 100 nm silver NPs.

**Figure S4. Increased capture strands.**

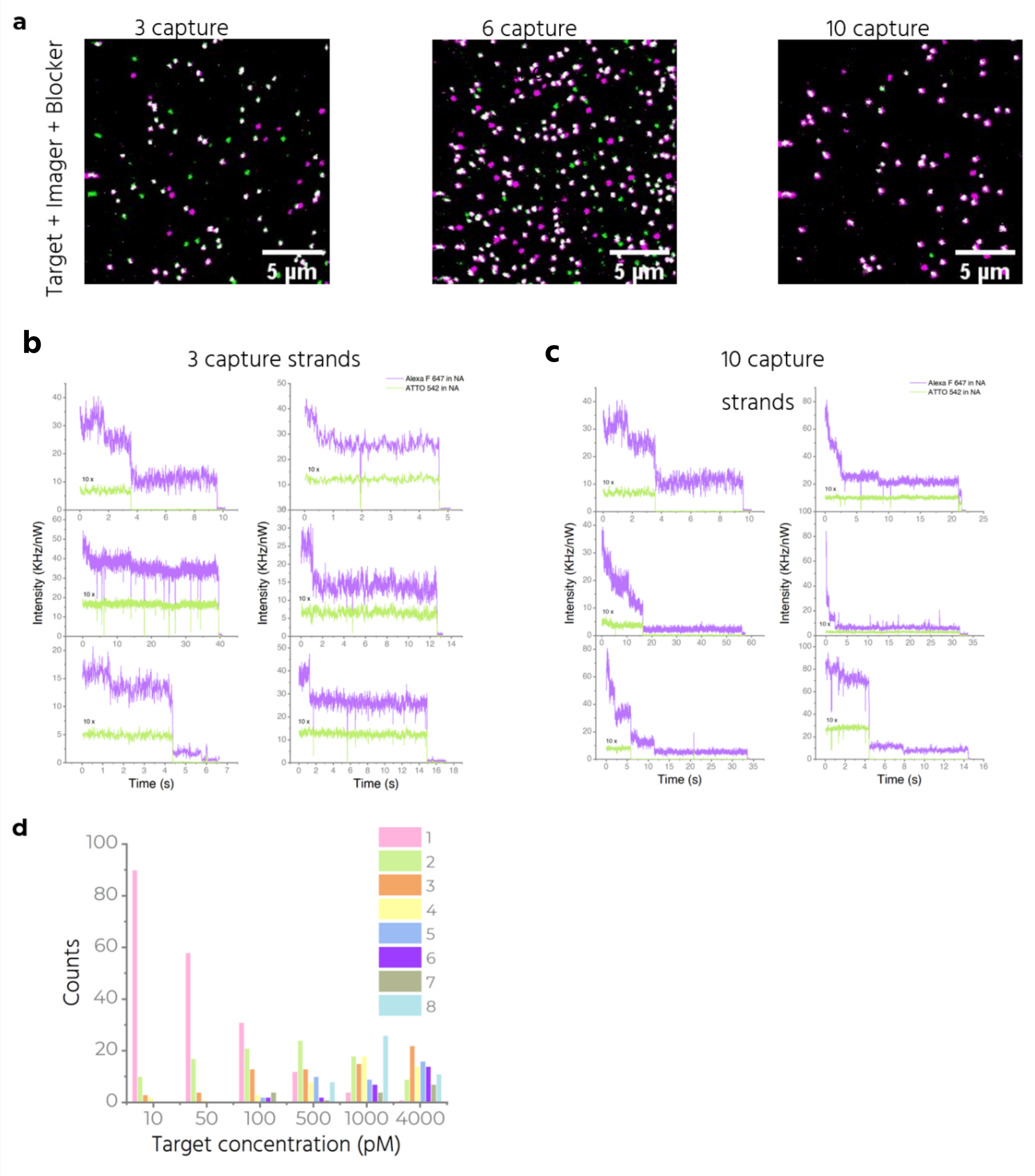


Figure S4. **a** Confocal fluorescence scans recorded after the assay comparing NanoAntennas with 3, 6 and 10 capture strands. **b** Exemplary transients for NA with 3 capture strands. A single step for ATTO 542 (green) represents a single Trident and 1-3 bleaching steps for Alexa Fluor 647 (in purple) represents the number of target strands captured. ATTO 542 is excited at 500 nW and the intensity is multiplied 10 times for better visibility. Alexa F 647 is excited with 50 nW. **c** Exemplary transients for NA with 10 capture strands. **d** Bleaching step analysis for NA with 10 capture strands at different concentrations showing up to 8 bleaching steps recorded at higher concentrations. ATTO 542 is excited at 500 nW and the intensity is multiplied 10 times for better visibility. Alexa F 647 is excited with 50 nW.

**Figure S5. Effect of increased number of capture strands on the fluorescence enhancement.**

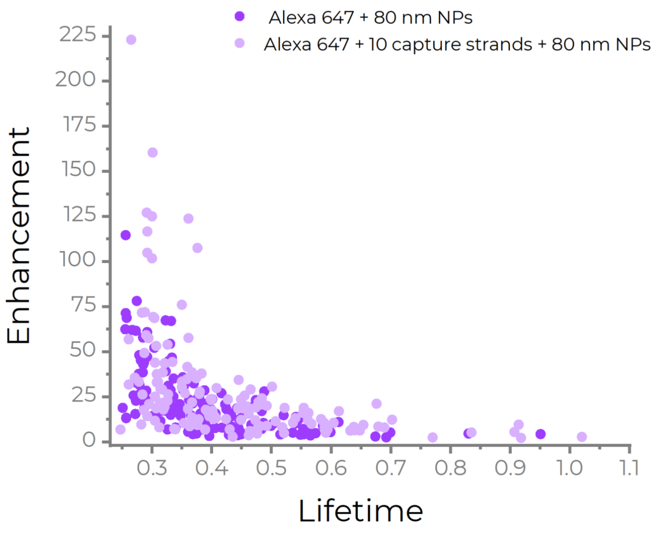


**Figure S5.** Two different Trident are folded, one with a fixed dye in the hotspot and no capture strands, and the other with a fixed dye in the hotspot and additionally 10 capture strands in the hotspot. For each type, NanoAntennas are assembled and confocal fluorescence scans are recorded to analyze and compare the effect of incorporating many capture strands on the binding of nanoparticles. Enhancement vs lifetime is plotted for each case, suggesting a similar trend for both samples.

**Figure S6. Example scans**

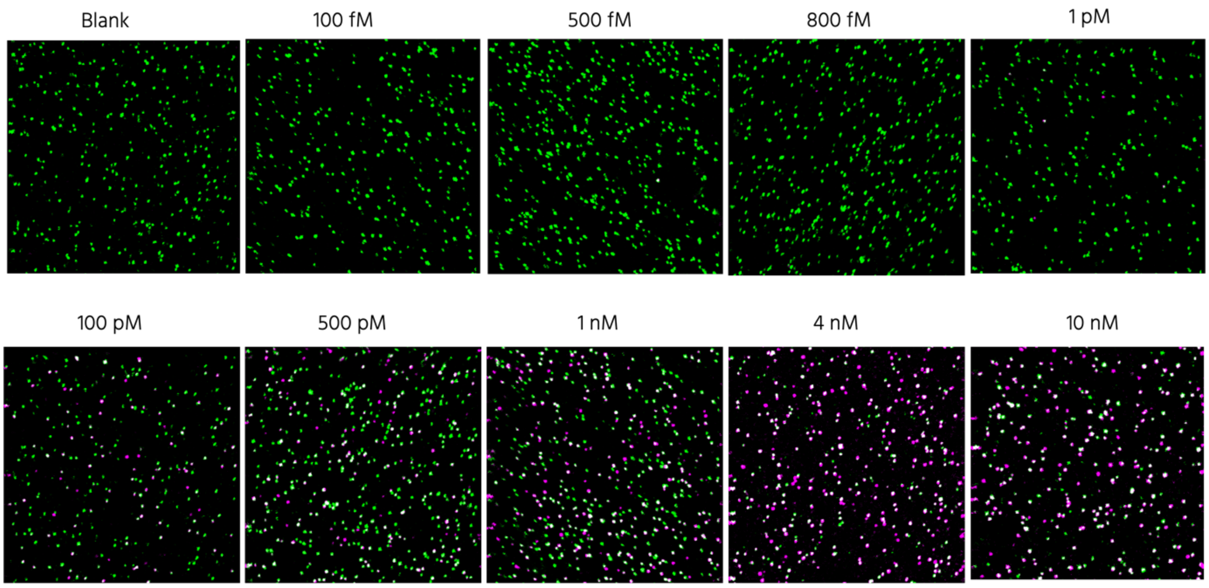


**Figure S6.** Exemplary confocal scans after the assay at different concentrations starting from absence of target (Top, left), to increase in target concentration from 100 fM up to 10 nM. Read from left to right, top to bottom. All samples had imager and blocker at 12 nM during the incubation of 1 hour at 37°C. Green spots represent the base dye on the Trident nanonatenna, and magenta represents the imager labelled with Alexa F 647.

##### Figure S7. Triangle DNA origami design

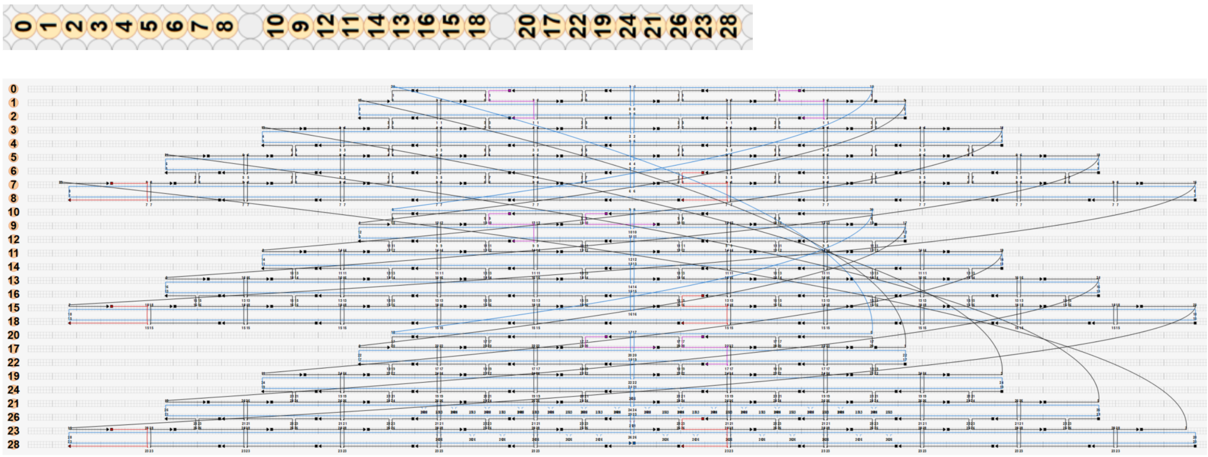


Figure S7. Helical design snapshot of the Triangle DNA origami in cadnano2 (above) and scaffold routing (below), showing DNA-PAINT docking sites in red and staples for capturing Trident in pink.

#### Figure S8. DNA-PAINT images of Trident directly bound to the binding site

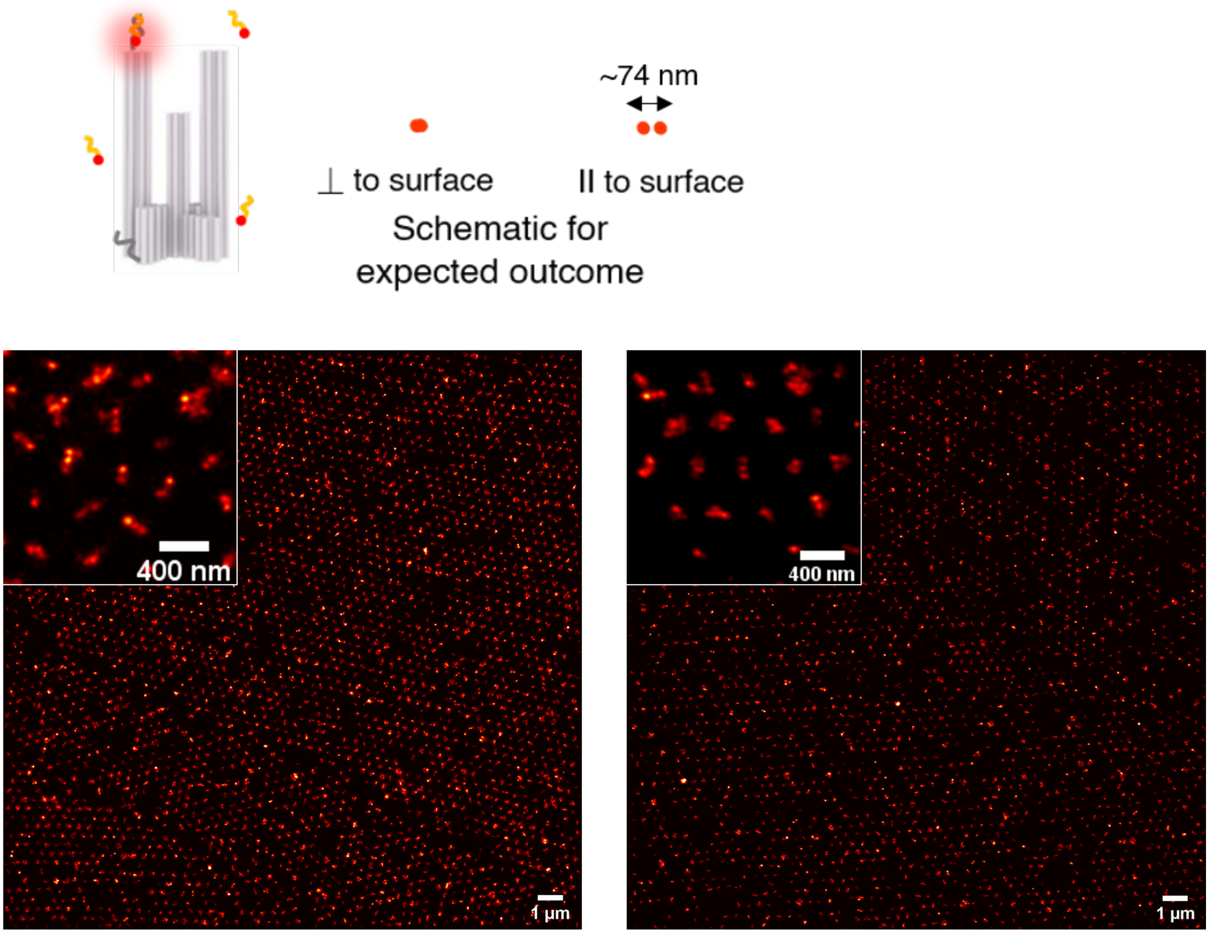


**Figure S8.** Trident is modified to include 3 protruding docking strands from the top of the left pillar-complex and 3 from the bottom. 3 strands per docking site allows to increase the chances of at least one of the strands being incorporated in most structures. (See sketch above. Only one strand at top and one at bottom in grey are shown for simplicity). A 6-nt or an 8-nt imager strand (orange) labelled with ATTO 655 (red sphere), having a sequence complementary to the docking strands transiently hybridizes to the docking sites. Parameters such as number of frames, exposure time of each frame, laser intensity, imager concentration, are optimized to achieve one hybridization event on each immobilized Trident at a time. Over 10,000 frames, the localization data is collected and analyzed using Picasso Localize and Picasso Render to achieve a super-resolved image as shown here. This design of docking sites allows us to differentiate between an upright orientation of the Trident and other random orientations. In these examples, the Trident is added directly on the coverslip with patterned hydrophilic binding sites. This causes Trident to get placed in random numbers and orientations as the binding sites have a much bigger surface area to cover. The Trident interacts with the binding sites only (no binding on the hydrophobic background) but we don't see defined orientations in this case.

**Figure S9. AFM image of Triangles placed on the binding sites**

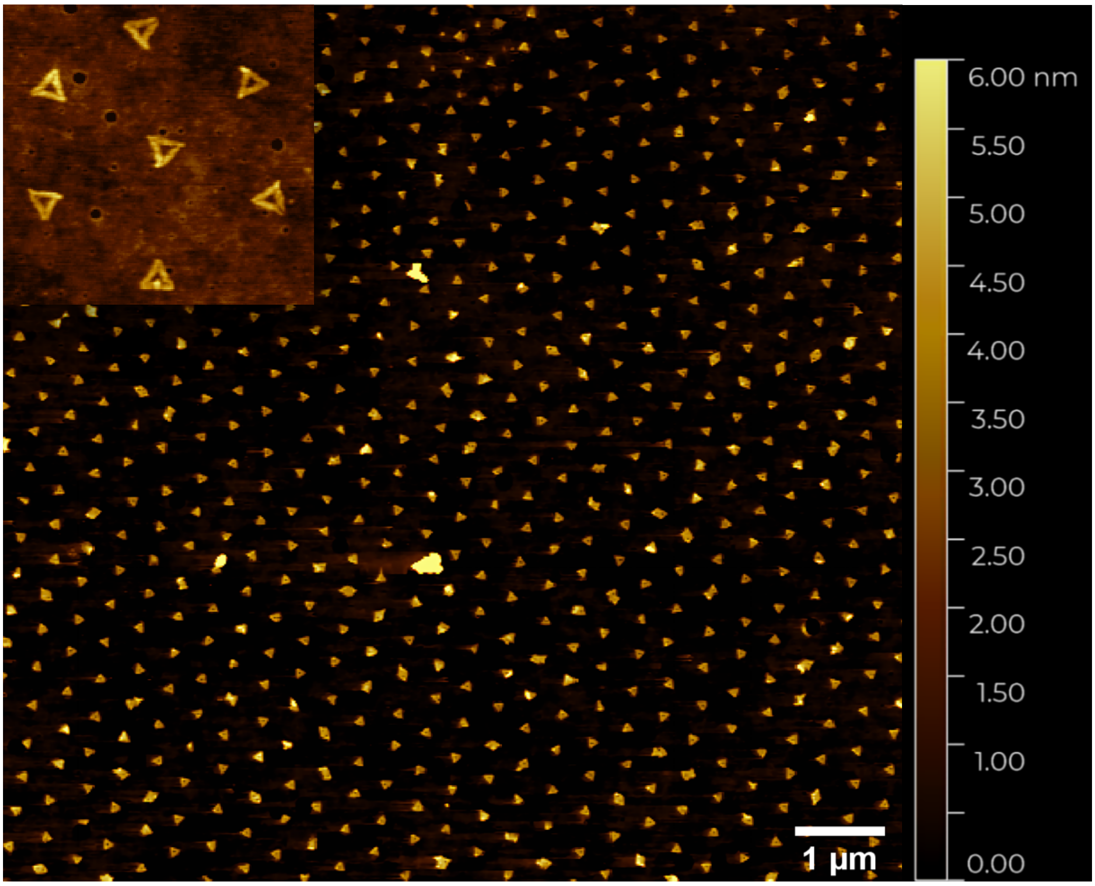


Figure S9. An exemplary AFM image after Triangle placement on the binding sites.

**Figure S10. DNA-PAINT images of Triangle placed on the binding sites**

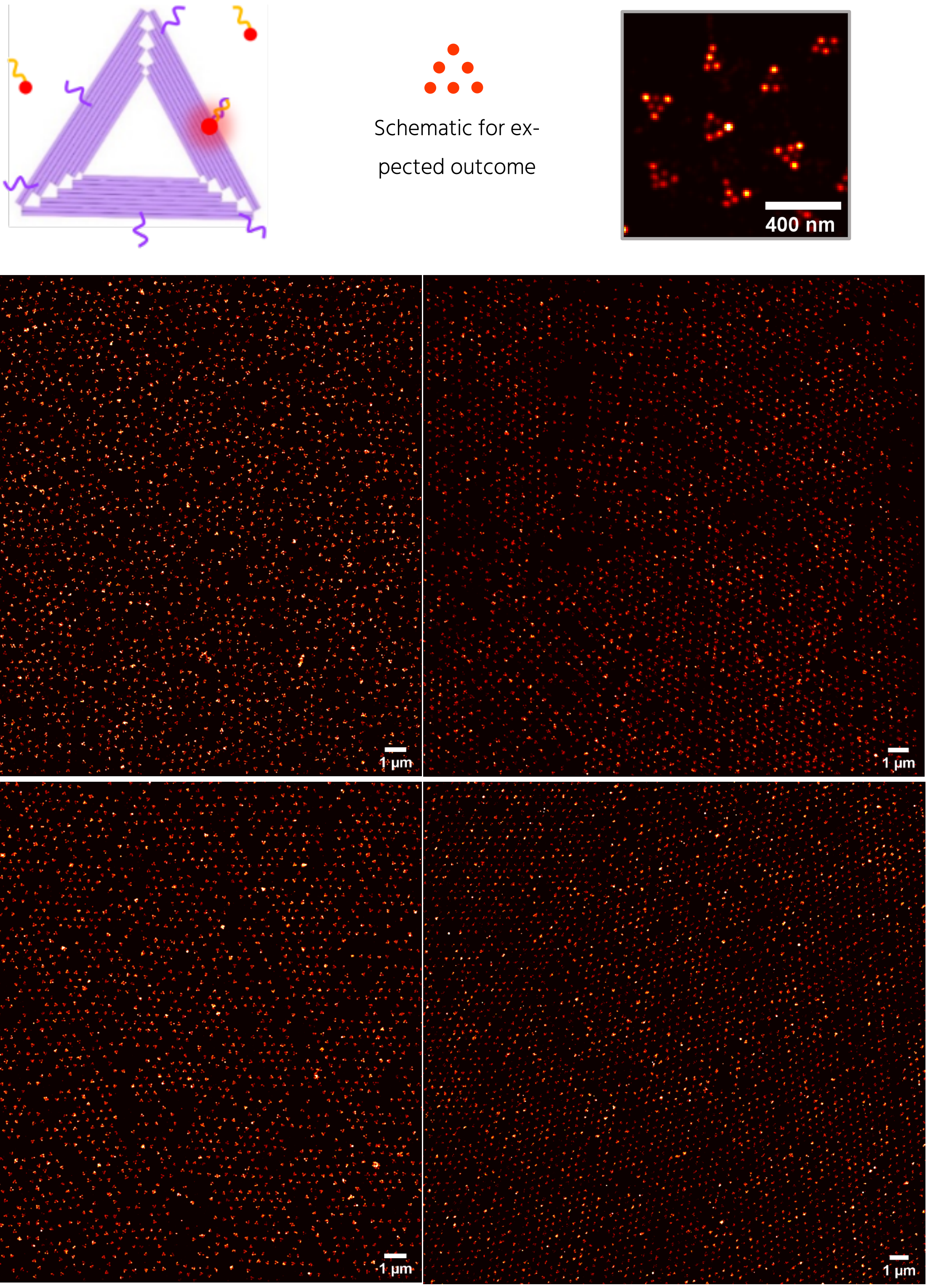

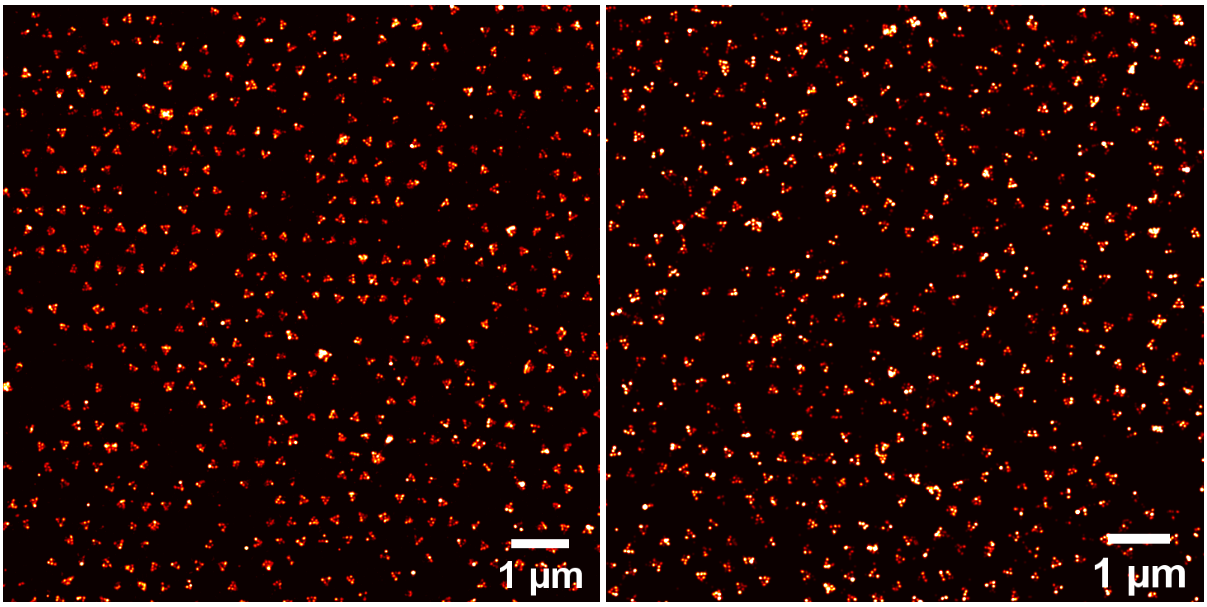


**Figure S10.** Sketch showing the design of 6 docking strands (purple) in a Triangle shape. 8-nt imager shown in orange was used labelled with ATTO 655 (red sphere) for the measurements. The expected outcome is shown on the top-middle and a zoom-in of a super-resolved image with the Triangle-shaped pattern shown on the top-right. Other example images of Triangles placed on the binding sites specifically.

**Figure S11. SEM images of polystyrene spheres assembled on a glass substrate**

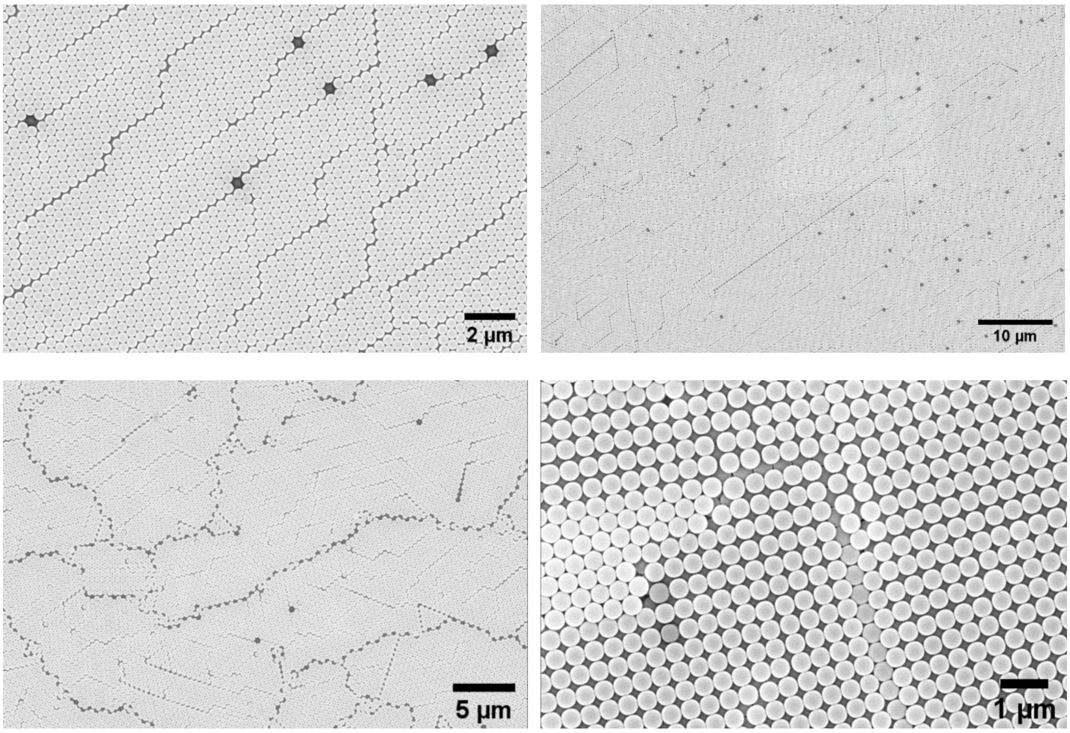


Figure S11. SEM images of polystyrene spheres assembled on a glass substrate showing mono- and muti-layer hexagonal assembly. The images reveal point and line defects in the assembly. In a smaller percentage, we also observe areas with a square lattice (last image).

**Figure S12. DNA-PAINT images for Trident placed on a nanopatterned surface and for Trident immobilized on a BSA-biotin-NeutrAvidin surface**

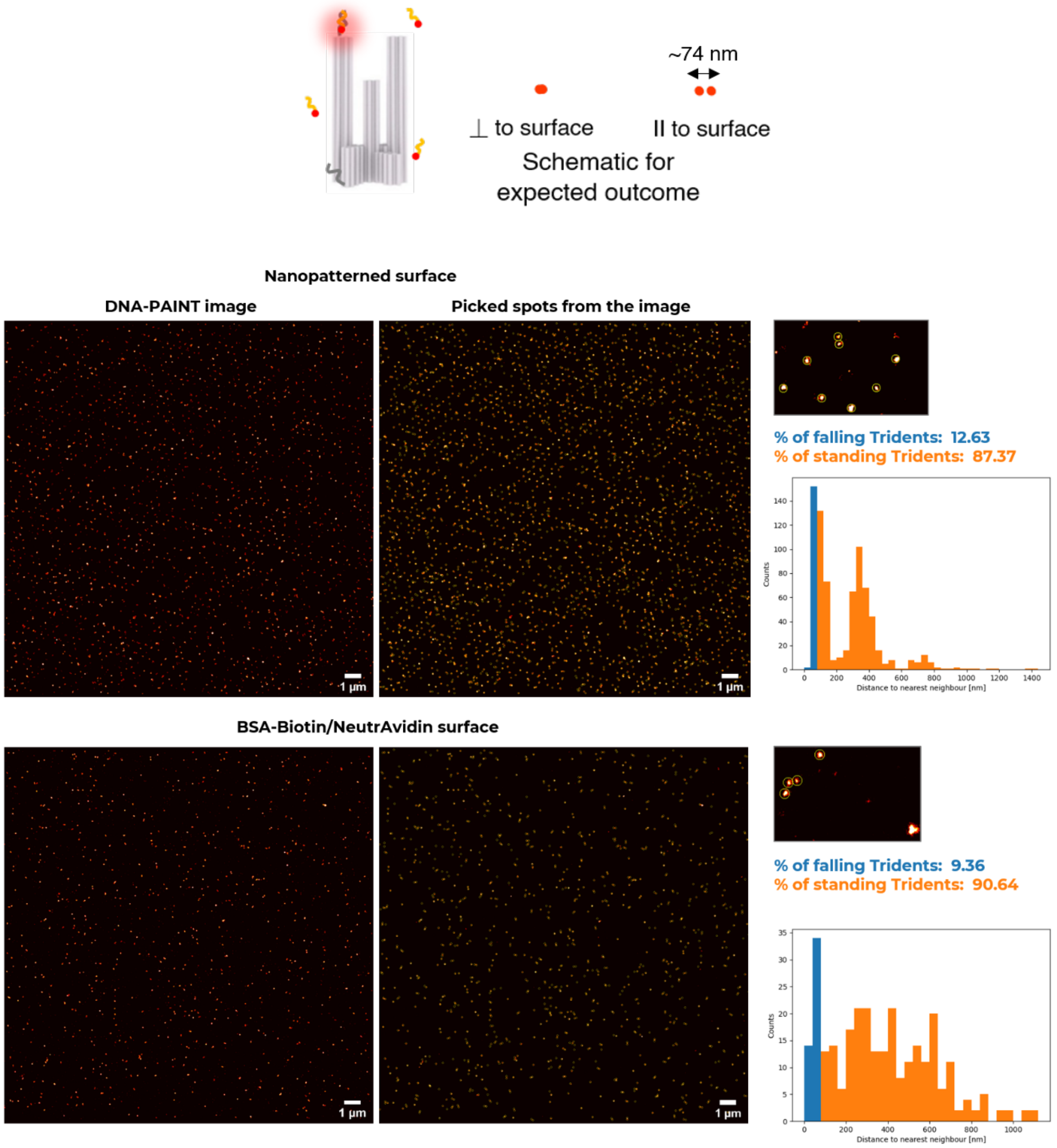


**Figure S12.** DNA-PAINT images are acquired and the % of upright orientation of the Trident is compared for NanoAntennas on a patterned surface vs NanoAntennas immobilized via biotinylated DNA. For the patterned surface, the Trident is bound to the Triangle via DNA hybridization. Individual spots are selected in Picasso Render and the localization information is used to plot the distance between each of those spots. An in-house python script allows segregating the data into '% of falling Tridents' and '% of standing Tridents'. According to the design of the docking sites, one spot is expected when the Trident is standing upright and two spots of varying (tilting and falling) distance of ~74 nm is expected when the orientation is not upright. To make it simpler, we assign any distance between two spots below 85 nm as falling (in blue) and any distance above that qualifies as standing (in orange) and plotted a histogram for the distance to nearest neighbor in each case.

**Figure S13. TEM images comparing the two methods for NP functionalization**

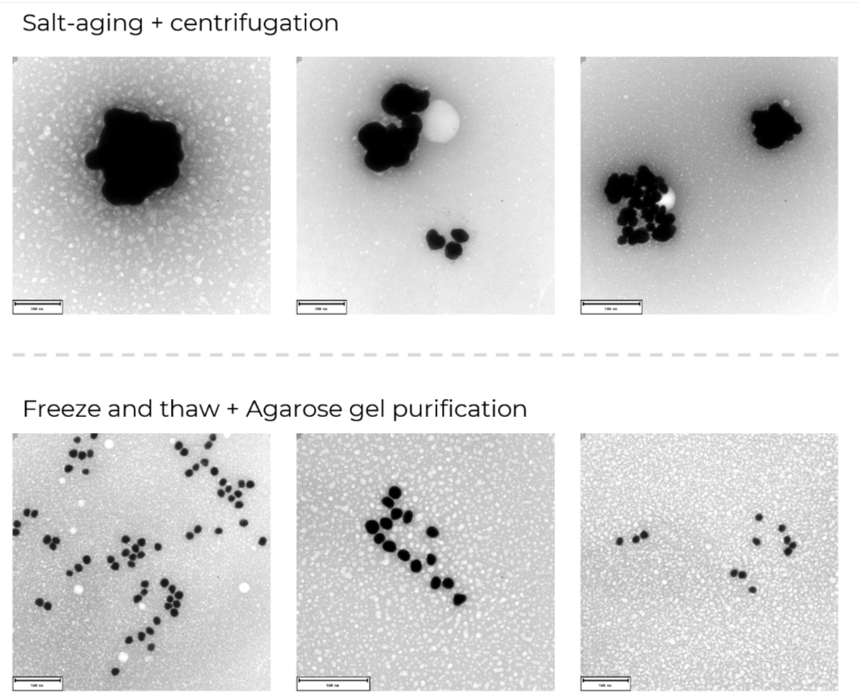


Figure S13. TEM images comparing two methods of nanoparticle functionalization and purification. We observe more aggregated chunks when we use the combination of salt-ageing followed by centrifugation, to functionalize and purify. NPs look less aggregated after freeze and thaw and purification with agarose gel electrophoresis. We use freeze and thaw followed by gel purification for this article.

**Figure S14. Example SEM images of Nanopatterned NanoAntennas**

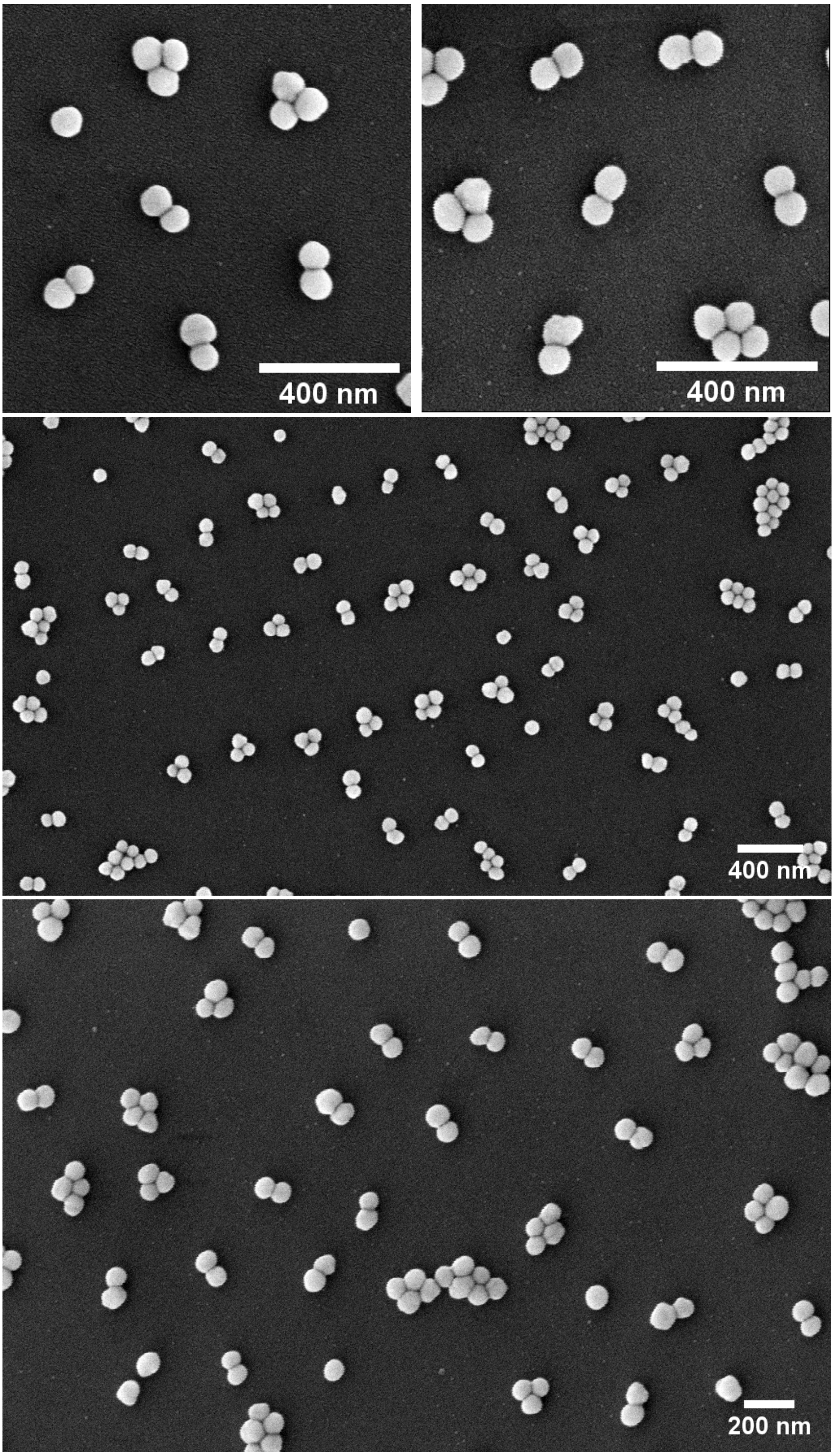

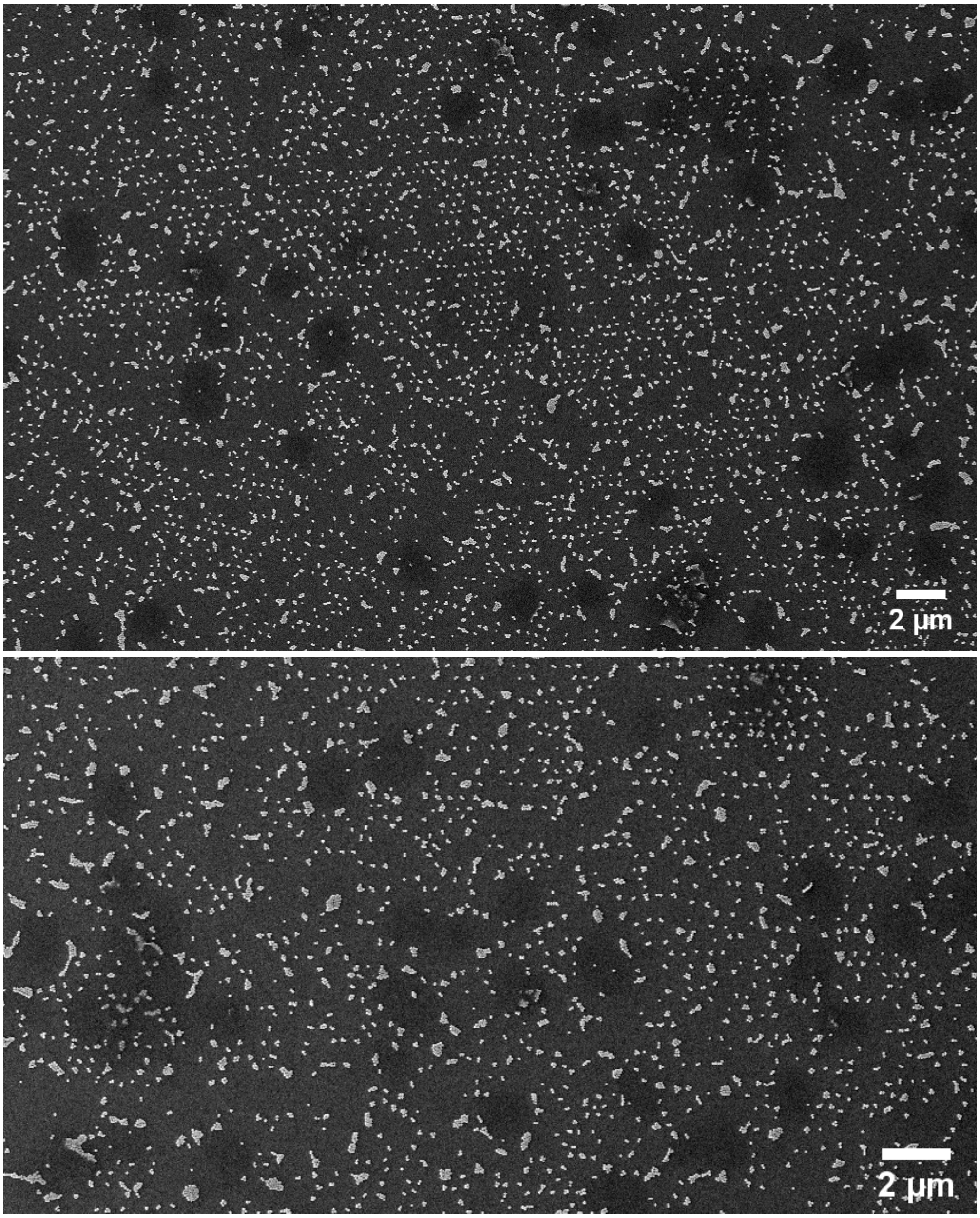


**Figure S14.** Example SEM images from three different coverslips with patterned NanoAntennas.

**Figure S15. SEM images after NP incubation with only Triangle placed on the binding site**

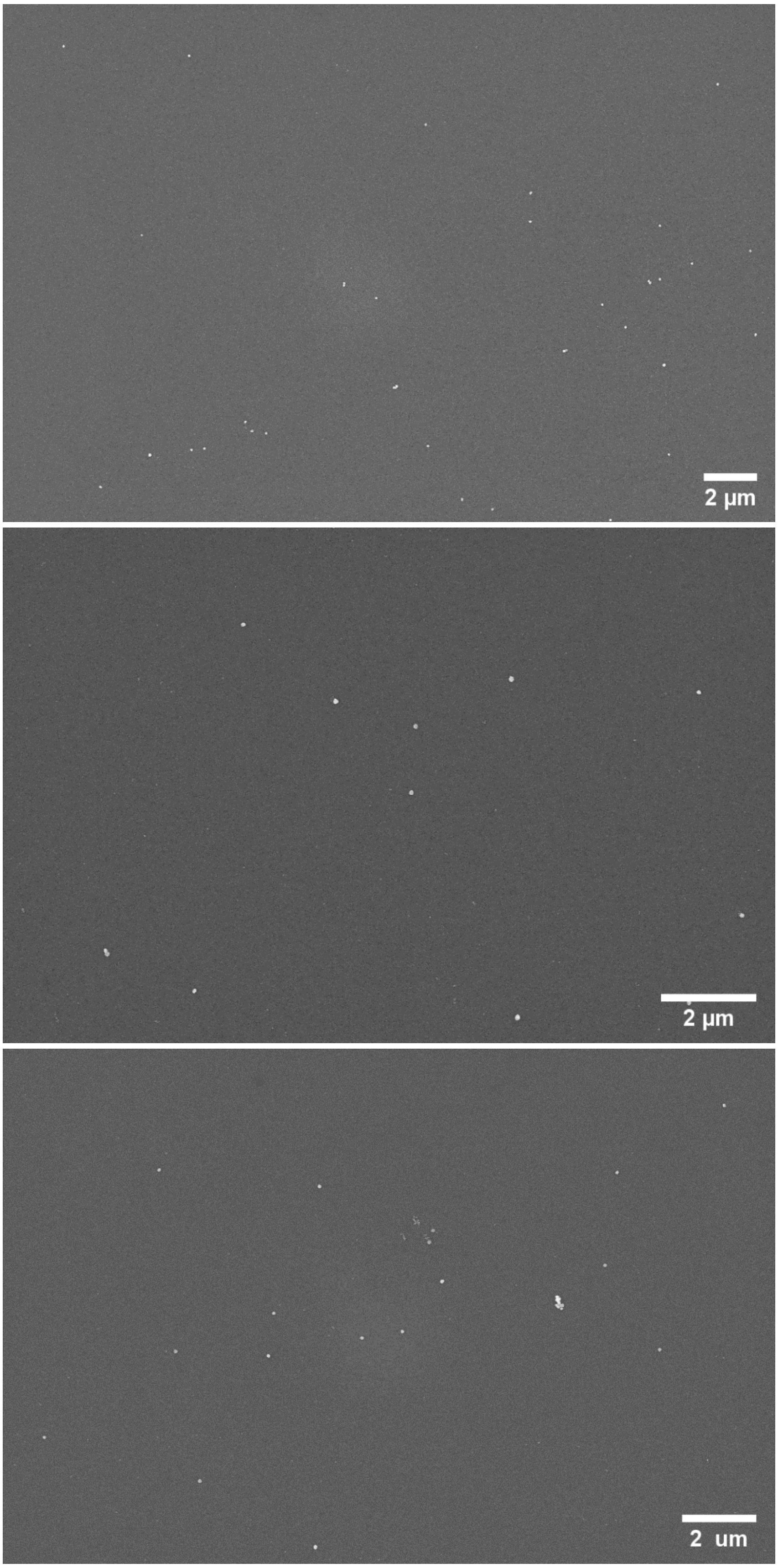

**Figure S15.** As a control, NPs are added to a patterned surface with Triangle placed on the binding sites and incubated overnight, washed and imaged with SEM. We observe no or minimal binding of NPs.

**Figure S16. SEM images after NP incubation with Trident placed directly on the binding site.**

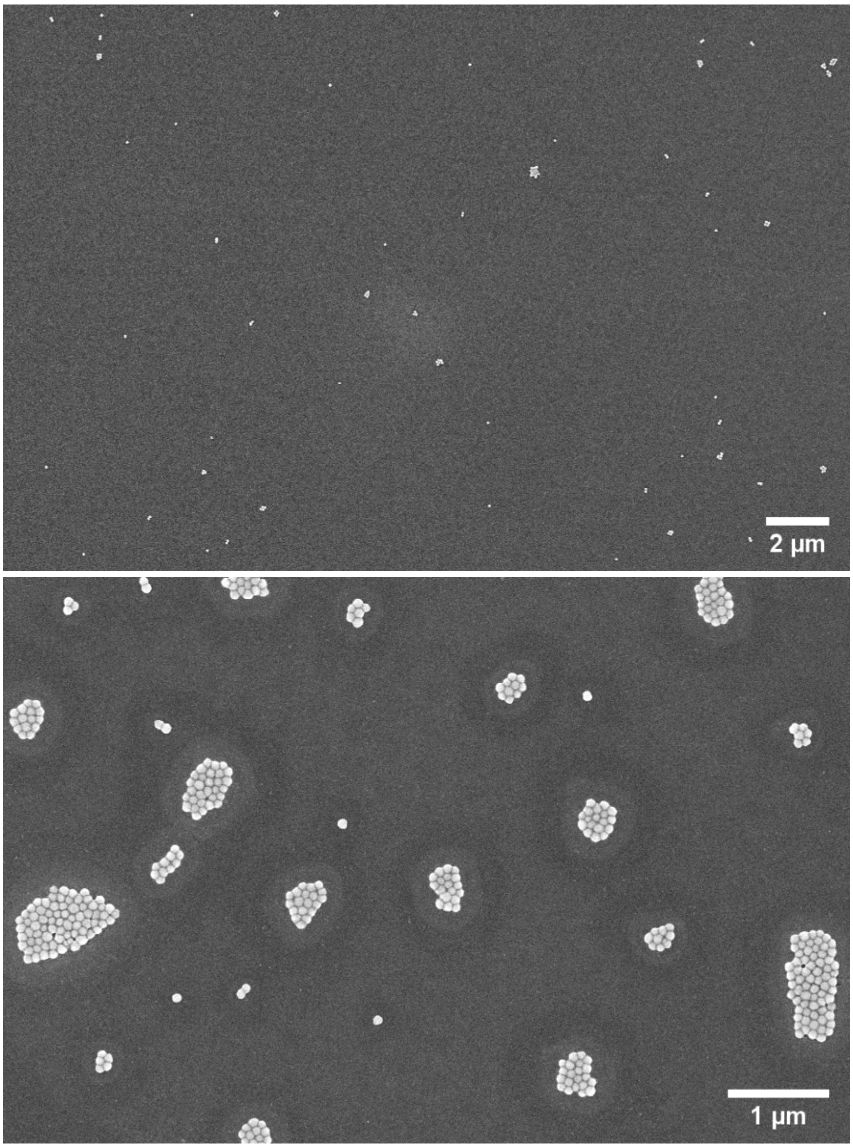

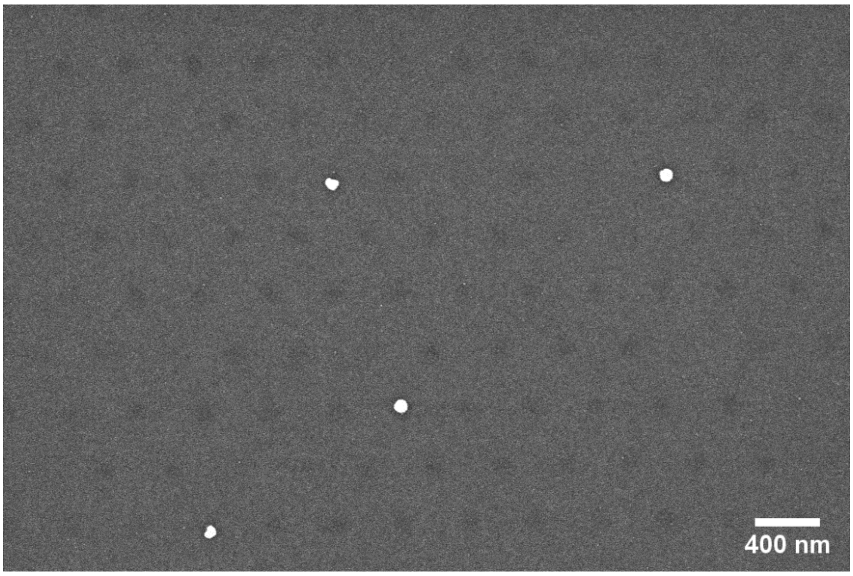


**Figure S16.** As a control, Trident is added directly to the binding sites, which does not allow the Trident to bind in the upright position as seen in DNA-PAINT experiments above, followed by NP incubation. We record SEM images showing minimal binding or formation of large aggregates in certain areas.

##### Figure S17. Snapshot after 100 pM assay with Trident without nanoparticles

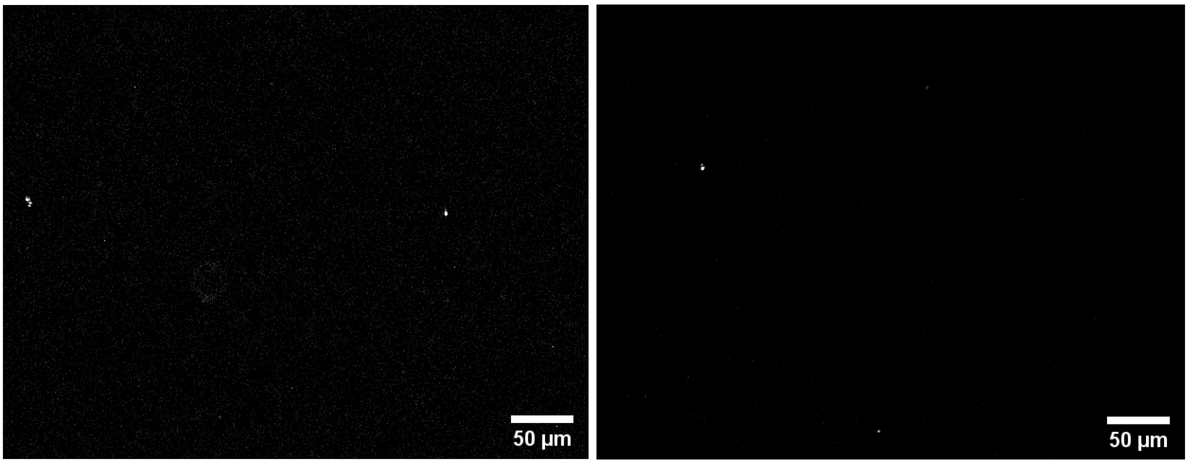


**Figure S17.** Zoom-in image from the reader, of scattering (left) and fluorescence (right) channels after running an assay with 100 pM T + 12 nM I + 12 nM B, with Trident immobilized on a BSA-biotin-neutravidin surface, in the absence of nanoparticles. In this case, a total of 9 spots were detected in the entire FOV by the analysis software, with no clear bleaching steps.

**Figure S18. Selective immobilization of antennas to cover two areas for measurements on the reader**

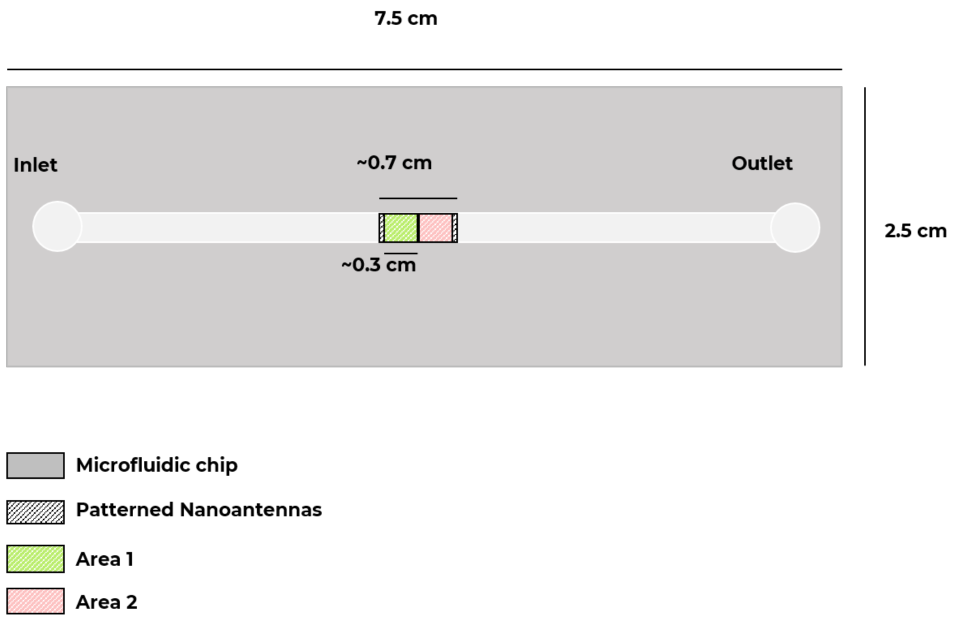


Figure S18. A schematic of the microfluidic chip showing the total area patterned (~0.7 cm x 0.3 cm) and the two areas imaged. The FOV of the reader is 2.5 mm x 3mm. The black and white sketched area is marked before patterning to place the NanoAntennas only within this region. Depending on the chip's orientation on the reader, two distinct regions can be imaged, highlighted in green and pink in the schematic. This orientation affects which area is centered within the reader's FOV, as the center alignment is slightly offset.

**Figure S19. Comparing assays on two chips patterned with NanoAntennas—one performed with repetitive flow and the other without.**

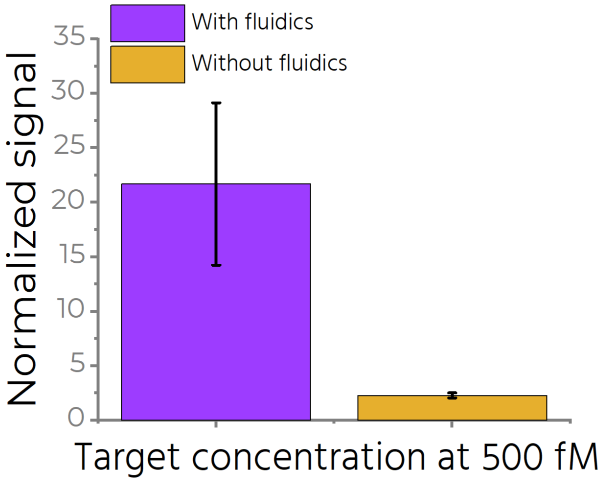


**Figure S19.** Applying flow allows us to achieve a ~10-fold increase in target capture. We demonstrate this by comparing assays on two chips patterned with NanoAntennas—one performed with flows and the other without, both with an incubation time of one hour and a target concentration of 500 fM.

**Figure S20. Exemplary zoomed in snapshots from different concentrations measured on the reader in buffer.**

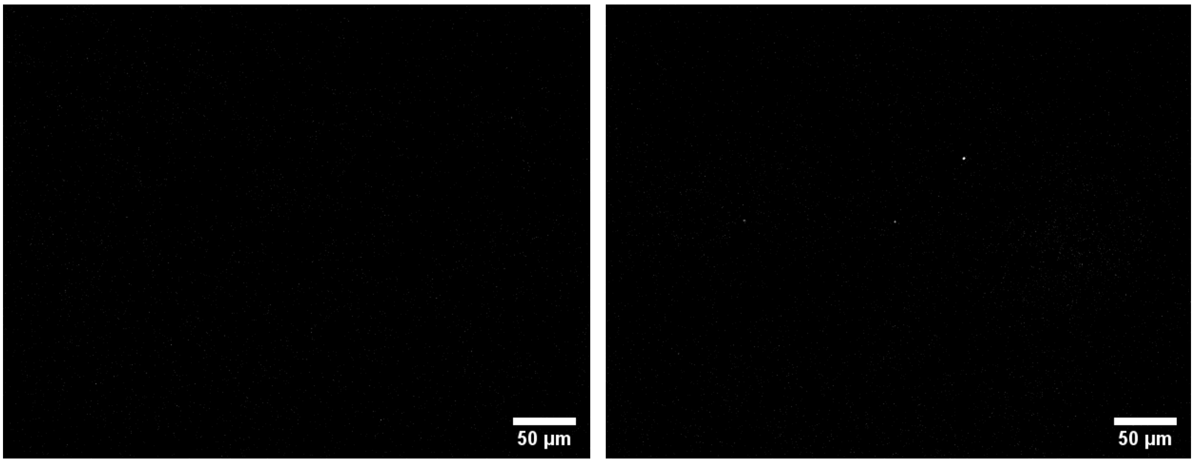


Blank 1 aM

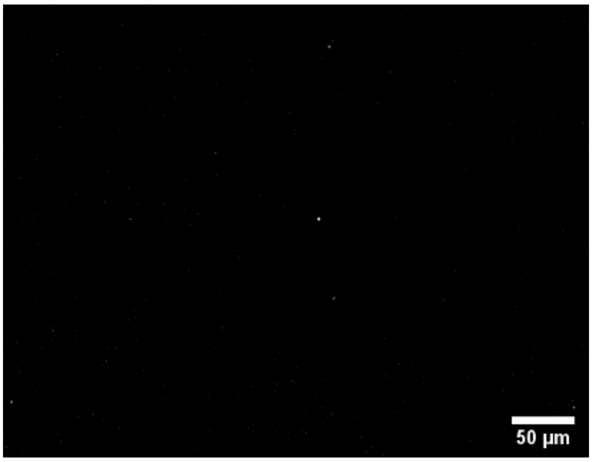


5 aM

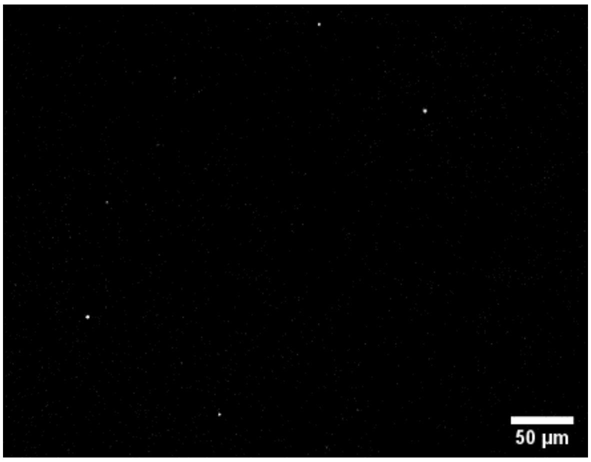


10 aM

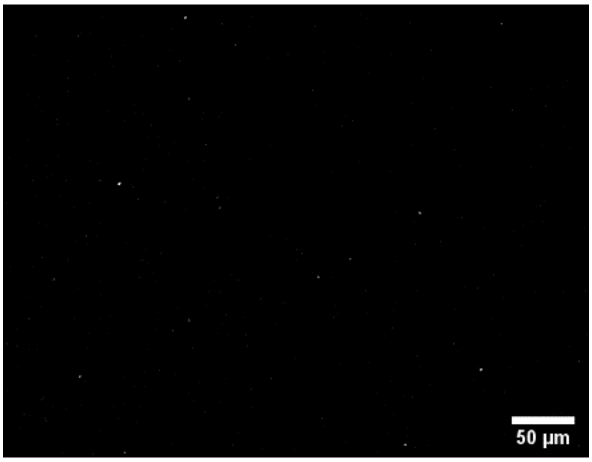


50 aM

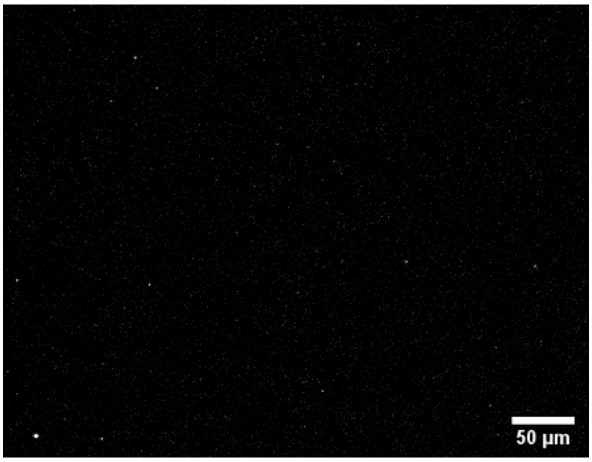


100 aM

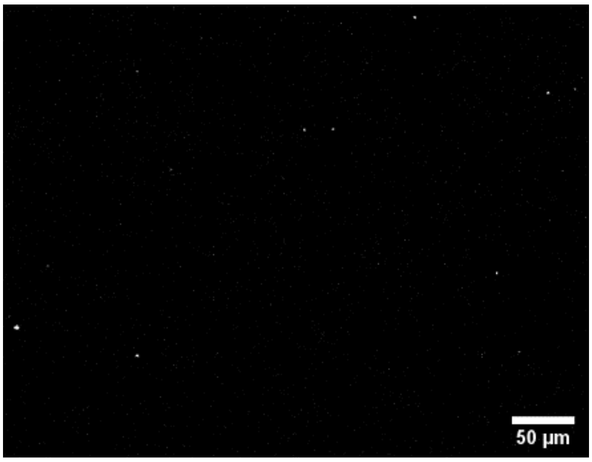


500 aM

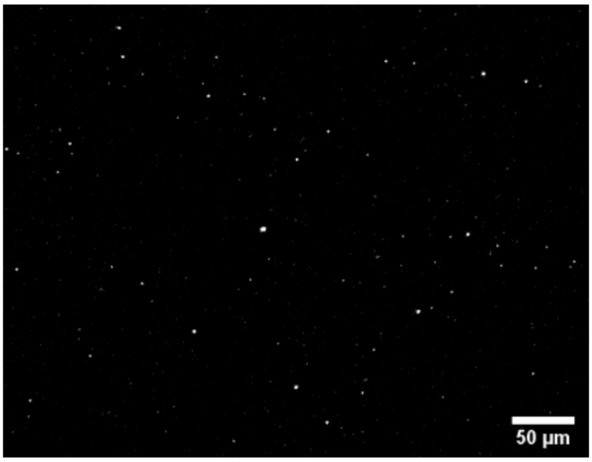


100 fM

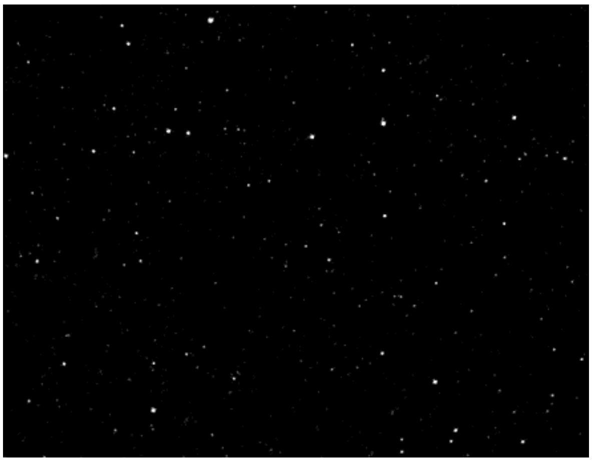

500 fM

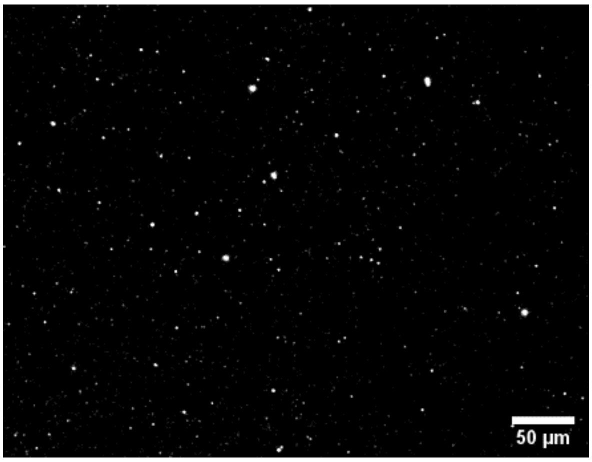


1 pM

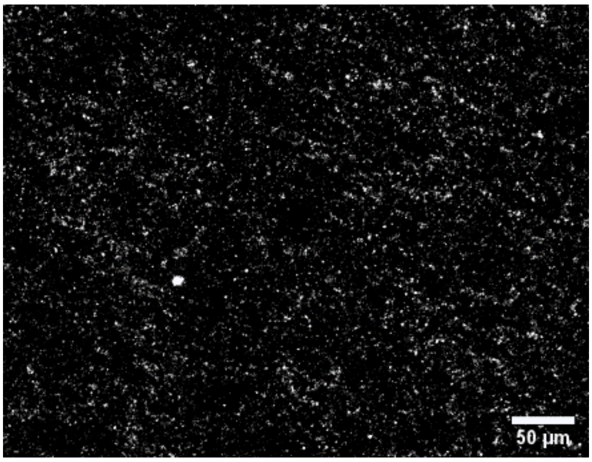


100pM

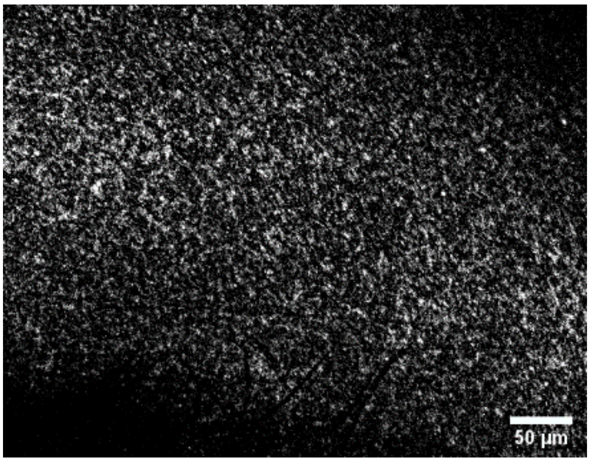


500pM

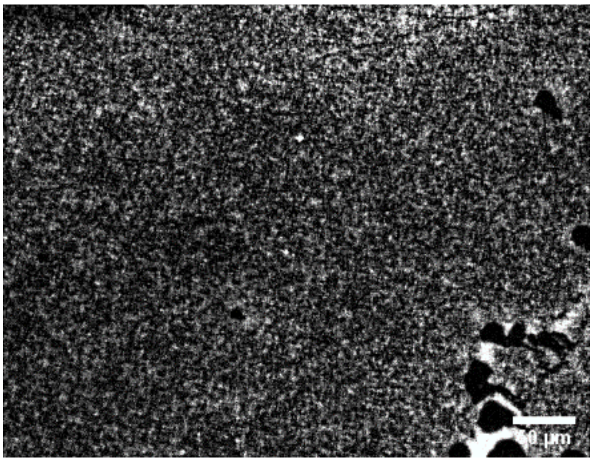

5 nM

**Figure S21. Washing with Sodium buffer helps remove weakly bound or unspecifically bound molecules.**

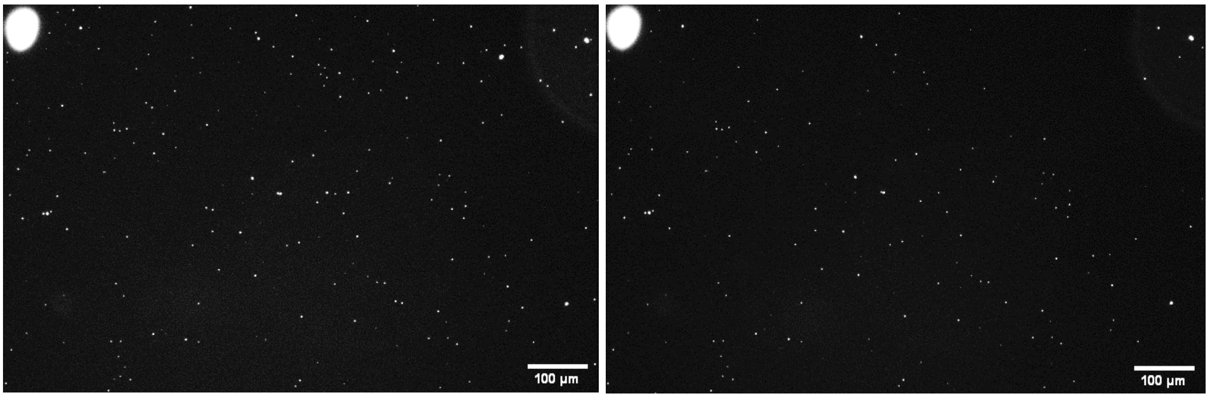


Figure S21. Snapshot of the fluorescence channel of a chip with silicified NanoAntennas after assay. Image on the left is after washing with $Mg^{2+}$ buffer and the image on the right is after washing the same sample with $Na^{+}$ buffer. The reflection of a bubble on the top left is used as a marker to compare the same areas.

**Figure S22. Effect of using Sodium-based buffer on patterning of unsilicified NanoAntennas**

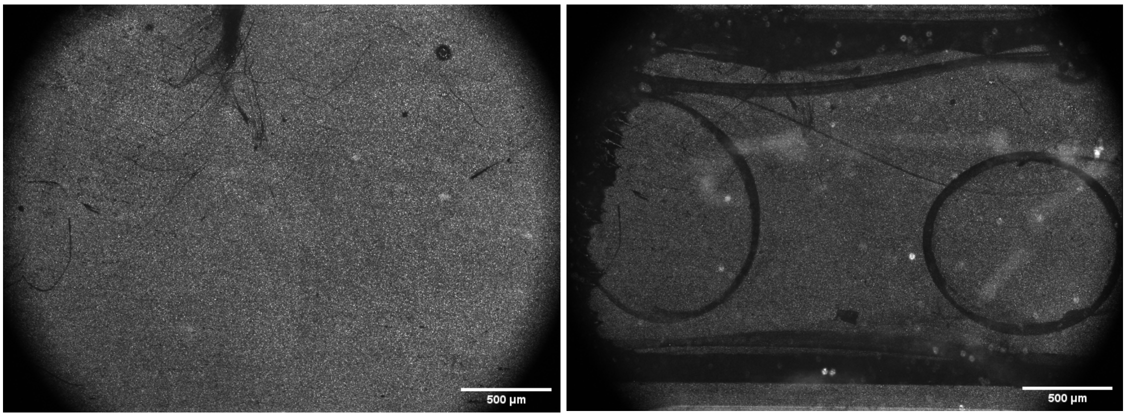


Figure S22. Snapshot of the scattering channel of the same area of a chip with un-silicified NanoAntennas before (left) and after (right) washing the chip with TE + 2M NaCl buffer. Washing the channel with sodium-based buffers causes lift-off of patterned NanoAntennas.

#### Figure S23. Effect of using Sodium-based buffer on patterning of silicified NanoAntennas

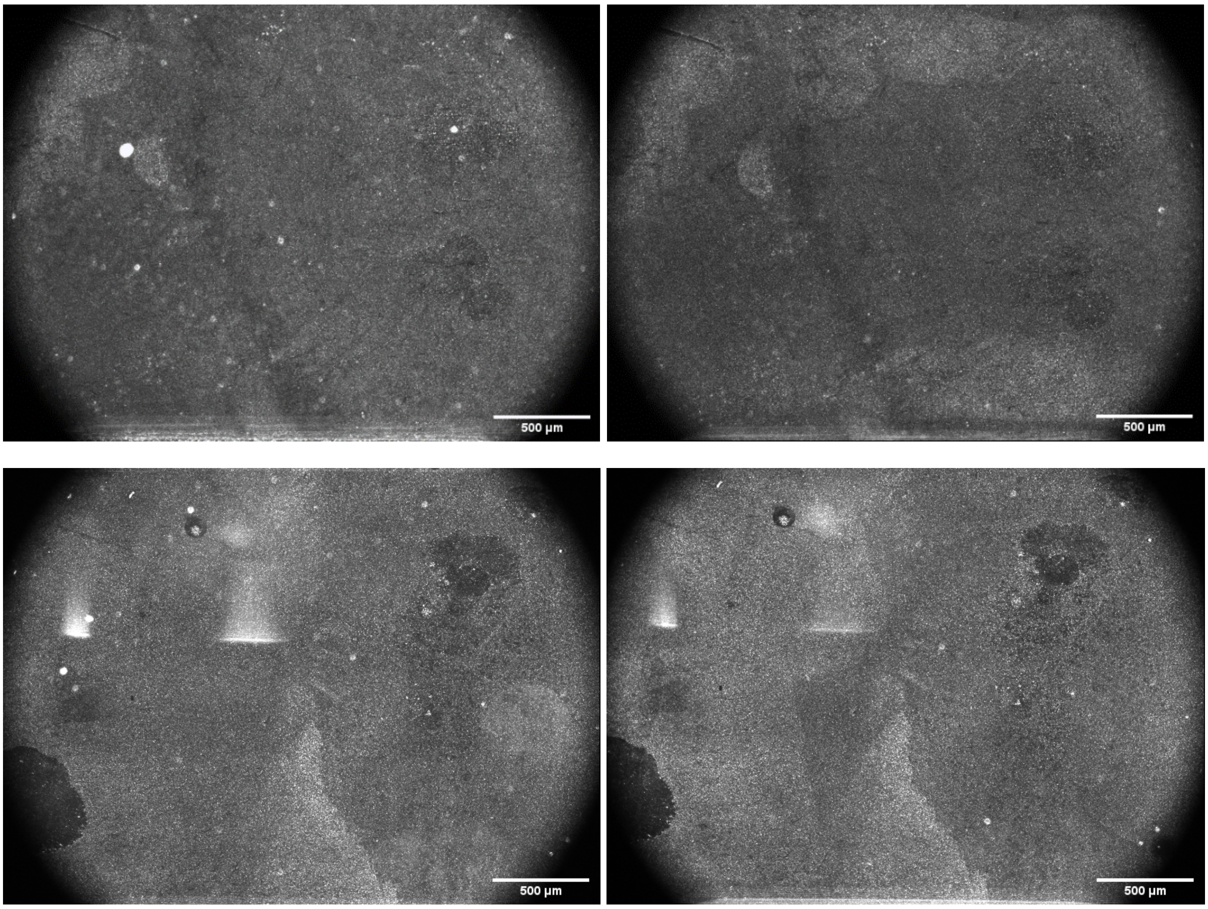


Figure 23. Snapshot of scattering channel from two silicified samples (top row and bottom row) before (left) and after (right) washing the chip with TE + 2M NaCl buffer. No visible lift-off is observed after silicification.

**Figure S24. AFM showing Trident before and after 1 and 4 days of silicification**

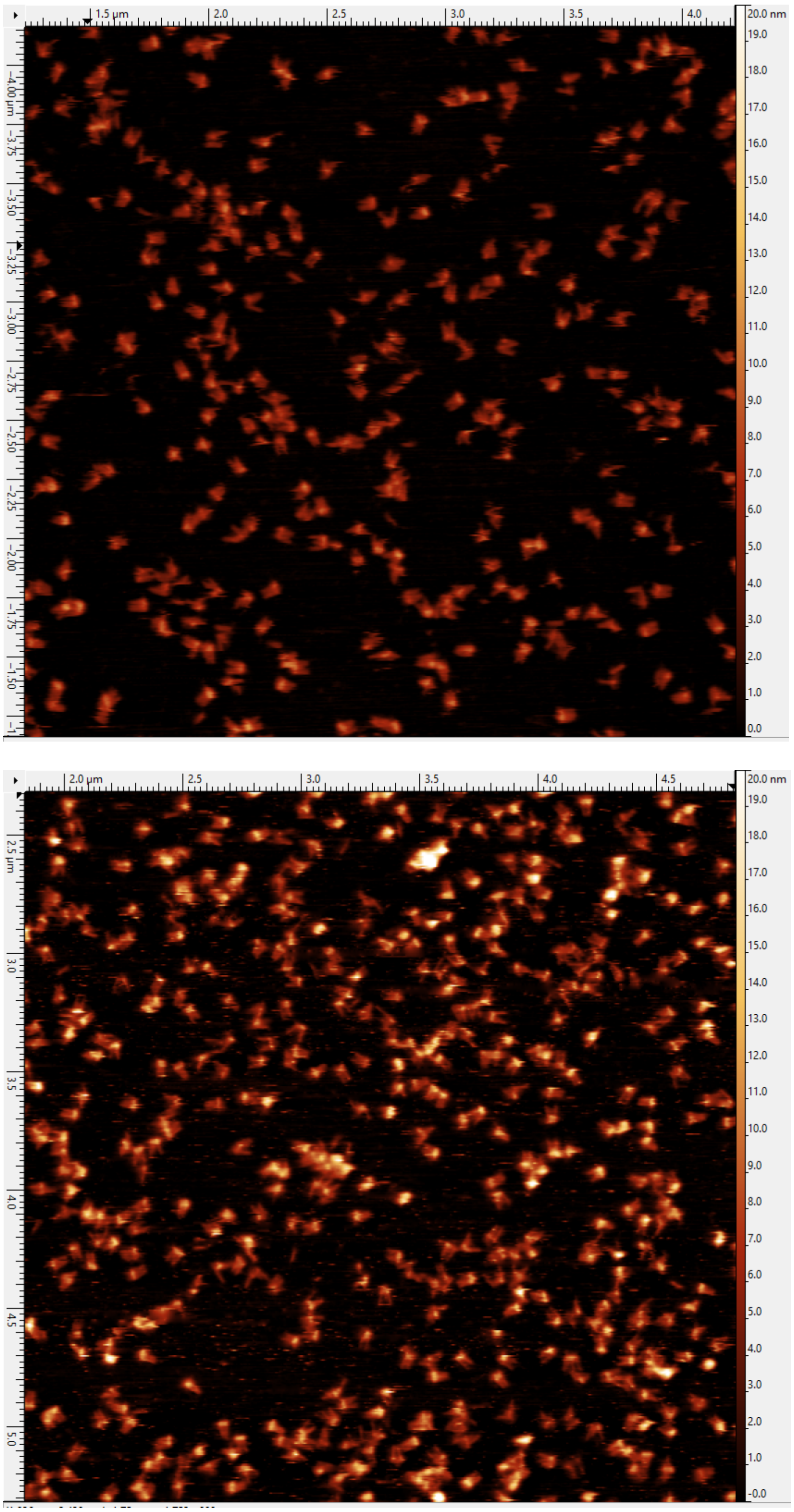

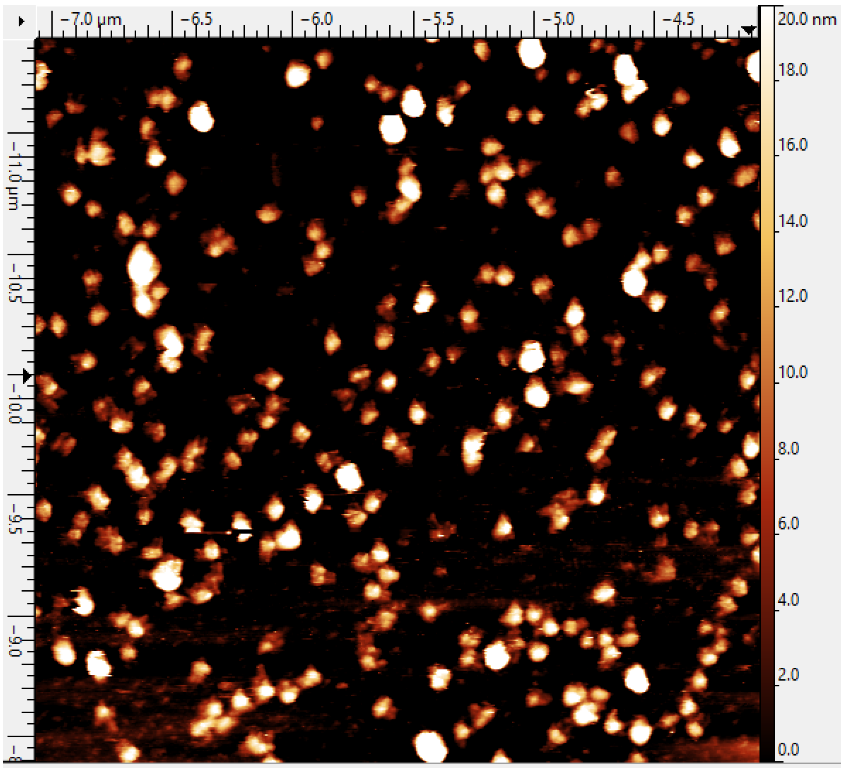


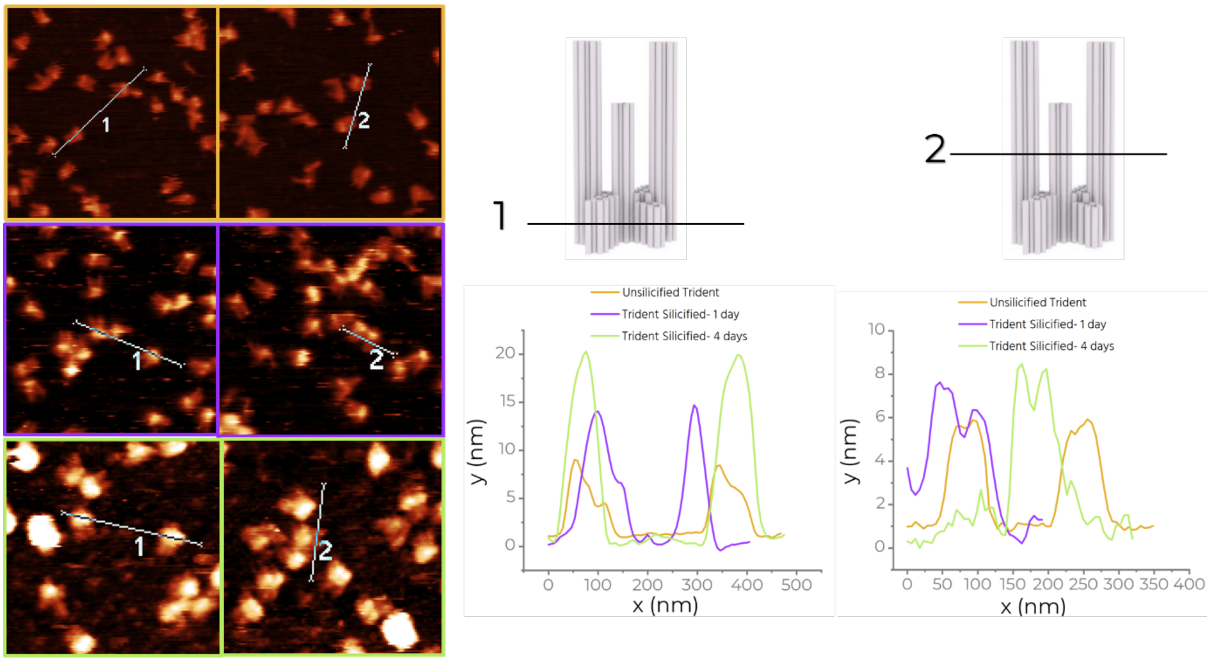


**Figure 24.** Example AFM images of Trident on mica before (first image), after 1 day (second image) and 4 days (third image) of silicification and line scan analysis showing height comparisons for each. After 4 days of silicification, it became difficult to distinguish structural features of the Trident. We also observe larger chunks of silica aggregates in this case.

#### Figure S25. Imager degradation in human blood plasma.

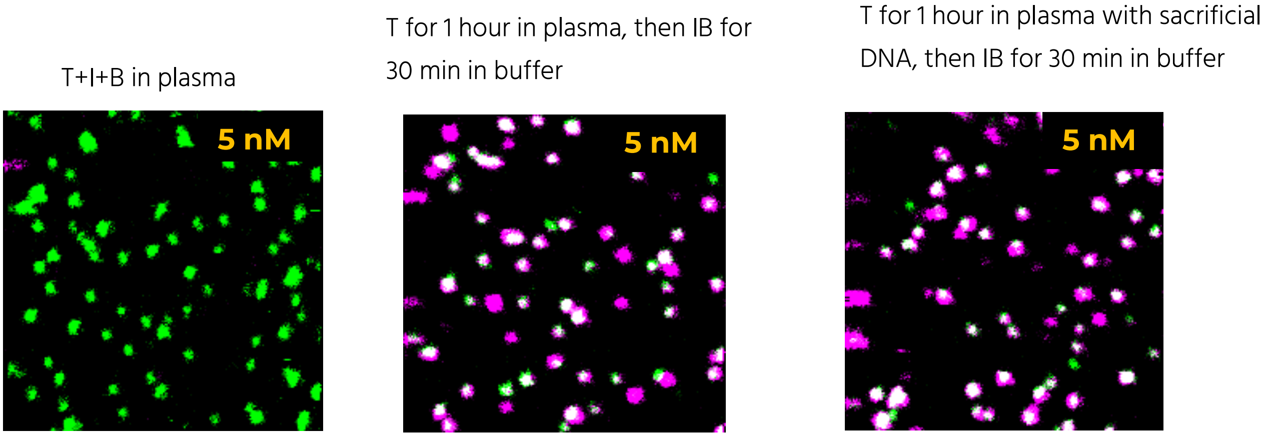


**Figure S25.** Fluorescence scans after incubating Target (5 nM), imager (12 nM) and blocker (12 nM) in human blood plasma for 1 hour with silicified NanoAntennas on the left, after incubating Target in plasma for 1 hour and then incubating imager and blocker in buffer for 30 minutes, after incubating Target with 1 $\mu$M random (sacrificial) DNA in plasma and then incubating imager and blocker in buffer for 30 minutes. All incubations are done in the incubator at 37°C.

#### Figure S26. Secondary structure: Target strand

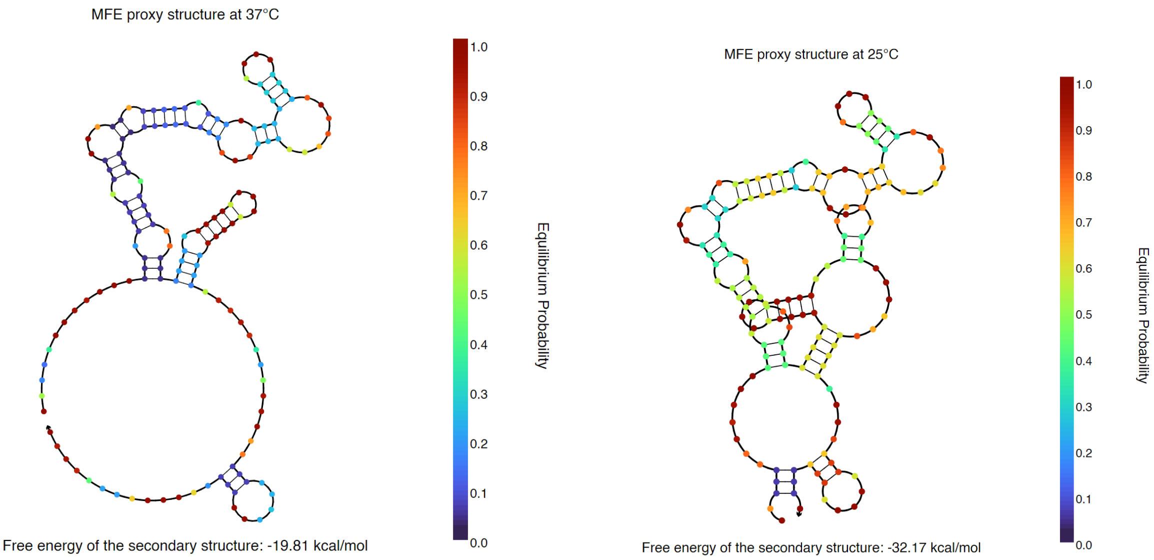


**Figure 26.** NUPACK analysis for secondary structure of the Target strand at 37°C (left) and at 25°C.

**Figure 27. Exemplary zoomed in snapshots from different concentrations measured on the reader in human blood plasma with silicified NanoAntennas.**

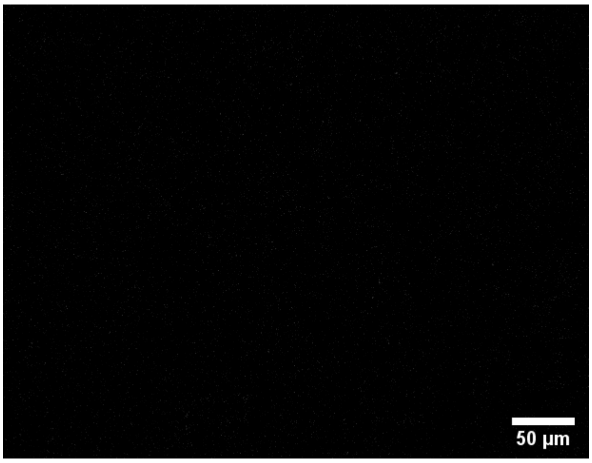


Blank

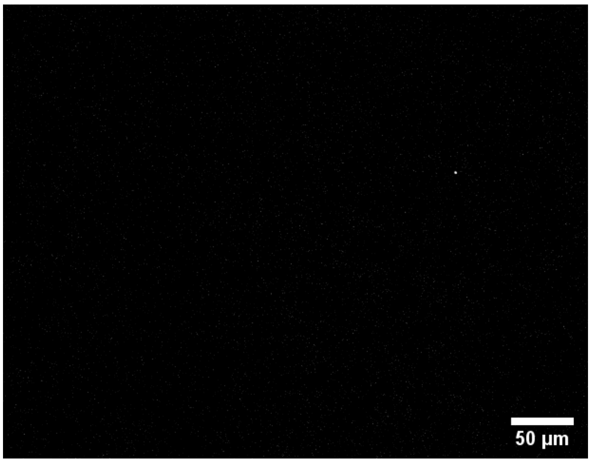


1 aM

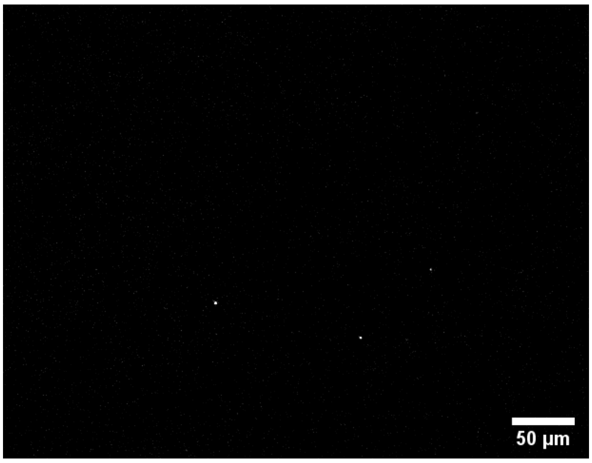


5 aM

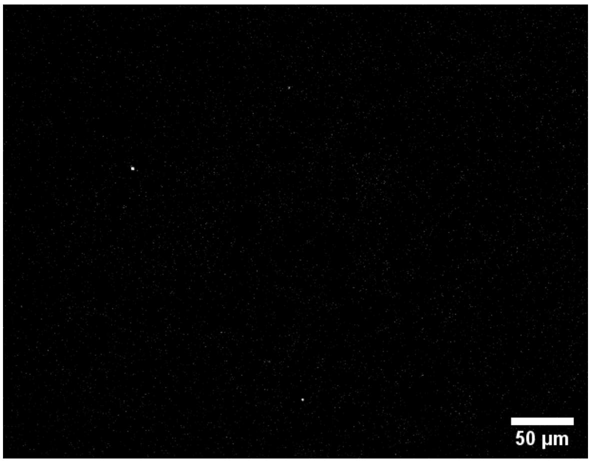


10 aM

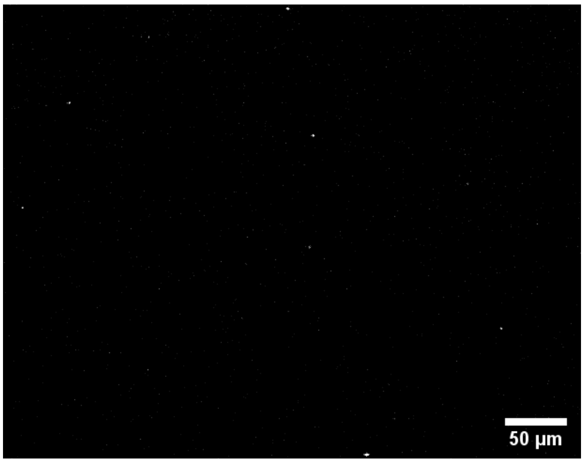


100 aM

**Figure 28. Reusable chips.** We do this by introducing a 'displacer' strand with 25 nt complementary to the target strand, which is thermodynamically more stable than the 17 nt interaction between the capture and target strands. Once displaced, the capture strand is available for use again (Figure 28a). To test this, we immobilize NanoAntennas on a BSA-biotin-NeutrAvidin modified glass substrate and incubate with 500 pM of target along with 12 nM of imager and blocker for one hour and record fluorescence scans (Figure 28b). We observe dye colocalization with a target binding yield of ~90% (Figure 28c). We then incubate the sample with a high concentration (1 μM) of displacer for up to two hours, reducing the bound fraction to ~30% (Figure 28b, c). Incubating overnight results in a <2% bound fraction. This likely arises from unspecific attachment of imager strands. We remove them through a short incubation with a high concentration of blocker strands, reducing the bound fraction to nearly zero. Upon re-incubation of the surface with 500 pM target, 12 nM imager and blocker, we achieve ~85% target binding yield. We perform the same protocol on our patterned NACHOS chip and successfully displace the bound target-imager complex, allowing the chip to be reused.

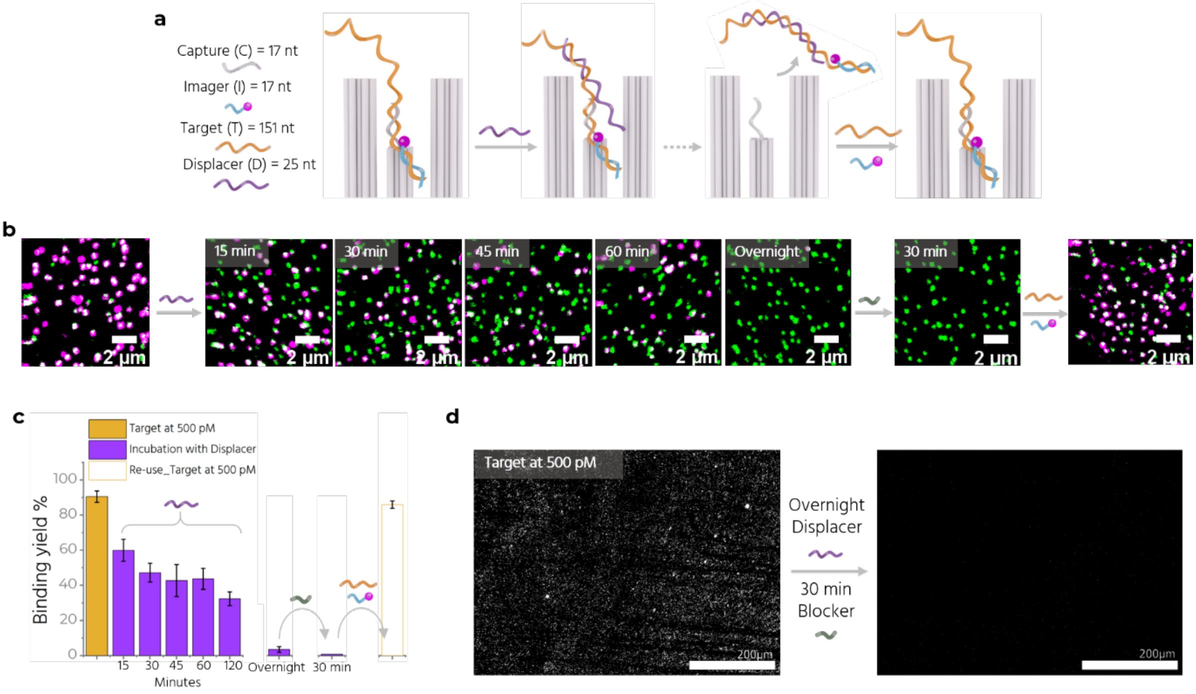


Figure 28. **a** Sketch showing the stand displacement of the target-imager complex by displacer strand, making the chip reusable. **b** Confocal fluorescence scans recorded after the 500 pM assay (left) followed by scans at 15, 30, 45, 60 minutes and overnight incubation with the displacer, after 30-minute incubation with blocker and after re-using the surface for the 500 pM assay. **c** Target binding yield over time. **d** Zoom-In of the fluorescence channel of a patterned NanoAntenna chip after an assay with 500 pM target (left). Incubating overnight with displacer followed by 30 minutes of blocker (right) displaces the target-imager complex, making the chip ready to be used again.

#### Supplementary Note. 1

##### 1.1 Klebsiella pneumoniae strain Kp11978 plasmid pOXA-48

We targeted a 151 nt sequence from the complete sequence that can be found at this link:

https://www.ncbi.nlm.nih.gov/nuccore/JN626286

##### 1.2. Design and purification of DNA origami nanostructures

**Trident DNA origami nanostructure:** The Trident utilized in this study incorporates a previously characterized design by Close et al.,[1] employing the p8064 scaffold derived from M13mp18 bacteriophages and designed using caDNAno software (version 2.2.0),[2] as depicted in Figure S1. The design integrates various elements including a base dye (ATTO 542), 12 biotin anchors, 12 nanoparticle binding staples, 6 PAINT docking strands, 6 strands for hybridization with the Triangle origami, and 10 capturing strands for assays. The scaffold is produced in-house, while both unmodified and modified staples are sourced from Integrated DNA Technologies, Inc. (IDT). The folding of the DNA origami structures is carried out in a one-pot reaction, mixing 30 nM of scaffold DNA with a 10-fold excess of unmodified oligonucleotides and a 25-fold excess of modified strands relative to the scaffold, in a folding buffer composed of 1x TE and 20 mM $MgCl_2$. This mixture underwent a detailed multistep thermocycling protocol, mentioned in Table S2.

**Triangle DNA origami nanostructure:** The Triangle origami is a modified version of the Rothemund Triangle, folded with p7249 scaffold produced in-house, derived from M13mp18 bacteriophages and designed using caDNAno software (version 2.2.0), as depicted in Figure S7. Modified strands In the structure include 6 strands for hybridization to the Trident and 6 strands as docking sites for DNA-PAINT measurements. The folding is carried out in a one-pot reaction, mixing 30 nM of scaffold DNA with a 10-fold excess of unmodified oligonucleotides and a 30-fold excess of modified strands relative to the scaffold, in a folding buffer composed of 1x TE and 12.5 mM $MgCl_2$. This mixture underwent a detailed multistep thermocycling protocol, mentioned in Table S2.

**Purification:** The origami structures are purified using Amicon Ultra filters (100 K, Merck, Germany).

1. Add 500 µl 1 x FoB buffer in the filter tube and spin at 10,000xg at 4°C for 5 minutes.
2. Remove the supernatant collected in the tube and add 100 µl of folded origami mix to the filter and add 400 µl of 1x FoB buffer and spin again.
3. Repeat Step 2 twice more with 400 µl of 1x FoB buffer.

4. Invert the filter and place into a new collection tube. Spin at 1000xg for 5 minutes.
5. Collect and store the purified origami in a low-bind Eppendorf tube.
6. Measure the concentration of the DNA origami using a Nanodrop 2000 spectrophotometer (Thermo Fisher Scientific), and store at -20°C for further use.

##### 1.3. Surface preparation: BSA-Biotin/NeutrAvidin coated coverslips

1. Clean the glass coverslips (24mm x 60mm; 170 µm thickness from Carl Roth GmbH, Germany or 25mm x 75 mm, 170 µm from Electron Microscopy Sciences, PA, USA by rinsing with Milli-Q water and isopropanol, then dry under a nitrogen stream.
2. Treat in a UV-Ozone cleaner (PSD-UV4, Novascan Technologies, USA) at 100°C for 30 minutes.
3. Affix Adhesive SecureSeal™ Hybridization Chambers (2.6 mm depth, Grace Bio-Labs, USA) to the coverslips and keep on a hotplate at 100°C while pressing on the chamber for 1-2 minutes.
4. Wash the chamber three times with 150 µl of 1x PBS buffer.
5. Add 150 µl of biotinylated BSA at a concentration of 0.5 mg/mL (Sigma-Aldrich, USA) and incubate for 5 minutes.
6. Wash the chamber three times with 150 µlof 1x PBS buffer.
7. Add 150 µl of NeutrAvidin (0.2 mg/mL, Thermo Fisher Scientific, USA.), and incubate for 5 minutes.
8. Wash the chamber three times with 150 µl of 1x PBS buffer.
9. Use the surfaces immediately for DNA origami immobilization.

##### 1.4. Surface preparation: Nanopatterned coverslips

The nanopatterning protocol we use was first introduced by Shetty et al.[3]

1. Mark a rectangle (0.7 cm x 0.5 cm) on one face of a 25mm x 75mm glass coverslip with a marker.
2. Wash both faces of the coverslip with Milli-Q water.
3. Wash the unmarked face with isopropanol and dry using an airstream.

4. Place the coverslip in a UV-Ozone cleaner (100°C for 30 minutes) with the unmarked side facing upwards, and position thicker pre-cleaned glass coverslips over them, leaving only the marked area exposed.
5. Take 350 µl of 400 nm sized polystyrene nanospheres (Distrilab) in a low-bind Eppendorf and centrifuge at 10,000xg for 5 minutes at 20°C.
6. Discard the supernatant and suspend the spheres in 350 µl of Milli-Q water.
7. Centrifuge twice more while washing with Milli-Q water.
8. Discard the supernatant and suspend and suspend the spheres in 100 µl of 25% ethanol.
9. Use the ozone-cleaned coverslip immediately.
10. Position it at a slight angle (30-45°) and add 10 µl of the cleaned nanospheres solution is to one corner of the marked rectangle and allow the solution to spread and air-dry for 15-20 minutes.
11. Carefully wipe spheres dried outside the marked area using a wipe (Kimtech Science™ wipes by Fisher scientific).

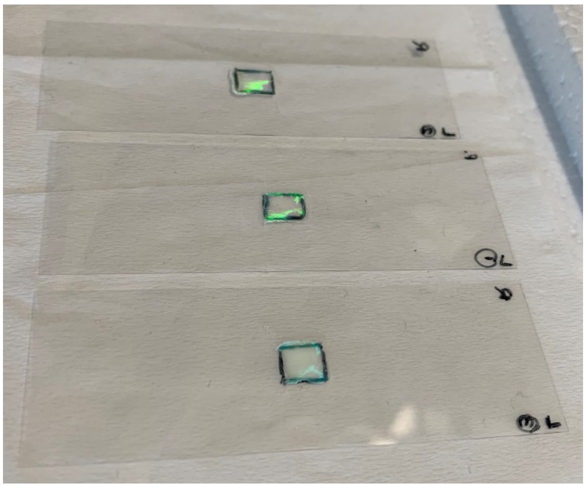

12. Heat the coverslip on a hotplate at approximately 60°C for 5 minutes.
13. Place in a sealed glass chamber with 5-10 ml of Hexamethyldisilazane (HMDS) by Merck, Germany, in a 10 ml glass beaker for 30-40 minutes.
14. Sonicate the coverslip in a beaker filled with Milli-Q water, fully submerging them until the spheres lift off, which usually takes about 30 seconds to a minute (Pipetting in the solution close to the patterned rectangle speeds up the process).
15. Dry with a nitrogen stream.
16. Heat at 120°C for 5-10 minutes, and store in a closed box at room temperature. Use immediately for DNA origami placement.

#### 1.5 DNA origami immobilization on BSA-Biotin/NeutrAvidin surface

100 pM of Trident modified with biotin anchors at its base, is diluted in 1xTE +2M NaCl buffer and added to a 150 µl chamber (prepared according to section 1.3), incubated for 1.5 minutes and washed 4 times in 1xTE +2M NaCl buffer. This gives a single-molecule density of the origami on the surface and was mostly used for measurements on the confocal microscope.

#### 1.6 DNA origami placement on patterned surface

This is a two-step process.

**1. Triangle placement:**

1. Add 200 µl of 150 pM Triangle diluted in placement buffer (PB) containing 40 mM Tris, 40 mM $MgCl_2$ at pH=8.4 to the coverslip (prepared as explained in section 1.4) and incubate for 1 hour.
2. Wash the coverslip 6 times with PB followed by 6 times with PBTween buffer (PB containing 0.05% Tween20) and 8 times (or longer until the surface becomes hydrophobic again) with PB.
3. Check the placement quality by DNA-PAINT (detailed description in section 1.16) using ONI Nanoimager. Assess the surface coverage, repeat the placement if needed.

**2. Trident placement:**

1. Add 200 µl of 500 pM Trident in PB to the coverslips with Triangles and incubate for 1 hour, followed by the same washing procedure as for Triangle.
2. Check the placement quality by DNA-PAINT using ONI Nanoimager. Assess the surface coverage, repeat the placement if needed.
3. If most of the Tridents are binding in random orientations on the Triangle (observed using DNA-PAINT), use a two-step, 1-hour long placement each with Trident concentration around 200-250 pM.
4. In case the concentration was too high and the surface has multiple origamis binding at each binding site, incubate the coverslip in a sodium-based buffer (~1 hour) to lift-off the origami and wash with PB 8-10 times and start again from Triangle placement.

#### 1.7 Functionalization and purification of nanoparticles

The freeze and thaw method[4] is used to functionalize silver nanoparticles (AgNPs) with DNA and agarose-gel electrophoresis is used to purify.

1. Add 100 µl of a 1 mg/ml AgNPs of 80 nm or 100 nm (nanoComposix) into a low-bind Eppendorf tube.
2. Take one tube of lyophilized thiolated-T20 DNA staples (4 nmol) from Ella Biotech, and add 675 µl of Sigma water. Mix thoroughly by shaking and vortexing. Slowly add this to the nanoparticle solution with constant pipetting.
3. Two such 675 µl aliquots are combined with the 100 µl NP solution in one tube.
4. Add 60 µl of 5 M NaCl with continuous pipetting.
5. Freeze at -20°C for 1-2 hours. Mostly left frozen overnight or until the day of use.
6. Thaw for 15-20 minutes wrapped in aluminum foil at room temperature on the day of use.
7. Centrifuge at 2800xg for 15 minutes at 4°C, discard the supernantant leaving 80-100 µl in the tube.
8. Add a tenth of the loading dye (BlueJuice Gel Loading Buffer, 10x by ThermoFisher scientific) before running in a 1.2% agarose gel at 100 V for 45 minutes.
9. Cut and squeeze the band to retrieve the functionalized nanoparticles.
10. Measure the concentration using Nanodrop 2000 spectrophotometer (Thermo Fisher Scientific). Store in dark at 4°C.
11. Sonicate before using, if not immediately used.

#### 1.8 DNA Origami NanoAntennas on BSA-Biotin/NeutrAvidin surface

If the NPs were not functionalized on the same day, they are sonicated before use and the concentration is measured again.

1. Dilute the AgNPs to reach an optical density (O.D.) of 0.05 in 1x TE buffer containing 2M NaCl.
2. Add this to chambers containing Trident DNA origami structures (prepared as described in sections 1.3 and 1.5) and incubate overnight in dark at room temperature.
3. The following day wash 4-5 times with 1x TE buffer with 2M NaCl. The NanoAntenna assembly is complete.

#### 1.9 DNA Origami NanoAntennas on patterned surface

If the NPs were not functionalized on the same day, they are sonicated before use and the concentration is measured again.

1. Dilute to an O.D. of 0.2 in PBTween buffer (40 mM Tris, 40 mM MgCl2, and 0.05% Tween20).
2. Add 50 µl of this to coverslips containing Trident DNA origami structures bound to Triangle origami (prepared as per sections 1.4 and 1.6) and incubate overnight in the dark at room temperature.
3. The following day, wash 5-6 times with PBTween buffer.
4. Store in PB until use.

#### 1.10 Addition of a microfluidic chip

The coverslips with nanonatennas are prepared according to section 1.9. Most of the liquid outside the marked area is carefully removed with a wipe without letting the surface dry. A straight channel microfluidic chip (Fluidic 268 by microfluidicChipShop) with self-adhesive tape is used. The tape is removed, and the chip is pressed onto the coverslip. After the chip is firmly glued to the coverslip, the channel is washed 3-4 times with PBTween. The inlet and outlet are blocked while the center of the chip is sprayed with a black paint (Buntlack Matt from OBI, Germany), to reduce the autofluorescence from the chip. The chip is ready once the paint has dried (~15 minutes air dry).

#### 1.11. Silicification

The protocol employed for silicification was adapted from the method outlined in here[5,6]

1. Take a 10 ml glass vial and add 3 ml PB.
2. Add a magnetic bead and stir at 900 rpm at room temperature. Make sure to keep the stirring unobstructed.
3. Slowly add 60 µl of TMAPS (N-[3-(trimethoxysilyl)propyl]-N,N,N-trimethylammonium chloride, 50% wt/wt in methanol, TCI America) to the mixture and stir for 20 minutes.
4. Slowly add 60 µl of TEOS (Tetraethyl orthosilicate, 98%, Sigma Aldrich) and stir for 20 minutes.
5. Add 100 µl of the precursor to each microfluidic chip, gently mix and keep the chips upside-down.
6. Incubate overnight (19-24 hours) at room temperature.

7. Remove the precursor and rinse thrice with Milli-Q water and thrice with absolute ethanol.
8. Let the chips air dry.

#### 1.12 Sandwich Assay

The sandwich assay includes incubation with the 151-nt target strand, 17-nt imager strand and 17-nt blocker strand. The imager and blocker are kept constant at 12 nM concentration. The target concentration varies depending on the experiment.

For measurements with BSA-biotin/NeutrAvidin surfaces, 1xTE 2M NaCl is used during the assay unless stated otherwise. For measurements with patterned surfaces, PBTween was used unless stated otherwise. The target, imager and blocker are incubated at 37°C for 1 hour in an incubator or coupled with back-and-forth microfluidic flow at 36°C for 1 hour on the reader.

Specific signal is determined by the target capture efficiency % in presence of both target and imager strands, while unspecific signal is calculated by the target capture efficiency % in presence of only imager strands.

For measurements with human blood plasma with patterned surfaces on the reader, the target is first incubated with 50% plasma, 40 mM Tris, 40 mM $MgCl_2$, 5 mM EDTA, 1 µM sacrificial DNA and 2 units of DNase I in water for 45 min with back-and-forth flow (at 36°C), washed 6-8 times with PBTween and with 1xTE 2M NaCl, then incubated 15 minutes with Imager and Blocker in PBTween with back-and-forth flow at 36°C, washed and imaged.

#### 1.13. Transmission electron microscopy (TEM)

The folding of the Trident DNA origami nanostructures was characterized with transmission electron microscopy (TEM). 5 µL of a sample was incubated for 30 s – 5 min, depending on concentration, on glow discharged TEM grids (formvar/carbon, 300 mesh Cu; Ted Pella) at room temperature. After incubation on the grids, the sample was wicked off by bringing the grid into contact with a filter paper strip. For the DNA nanostructures, a 5 µL drop of uranyl formate staining solution (2% uranyl formate aqueous solution containing 25 mM sodium hydroxide) was applied to the grid, immediately wicked off, followed by applying another 5 µL drop of uranyl formate staining solution. This drop was allowed to incubate on the grid for 10 seconds and then wicked off. The grid was dried for 5 minutes before imaging. Imaging was performed with a JEM1011 transmission electron microscope (JEOL) operated at 80 kV.

#### 1.14. SEM

The SEM instrument used in this work is the Raith eLINE SEM instrument. The beam settings for imaging are 10 kV acceleration and 20 μm aperture. The samples were imaged using the SEM after 20 s sputtering using an Edwards Sputter Coater S150B. The sputter target contained 60% gold and 40% palladium. The process parameters used for sputtering were 5 mbar Argon, 1.5 kV, 11 mA. Here 20 s of sputtering results in the deposition of a layer of gold/palladium with a thickness of a few nanometres. SEM imaging was performed on horizontal samples.

#### 1.15 AFM

AFM scans in aqueous solution (AFM buffer = 40 mM Tris, 2 mM EDTA, 12.5 mM Mg(OAc)2·4 H2O) were realized on a NanoWizard® 3 ultra AFM (JPK Instruments AG). For sample immobilization, a freshly cleaved mica surface (Quality V1, Plano GmbH) was incubated with 10 mM solution of NiCl2 for 3 minutes. The mica was washed three times with ultra-pure water to get rid of unbound Ni2+ ions and blow-dried with air. The dried mica surface was incubated with 1 nM sample solution for 3 minutes and washed with AFM buffer three times. Measurements were performed in AC mode on a scan area of 3 x 3 μm with a BioLeverMini cantilever (νres = 110 kHZ air / 25 kHz fluid, kspring = 0.1 N/m, Bruker AFM Probes). Leveling, background correction and extraction of height histograms of obtained AFM images were realized with the software Gwyddion[7] (version 2.60).

#### 1.16 DNA-PAINT

DNA-PAINT measurements were carried out on a commercial Nanoimager S (ONI Ltd., UK). Red excitation at 640 nm was realized with a 1100 mW laser, green excitation at 532 nm with a 1000 mW laser, respectively. The microscope was set to TIRF illumination and a pixel size of 117 nm.

**DNA-PAINT with Triangle:** The Triangle DNA origami structure is modified to incorporate six docking sites, each with an 8-nt sequence. These docking sites are positioned to ensure that the super-resolved image represents a triangle shape. The docking sites are oriented towards the interior of the Triangle. This orientation ensures that the docking sites remain accessible for interaction, regardless of which side of the Triangle is facing the surface upon immobilization. For the measurements, ~300 pM of 8 nt Aptamer imager (complementary sequence to the docking site) in PBTween buffer, labeled with ATTO 655 is added to the coverslip and the transient binding of the imager to the docking sites is observed over 10,000 frames at 100 ms exposure time on the ONI Nanoimager. The data is processed using Picasso Localize and Picasso Render.[8]

**DNA-PAINT with Trident:** The Trident consists of 6 docking sites (8-nt long), 3 at the bottom of the left pillar complex and 3 at the top of the left pillar complex such that if the Trident is immobilized perpendicular to the Triangle (desired orientation), the output super-resolved image would be one circular or elliptical (flexibility of the pillar at the top) spot, while a parallel orientation would result in two spots separated by a distance equivalent to the height of the structure ~74 nm. For the measurements ~300 pM of 6-nt fast imager (complementary sequence to the docking site) in PBTween buffer, labeled with ATTO 655 is added to the coverslip and the transient binding of the imager to the docking sites is observed over 10,000 frames at 100 ms exposure time on the ONI Nanoimager. The data is processed using Picasso Localize and Picasso Render.

##### 1.17 Confocal measurements

532 nm wavelength is used to excite ATTO 542 and 639 nm for Alexa fluor 647. 20 µm x 20 µm scans are recorded with 1 ms integration time. Samples are excited at 1 µW laser power. For enhancement factor measurements, samples without NPs are excited with 500 nW and samples with NPs at 50 nW. The confocal setup used is as described by Trofymchuk, Glembockyte et al.[9] The data acquired is processed with a custom-made LabVIEW software (National Instruments, USA) and further analysis was done with OriginPro2019.

#### Supplementary Note. 2

##### Reader specifications

The fluidic automation is implemented with the help of a syringe pump and a needle, whereby the needle can be moved in a motorized manner between liquid reservoirs and a reaction chamber. It is therefore a miniaturized pipetting robot. The cartridge is designed accordingly (Figure A): It has liquid reservoirs that are arranged next to each other and an inlet into a reaction chamber that can be read optically. All reservoirs and the inlet can be closed with septa, which close tightly again after perforation by the needle. The cartridge therefore represents a closed system.

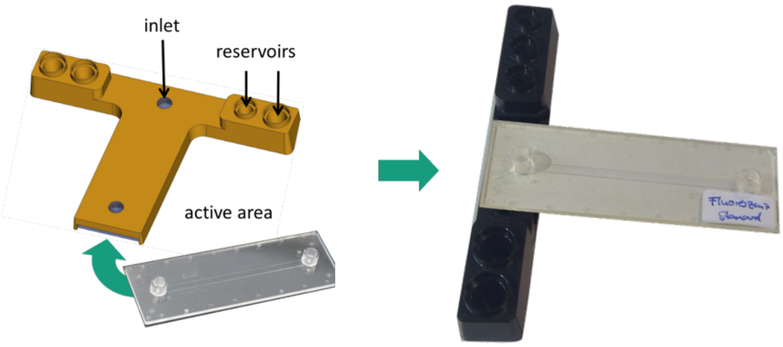


**Figure A.** Sketch of the 3D-printed chip holder with reservoirs on the left and the microfluidic chip shown on right.

Fluidics is implemented as a modular concept for validation purposes. A polymer carrier contains all access points for the needle including the septa. This carrier was created as part of the project using a 3D printer. There are several design variants that particularly affect the reservoirs. These can either be filled with chemicals directly in the carrier or equipped with standard vials. The latter variant is not suitable for mass production, but it has made the laboratory processes in the project easier. The polymer carrier can be produced economically in large quantities as an injection molded part. The fluidic chip with the chamber for optical readout is inserted into the carrier. This is constructed by first capping the chemically prepared glass substrate with a polymer chip made of Zeonex® cycloolefin polymer (COP) with high optical quality. This creates the flow cell for the detection reaction.

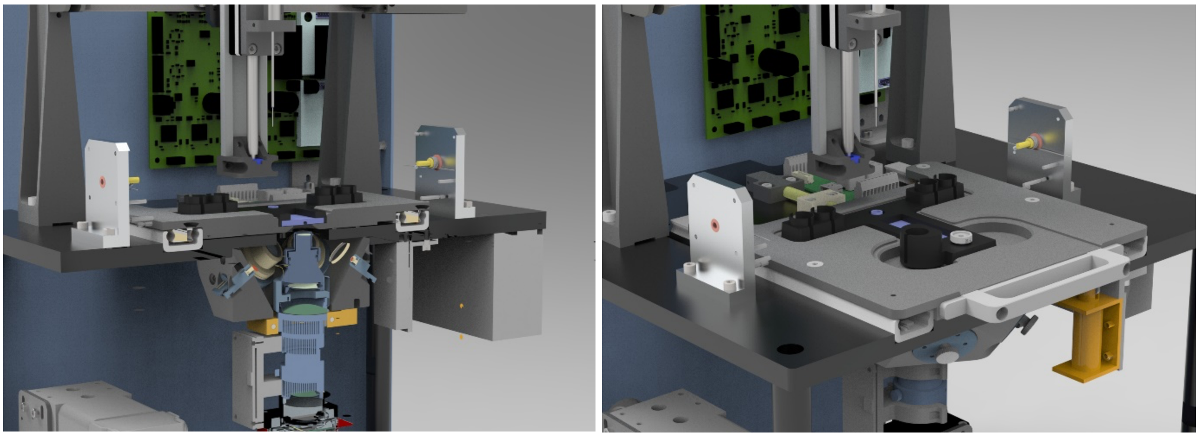

**Figure B.** A sketch of the inside of the reader focusing on the optics unit on the left and pipetting needle and the sample holder on the right.

The optical unit consists of the illumination to stimulate fluorescence and a microscope including a camera for imaging (Figure B). Using spectrally filtered LEDs, an optical power density of 2 $W/cm^2$ is achieved, which is sufficient to detect individual molecules. The microscope has a so-called tandem lens - two standard camera lenses arranged in opposite directions. The detection filter is located in the area of the parallel beam path. One of the two lenses has a liquid lens that can be used to electronically adjust the focal plane. By choosing a liquid lens, the entire optical system has no moving parts. The imaging is done on a CMOS camera with 12-megapixel resolution. With a numerical aperture of 0.2, this microscope achieves an optical resolution of < 2.7 µm (0.8 µm per pixel) over an image field of 2.5 × 3 $mm^2$. The optics are extremely compact with a length of only 16 cm and can be set up at a moderate cost. The image field is large in order to be able to perform multiplexing using several spots and to collect sufficient data for the measurements to be statistically meaningful.

**Reader data and analysis:** Samples are focused using the scattering signal from the NPs (See Figure A). Samples are imaged before doing the assay to ensure there is no unspecific signal in the fluorescence channel (Figure C). The microfluidic chip with the assay mixture is loaded in the sample holder and an automated script allows a pipetting needle to create a back-and-forth flow in the channel. A heating block that touches the chip on top is kept at ~36°C during the incubation. The script is written to continue for 1 hour after which the chip is taken off and washed 5-6 times with PBTween. The sample is excited by spectrally filtered LEDs for several frames at 100 ms exposure time with a 500 ms gap between each frame. To avoid overheating of the sample by the LEDs, the movie recording is done in groups of 15 frames, after which a pause of 1-2 minutes is given before the next 15 frames. This is repeated until most of the single molecules are photobleached.

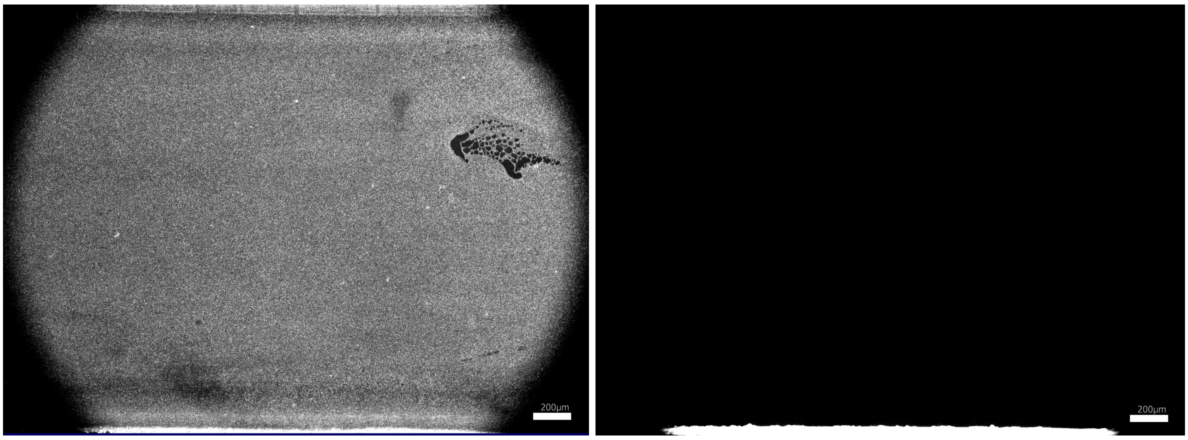


**Figure C.** Image taken on the reader before performing the assay. The left image shows scattering from nanoparticles, allowing us to focus on the right plane. The darker holes are defects from the nanopatterning. Image on the right is the fluorescence channel (after background subtraction), showing no signal before the assay. The bright part at the bottom of the Image Is the autofluorescence from the edge of the microfluidic chip.

After recording several frames, the software subtracts the background and counts single molecules (Figure D). The intensity threshold is set to capture most molecules, and if there is significant variation in intensity, two different thresholds can be used, ensuring zero overlap between spots. The software also plots bleaching steps for each identified spot. However, with the current code, the software cannot accurately identify spots that do not blink or bleach. A masking function allows the user to exclude certain spots, which can be useful for eliminating reflections from bubbles that may move over time and cause multiple spots to be counted. Masking these bubbles or similar defects can prevent this issue.

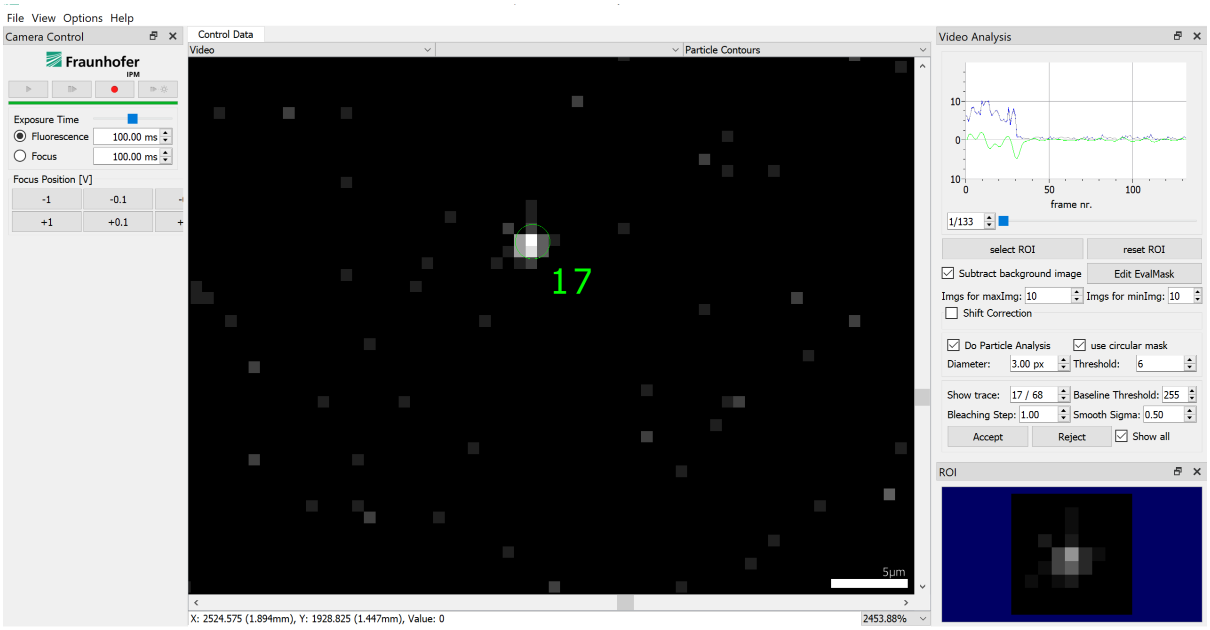


**Figure D.** A snapshot of the software showing a detected single molecule and the time transient on the top right showing a single bleaching step.

At lower target concentrations, the software can automatically detect single molecules (Figure E left). However, at higher concentrations, the surface becomes densely populated, resulting in inaccurate counting due to spots appearing as clusters or continuous bright areas (Figure E right).

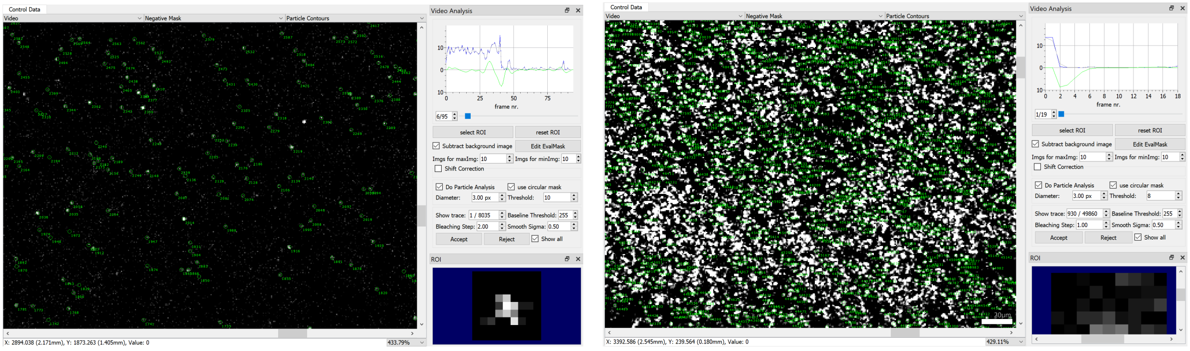

**Figure E.** A snapshot of the software with multiple single molecules identified accurately in case of lower target concentrations on the left, and the inaccurate counting at high target concentrations shown on the right.

#### Normalized signal

By observing the scattering from the NPs, we can detect variations in NanoAntenna surface density between the chips due to defects in patterning, as well as varying brightness due to heterogeneity in NP binding. To compensate for this and enable comparison of the chips at different target concentrations, we divide the total number of detected spots by the product of fraction of pixels in the image and the mean brightness of the pixels (after background subtraction) in the scattering channel. We refer to this value as the 'normalized signal' (N) defined as:

$$\mathrm{N} = \text{Mean scattering brightness}\ (\,\overline{\mathrm{B}}\,) * \text{Area covered}\ (\mathrm{A})$$

where B and A are computed using the scattering by nanoparticles. By illuminating the surface with white LEDs, we observe scattering from nanoparticles that allows us to focus on the sample without using fluorescence LEDs which would result in photo bleaching of the fluorophore while focusing. This scattering channel also shows the variability in nanoparticle binding across the chip. We use the following process to compute N by using ImageJ software:

1. Load the image in ImageJ.

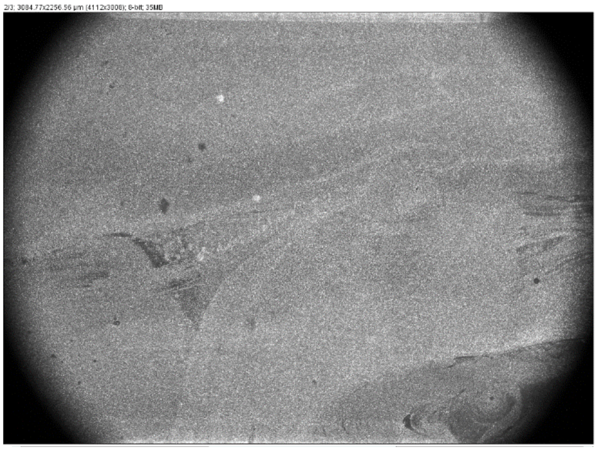

2. Subtract the background by using rolling ball radius.

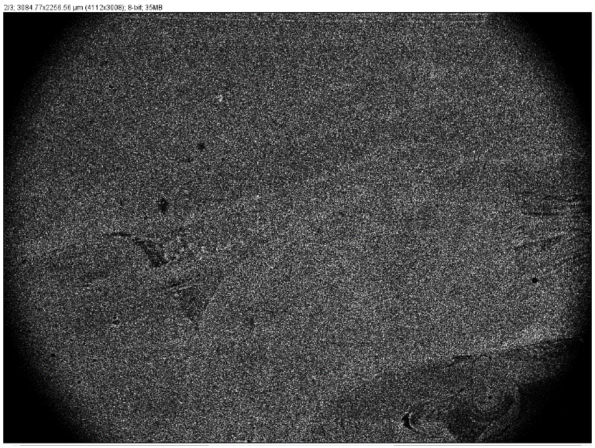

3. Use the select tool to roughly select the circular FOV.

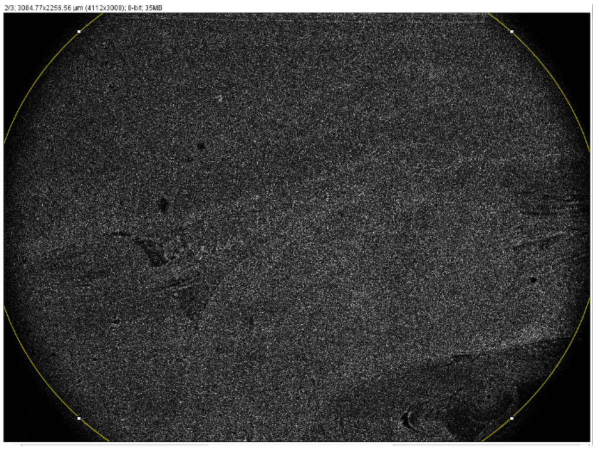

4. Measure the Mean gray value (the mean gray value is the sum of the gray values of all the pixels in the selection divided by the number of pixels.). This gives us an average brightness or B.

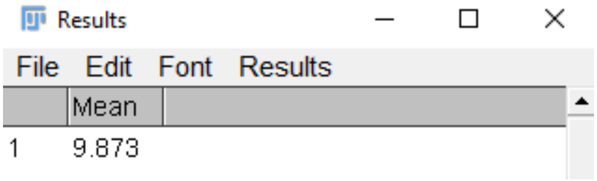

5. Use the threshold option to select the desired intensity range (included pixels are shown in red). Apply the threshold. This converts the image into a binary image, where pixels are either foreground (typically white) or background (typically black).

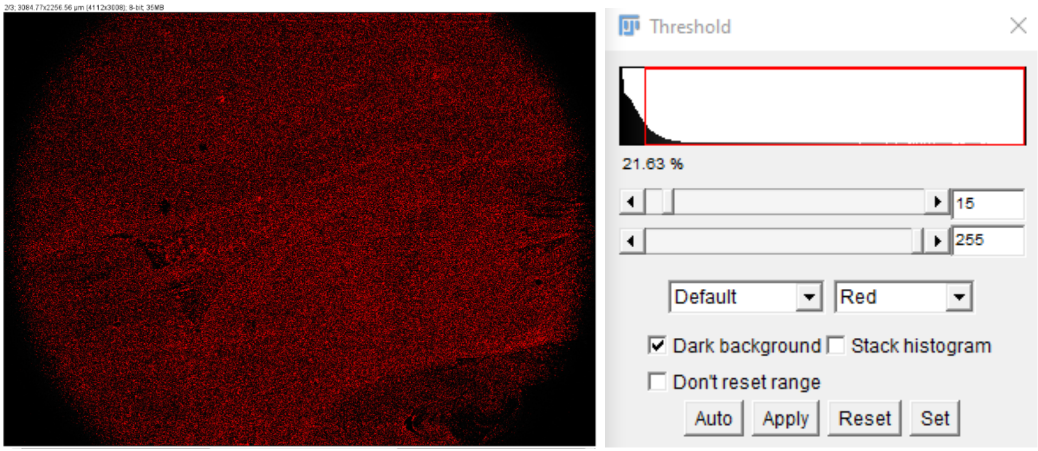


6. Use the select tool to roughly select the circular FOV and measure the 'area fraction' or the percentage of pixels in the image or selection. This gives us the % area covered or %A.

We covert the %A to decimal before using it for calculations. Normalized signal is thus given by:

$$\text{Normalized signal}=\frac{No.of\ spots\ in\ the\ fluorescence\ channel}{Mean\ scattering\ brightness\ (\,\overline{\mathrm{B}}\,)\ *Area\ fraction\ (A)}$$

As we measured two areas (See figure S18) for each sample by changing how the chip was placed in the sample holder, we combine the signal from both areas to arrive at the final signal given as:

$$\text{Normalized signal}=\frac{Total\ no.of\ spots\ in\ the\ fluorescence\ channel}{(B_1A_1)+(B_2A_2)}$$

#### Intensity-based analysis

1. Open the after-assay image with both scattering and fluorescence channel in ImageJ.
2. Look for defects in patterning in the scattering channel, select a circular area, check mean grey value and accordingly set a value to subtract background. This value for our samples was mostly around 20.
3. Set rolling ball radius to 20 and subtract the background from both channels.
4. Select the image with a rectangle, avoiding edge of the chip from the selection. Keep the same selection for both channels.
5. Use the 'transform image to results' option to get xy coordinates for each pixel for the fluorescence channel.

6. Save the files separately as .csv.
7. Import the files in Origin2019.
8. Delete the first column and select the rest of the data. Stack the data in a single column.
9. Select the data to get the sum of pixels.
10. Use the steps from the previous section to get the mean brightness and area covered for the scattering channel.
11. Divide the sum of pixels for the fluorescence channel by the product of mean brightness and area covered to get the 'normalized signal'.

#### Double-event estimate

At low target concentrations, compatible with single-molecule counting, the number of fluorescent molecules captured within a single pixel in the reader FOV follows a Poisson distribution, with the huge majority of pixels containing no molecules at all. This is well justified if one considers the high number of capturing strands contained in the area of a pixel, so that the probability of further target captures can be assumed to be independent from the presence of other captured molecules. Let's consider a scenario where $N_{sp}$ spots have been detected in a field of view (FOV) subdivided in $N_{pix}$ pixels. Assuming that the probability of capturing a molecule is homogeneous within the sample and if $N_{sp} \ll N_{pix}$, the expected number of fluorescent molecules captured within any pixel is approximately $\sim N_{sp}/N_{pix}$.

From Poissonian statistics, it is immediate to compute the probability for a pixel to host two molecules as $e^{-\frac{N_{sp}}{N_{pix}}}(N_{sp}/N_{pix})^2/2 \approx (N_{sp}/N_{pix})^2/2$. On a field of view with $N_{pix}$ pixels, that corresponds to an average of $N_{sp}{}^2/(2N_{pix})$ pixels hosting two molecules within their area.

Given the finite resolution of the setup, two molecules might be detected as a single spot if they are less than 3 pixels apart in any direction. In other words, given a detected molecule, a second one might not be recognized and counted as a distinguished molecule if it sits in a $5 \times 5$ pixels grid centered on the first one.

So, to estimate the correct number of double events we have to consider, instead of the total number of pixels in the FOV, the number of $5 \times 5$ pixels grid, which is $N_{pix}/25$. This gives a $25 \times N_{sp}^2/(2N_{pix})$ average number of molecules in the FOV which are not being counted due to their overlap with other target's signals. Finally, the proportion of molecules that are not being counted with respect to those which are, will be $\sim 25 \times N_{sp}/(2N_{pix})$, which turns out to be $0.008\%$ for 5 aM and $0.45\%$ for 1 pM (based on the average of experimental data).

It's important to notice that this calculation holds only in the case $N_{sp} \ll N_{pix}$ (verified in the whole aM-fM range), because we are neglecting triple or higher order overlap events and treating each additional undetected molecule as statistically independent. Effects such as target molecules depletion in the sample can only make further captures even less likely, diminishing the expected number of double events, so that this can still be considered a good upper bound. Also, if only a proportion $\alpha$ of the total surface is actually properly functionalized and hence capable of target capturing (let's say $30\%$, or $\alpha = 0.3$) the final result will have to be divided by $\alpha$, leading to the formula $\sim 25 \times N_{sp}/(2\alpha N_{pix})$.

#### Supplementary Note. 3

**Table S1.** The list of buffers with recipes.

| Buffer | Recipe |
|---|---|
| FoB 5 | 10 mM Tris-HCl, 1 mM EDTA, 5 mM $MgCl_2$ |
| FoB 20 | 5 mM Tris-HCl, 1 mM EDTA, 20 mM $MgCl_2$, pH 8.0 |
| Gel Buffer | 40 mM Tris, 20 mM Acetic acid, 1 mM EDTA, 12.5 mM $MgCl_2$, pH 8.0 |
| PB | 40 mM Tris-HCl, 40 mM $MgCl_2$, pH 8.4 |
| PBTween | 40 mM Tris-HCl, 40 mM $MgCl_2$, 0.05% Tween 20, pH 8.4 |
| TE2MNaCl | 10 mM Tris-HCl, 1 mM EDTA, 2 M NaCl |

**Table S2.** Folding programs used for the folding of the DNA origami nanostructures.

##### Folding program for Triangle and Trident DNA origami nanostructures

| Temperature (°C) | Time per °C (min) | Temperature (°C) | Time per °C (min) |
|---|---|---|---|
| 65 | 2 | 44 | 75 |
| 64-61 | 3 | 43 | 60 |
| 60-59 | 15 | 42 | 45 |
| 58 | 30 | 41-39 | 30 |

| | | | |
|---|---|---|---|
| 57 | 45 | 38-37 | 15 |
| 56 | 60 | 36-30 | 8 |
| 55 | 75 | 29-25 | 2 |
| 54-45 | 90 | 4 | Storage |

**Table S3.** The list of unmodified staples for the Trident.

| Plate No. | Well no. | Sequence (5' to 3') |
|---|---|---|
| Plate 1 | A1 | CCCCCTGATATTCAACCGTTCCAA |
| Plate 1 | A2 | ATCAAAGGGTGATTAAGACGGAATAGGAAACCAGA |
| Plate 1 | A3 | CAAAATCGGCCAACGCGCGGGGTGGAA |
| Plate 1 | A4 | CGCCTGTGCAGGTAATGGCATCAGCGGTGGTGCCA |
| Plate 1 | A5 | CCTATAAATCCAGGTTGAAGCCCCCAATAGCGTCA |
| Plate 1 | A6 | TTCATACATAAGCTTGAGA |
| Plate 1 | A7 | CGAGTTGGGAAGAAAAATCCCCCC |
| Plate 1 | A8 | CCCCCCAGTATGTTAGCAAACGAAAGCGCATTAGACCCCCC |
| Plate 1 | A9 | CCTTCCAGTAAGCGTCTCAGTGCAGGCGGATAA |
| Plate 1 | A10 | CCCCCAGGGCGATCGGTAAGGGGGATGTGCCCCC |
| Plate 1 | A11 | CCCCCATTTCTGCTATCGACATACCCCC |
| Plate 1 | A12 | AAAACAGGTCTCCAGAGCCACCACCCCACCCTTAC |
| Plate 1 | B1 | CCCCCCTCATTTTCCAGACGATTGGCCCCCC |
| Plate 1 | B2 | TCATAGGTCTGAGAGACTACCCCC |
| Plate 1 | B3 | CTTGTGTCACCAGTTGAGGATCCCAAGCCGGCTTT |
| Plate 1 | B4 | CCCCCCCGGAAACCAGGCAAAGCGCCATTCGTAAGCTTTC |
| Plate 1 | B5 | CACCTTGCTGGTAATATCCAGACCCCC |
| Plate 1 | B6 | AGTGTTTACCGGCCACCAACCGGAATTACCCTGAC |
| Plate 1 | B7 | ACCGCCAGACAGAAGTATAGCCCGGACGTCGAGAAGTTTTAACG |
| Plate 1 | B8 | CTAGGGCGCTGGGTTTCTGGGCCGTTTTCACGGTCCCCC |
| Plate 1 | B9 | ATCAAAAGCCTCGCTTTCCAGTCGGGTGAGACGCA |
| Plate 1 | B10 | CCCCCCGCTAACGAAAATAAACACCCCC |
| Plate 1 | B11 | ATGAAGGGCGATAAAGAACGTGGACTCCCCCC |
| Plate 1 | B12 | CTTTCCTGAATCTTACCAACCCCC |
| Plate 1 | C1 | AACGTATCACCGTACTCACAGTACCCTTGAGTAAC |
| Plate 1 | C2 | CCCCCACGACGGCCAGTACGGATAACCTCCCCCC |
| Plate 1 | C3 | GCTTAAGAGGTCGTACCTTTAATTGCTCCCCCC |
| Plate 1 | C4 | AGCTCATATGGGTAATCGGAGCAACTATCAGGCTA |
| Plate 1 | C5 | CCCCCCAAGAAACATTTTTAAGACCCCC |

| | | |
|---|---|---|
| Plate 1 | C6 | GCCAACGGCATTTAAAAAATCCTTCCGTAATGGGA |
| Plate 1 | C7 | AACAATTTCATTTGAACCAAGTTTTCAGGTCCGAC |
| Plate 1 | C8 | AGGCCCTGAACAAGAAAAAGTAATTAAATTGCTCC |
| Plate 1 | C9 | CCCCCTTTGTCACAATCAGACAAAAGGGCCCCCC |
| Plate 1 | C10 | CCTTCCTGTAGCTTAATTATAAAGCCCCTCATATA |
| Plate 1 | C11 | GAATGAGTAACAACCCGTCGGATTCTCCCCCC |
| Plate 1 | C12 | CCCCCAATGCTGTAGCTCAACATGTTTTATATGGCTTAGA |
| Plate 1 | D1 | CGTCGGGGTCCGCCGCTGGAAGAAAGCGAAACTGT |
| Plate 1 | D2 | CCCCCGGTGAATTATCACCGTCACCGACTGAAATATTGAC |
| Plate 1 | D3 | TATTCACGTTGCGTTAGTAAATGAAAACACTGCAC |
| Plate 1 | D4 | TCACTCTGTCCGACAGGAGAATCAGCTAAAGGGAG |
| Plate 1 | D5 | CCAATAAAGCGAAGGAAGCAGCGGATAATT |
| Plate 1 | D6 | TGATTGTTTGGATTATACAAACAGATTAT |
| Plate 1 | D7 | AACTGACCAACCTGATATACGTAACAGCATCCTT |
| Plate 1 | D8 | TTATGCTTTCCTCGTTAACGGTACTGTGTTCGTTG |
| Plate 1 | D9 | GACGATAGTGAATTTATCGAAAGCGA |
| Plate 1 | D10 | AAAGGAATTGTGGCTATGTAATAAAAGGGACTGAG |
| Plate 1 | D11 | GGCGAGAAGAACTCAAACAGGAAAATGAGGCGGTC |
| Plate 1 | D12 | CCCCCCTGCAAGGCGATCGACGTTGTAAACCCCC |
| Plate 1 | E1 | GGCGTTTTAGCGAACCTCCCCC |
| Plate 1 | E2 | ATCATTGTTTGCCCTACCG |
| Plate 1 | E3 | CCGCGACAACTTAATACATGAGCCGATGCGGCGCC |
| Plate 1 | E4 | ATTCATTATCAGGACACT |
| Plate 1 | E5 | TAAACTGAAAGCGTAAGAATACGTTTTAGGAGTTT |
| Plate 1 | E6 | GAGCCTTGAATGACCCTCC |
| Plate 1 | E7 | CCCCCCATACCGGGGCAAGTGTAGCGCCCCC |
| Plate 1 | E8 | CTGAACGAACCACTTTTGACGCT |
| Plate 1 | E9 | AGAGTTTTTTGGGGTCGCTCA |
| Plate 1 | E10 | AAGCATACCGATCTGACCTAAATTTAGAAAACGGT |
| Plate 1 | E11 | TCTGTGCTGCGGCCAGAGGTCACTGCGCTTTGAAT |
| Plate 1 | E12 | CCCTGTAATACGCATTAACCCATCCTAAT |
| Plate 1 | F1 | GTATTTTGCGTTGAATATTACC |
| Plate 1 | F2 | GCCTCCTGAGTATAACGGAGCTTGACGGGGACGGG |
| Plate 1 | F3 | AAGAGAATACGAGCATTACTAATAGTA |
| Plate 1 | F4 | GACTATAGAAATTTCAACAGTTTCAGCCCCCC |
| Plate 1 | F5 | GGAACTGCTCCATGTTACTTAGCATCCAAGACTTT |
| Plate 1 | F6 | GCTCTCTGTGTCGCGTCCGTGAGCAAGGAAGACAG |

| | | |
|---|---|---|
| Plate 1 | F7 | CAACAATAACAATAAGCA |
| Plate 1 | F8 | TAGCTTAGATTAAGACGCTCCCCC |
| Plate 1 | F9 | CCCCCAGAATCAAGTTTGCCTTAAAGAACAAATAAAAGAGA |
| Plate 1 | F10 | CCCCCACATTAATTGCGTTGCGCTTGCCCTTGGTC |
| Plate 1 | F11 | CCCCCCTTTTGATAATTGCTGAATATCCCCC |
| Plate 1 | F12 | TCGTCGCTATTAATTAATTCCCCC |
| Plate 1 | G1 | CTGCAGGTAGCGACGATATAGCGTCCAATACTTG |
| Plate 1 | G2 | GATTAAACAGTTAATGCCATGGAAAGCCGCCGCAT |
| Plate 1 | G3 | TCATGCAAAGACACCACGGCAA |
| Plate 1 | G4 | TATAGAAGGCTTATCCGCCCCC |
| Plate 1 | G5 | CCCCCTGAAGGGTAAAGTTAAATTTTA |
| Plate 1 | G6 | CCCCCCAACAGGTCAAGTACGGTGTCCCCCC |
| Plate 1 | G7 | TTTTTTAATGGAAACATTCGCCTATATACAGTAAT |
| Plate 1 | G8 | AACTAAGGATTAGACCGGAAGCAAACTCCCCCC |
| Plate 1 | G9 | GAGTTGCCCCAGGGCAACGCAAATGAAA |
| Plate 1 | G10 | CCCCCCCTTTTTAACCTCCGCTTCTGGTGCCCCC |
| Plate 1 | G11 | CCCCCCCTCAGGAAGATCGCACTCCAGCCTTCCTTGAGGG |
| Plate 1 | G12 | TCAAACATTACGCGCAGCATTT |
| Plate 1 | H1 | CCCCCACAAAGAAACCACCAGATTATCATATTAATGCAC |
| Plate 1 | H2 | TTTTGATAAAGTTATACATGCCTGAGTAATGTGTAGCCCCC |
| Plate 1 | H3 | ATTAGCTCATTATACCAGCCCAAATCAGACCAGGC |
| Plate 1 | H4 | CCCCCAGATTAGTTGCTATTTTGCACCCATAAGCAATAGC |
| Plate 1 | H5 | AAGGAAATGCAAAATTCTTCATAATACGTACAGAG |
| Plate 1 | H6 | TTAAGCTAAATTTCAATTCAAG |
| Plate 1 | H7 | TTCATCAACTAGGCATAGTCCCCCTCAAATGCTTG |
| Plate 1 | H8 | CCCCCACCGGAAACAATCCGGAATTTCCCCC |
| Plate 1 | H9 | CCCCCGGAGTGAGAGTAGCGCGTTTTCCCCC |
| Plate 1 | H10 | CCCCCCATCGTAACCGTGCATCTGCCAGTAGGGAGGTCAC |
| Plate 1 | H11 | CCCCCTTCCCTTAGAATTGTAGATGGGCGCCCCC |
| Plate 1 | H12 | AAGGTGCATCATTATTAGCGTTTGCCCAGCATAAC |
| Plate 2 | A1 | TTTTCCAGCTATATTTTCAAGCAAATCAGAACTTA |
| Plate 2 | A2 | CCCCCTCAGCTCATTTTTTAACTCGATGAACTACCCCGCAG |
| Plate 2 | A3 | CCCCCGAGAGTTGCAGCAAGCCACCGCCTGGCCCTGACCCCC |
| Plate 2 | A4 | CCCCCAGAGCCAGCAAAATCACCAGTAGCGAATTTTTGCG |
| Plate 2 | A5 | TAGCGAAGCCCATGAAATAGCCCAATAATAAGAGCCCCC |
| Plate 2 | A6 | AGTAGCCAGCAAGCTGATCACTGCCGGGGTGCCTA |
| Plate 2 | A7 | AGAAACCGAGTAAAAGAGACGACCATAGTCTTTAA |

| | | |
|---|---|---|
| Plate 2 | A8 | CGCCCATTCAGGCTGCGCAACTGTTGGGACCCCC |
| Plate 2 | A9 | GTTGGCCTTGAAAACATTGGGGTAAA |
| Plate 2 | A10 | CCCCCCTTGATATTCACAAACAAATTATTCTGAAACCCCCC |
| Plate 2 | A11 | CCCCCCGGGTATTAACTTCAAATATCCCCCC |
| Plate 2 | A12 | ATCGGCACAGCCAACAGAGATAGACACGCAACCAG |
| Plate 2 | B1 | CCCTAGCAATACTTCTTTCGTCTGATAAAAATACC |
| Plate 2 | B2 | CTCAGAGCCGCCACCCAGTTCAGAAAACGATAATT |
| Plate 2 | B3 | CAGGTGAACCATCACCCCGAGCCGGTCGTGCC |
| Plate 2 | B4 | TGAATAAGAGCAACACTATAG |
| Plate 2 | B5 | CCCCCCAACGTCAAGTGAGCTAACTCCCCCC |
| Plate 2 | B6 | TAAACACCGTTTGAATTTCAGAGGTTTTCCCAGTCATAAG |
| Plate 2 | B7 | TTGCGCGAGGCTGTCTTTCCTTCTAATTTAAGTA |
| Plate 2 | B8 | GTTGATCGGAACGAGGCGTAG |
| Plate 2 | B9 | GTATCATCGCTTTGAATCA |
| Plate 2 | B10 | AGCCTTTATTCAATTCGTAGAAACCAA |
| Plate 2 | B11 | CGGCACCGGCTTAGGTTGGCGCAAAA |
| Plate 2 | B12 | CTATAATCAGATTCTGGACAATATTTTTGAAAGGA |
| Plate 2 | C1 | CCCCCAGTATCATATGCTTAATGCCGCCCCC |
| Plate 2 | C2 | AGTGAGAATCGCCATGCTTGAGAGCATGTTTAACG |
| Plate 2 | C3 | GAGATGGTTGCGAACGTGGCGAGACTCCTCATGCG |
| Plate 2 | C4 | CCCCCGTCACGCTGCGCGTAACCAAATCCGCCGGGCCCCCC |
| Plate 2 | C5 | ATGTTCCACACAACATAAAATCAATAGGGTT |
| Plate 2 | C6 | CCCCCGTATTCTAAGAACGCGATTAGAAACGCATAAAACTA |
| Plate 2 | C7 | CGGTAAAGCCGCACAGGCGGCCTTTAGTGACCCCC |
| Plate 2 | C8 | GCCCTAAAACACAATATCGAAGAGGCGGTTTGAAT |
| Plate 2 | C9 | CCCCCTAGATTTAGTTTGACCATTAGATAACATTTGATTC |
| Plate 2 | C10 | AGACGACGGATAAGTAAGGCAAAGAA |
| Plate 2 | C11 | ATTGTTATCCGAGGTGCCAATCAAAAGAATAGTCG |
| Plate 2 | C12 | ATCCAAAAAGAGATTTTTTCGTCTCGTCGCCCCCC |
| Plate 2 | D1 | CTGGAAAAACCAAAATAGGAACAACGAAAGAGCGC |
| Plate 2 | D2 | CCCCCAGGAAGATTGTATAAGGAAA |
| Plate 2 | D3 | TTTGAAGCCTTAAATCACCCCC |
| Plate 2 | D4 | CCCCCCCCGACTTGCGGGAGGTTAATTTGCCCAATCCAAA |
| Plate 2 | D5 | CCCCCAAAATAATTCGCGTCTGCTACAAAGGACAAGAGCAC |
| Plate 2 | D6 | CCCCCGAGAAGAGTCAACGACAGTATCGGCCCCC |
| Plate 2 | D7 | AACGTACCGTTTTTCTGAATA |
| Plate 2 | D8 | CCATCGCCACCCTCAGAAGAGACTCTATTTCGGAA |

| | | |
|---|---|---|
| Plate 2 | D9 | CATGCTAGAAAATACATACCCCCC |
| Plate 2 | D10 | GGACTTGTAGAACCGCAACGCACTCCCACACCGCG |
| Plate 2 | D11 | TGCGATCAACAAGCAAATGCCAGCGGGTCATAGCT |
| Plate 2 | D12 | GTGCATAGGTGCCTGTAGCGATCTACCAAAAG |
| Plate 2 | E1 | CCCCCATGAAAGTATTAAGAGGCTCCGCCACGCAAGCCAAA |
| Plate 2 | E2 | TTGGAACATTTCGCAAATTACCGCACATCGTAAGA |
| Plate 2 | E3 | TTATAAAGTATTAACGTGAT |
| Plate 2 | E4 | CAGTTAATTTAACAACGCCAACATGAATAACCTGT |
| Plate 2 | E5 | CCCCCCGTGGGAACAAACGGCGGAT |
| Plate 2 | E6 | TGATGATACAGGAGGGCGCCTCAGA |
| Plate 2 | E7 | TGATGCCCGATAGATTATGCG |
| Plate 2 | E8 | GACGAGCACGAAGTGTTCATAAACTTATCTAAAAT |
| Plate 2 | E9 | CCCCCGGGATAGCTCAAACTTAACCCCC |
| Plate 2 | E10 | CAGACAATTCCACGGGAGCC |
| Plate 2 | E11 | TTAAATTTTTGTTAAACCCCC |
| Plate 2 | E12 | CTTTACAGAGGCTTTGAGGACTACTATCGGTTTAT |
| Plate 2 | F1 | GGACACCAACGTCACCCGACAATGACAACAAAGA |
| Plate 2 | F2 | CCCCCGCGTTTTAATTCGAGCTTCTCTGCGAACGAGCCCCC |
| Plate 2 | F3 | TTTTCTCATCGGATTAAGACAGCAGCACCGTAAAT |
| Plate 2 | F4 | CCCCCAATAGCAGCCTTATTTTTTATAGTCATCATAATCA |
| Plate 2 | F5 | ACCCTGAACAAAGTCAGCCCCC |
| Plate 2 | F6 | TCATCGGTTGTACCAAATAC |
| Plate 2 | F7 | TGCTGGAGGTTTCACCAGTTCCAGAAAAATCTCCA |
| Plate 2 | F8 | CCCCCGGTCTGGTCAGCAGCAACGT |
| Plate 2 | F9 | CCCCCTAGACTTTACAACGTGGTGCTCCCCC |
| Plate 2 | F10 | AAAACAAAATTAATTAAGGCGAAAATAAAGCTGTCC |
| Plate 2 | F11 | CCCCCAAAAAATCCCGTAAAATGTGTACCATTTGCAGCG |
| Plate 2 | F12 | ATATACAGAGGGAATCATTACCCCCCC |
| Plate 2 | G1 | CCCCCATTTTGTTAAAATTCGCTGATAATCACAAATATGGG |
| Plate 2 | G2 | CCCCCGCGCCCAATAGCATTTGGGTTTAGAACAACGCTAGT |
| Plate 2 | G3 | GGTCAATCACCGCGACGTTTCCAAACG |
| Plate 2 | G4 | GCAGGTTGCCCGAGCCGTCAATAGACGTATTAGTC |
| Plate 2 | G5 | TACGACGATCCAGCGCATGCTCGTTTTTACGGCTG |
| Plate 2 | G6 | CCCCCTACGTTAATTAAAACACTCATCCCCC |
| Plate 2 | G7 | GGCAATTCATCAATATAAGTAGATTACAAAATTGA |
| Plate 2 | G8 | AAGGTGGCATTCAACGTAACGGAGTACATAAATCA |
| Plate 2 | G9 | AAAAGGTAAATAATATCATCCAATAAA |

| | | |
|---|---|---|
| Plate 2 | G10 | CCCCCCCGGCAAACGCGGTCCGCGGTA |
| Plate 2 | G11 | GGGCCAGAAGGAGCGGAATTATCATACCT |
| Plate 2 | G12 | GTTTCGGAACCAGCGGGAGCTAAACAGGAGTAGTA |
| Plate 2 | H1 | CCCCCCACAAGAATTGAGTGCTACAATTTTATCCAGAGCC |
| Plate 2 | H2 | CGAAATTAAGGGAGACGAGAAACACCAAAT |
| Plate 2 | H3 | TTGTTTTTCACGCAAGACAAAGAAGTTATATTCTT |
| Plate 2 | H4 | TGAGCCATTTGGGAATTCCCCC |
| Plate 2 | H5 | GCATTACCAAGGCAAAAGAAAGGCCCCACGCATAA |
| Plate 2 | H6 | AGCACGCGTGCGGAGCGGCGCCGCGCTTAATGATT |
| Plate 2 | H7 | ATATCAGAGAGATAACCCCCCC |
| Plate 2 | H8 | CAACGGAACAAACAGGGAGCCGTTTTGGCATGAGA |
| Plate 2 | H9 | CCCCTAATCATGTGCCGGTGCCCCCACACTGGGCC |
| Plate 2 | H10 | AAGCACTAAATCCTGTGTCCGGGTTACCTGCACGT |
| Plate 2 | H11 | GAACGTGCTTGCCAGAGAACAATAGGAACGCCATCACCCCC |
| Plate 2 | H12 | CCCCCGTGAGAGATAGACTATACCAGTCCGGCGAATACTAGATAAGAA |
| Plate 3 | A1 | TCACGAGCCAGTAACAGTCATA |
| Plate 3 | A2 | CCCCCGCCATATTAGTTTAACGTCAAAAATGAACCCCC |
| Plate 3 | A3 | TACCAGAAAAGATTACGAAGGGATTT |
| Plate 3 | A4 | AATTGTAAACGTTAAT |
| Plate 3 | A5 | CCCCCCTGAGGCTTGCAGGGAGTTAATACACAAAA |
| Plate 3 | A6 | ATAAAGGCTAAGTTTTGTCGTCTTACAAACCAGAG |
| Plate 3 | A7 | TTGGCTTAAATGTGAGCAACCTTGCTTCTAATACC |
| Plate 3 | A8 | ATCATGGTCAGTTGGCAACGAACTGGATTCACCAG |
| Plate 3 | A9 | CTTTGAAGCAACCGAAAGAACC |
| Plate 3 | A10 | CCCCCAACGTCACCAATGAAACCATCGATCAGAACCATTA |
| Plate 3 | A11 | TAAATTTGTACTGGTAATAGG |
| Plate 3 | A12 | GGATTATTTACCCGTTGTTAGCCGATTAAAGGGGC |
| Plate 3 | B1 | AACGACGCCAGCTGGCGAGCGG |
| Plate 3 | B2 | CCCCCTGGAAGTTTCATTCCATATAACAGGGGGAATATGC |
| Plate 3 | B3 | ATAAATTTTTTTTTATCCAGTTACAGCGT |
| Plate 3 | B4 | TAAAGCCAGACCCTGCCCTCAAGAGAAGGATTCAG |
| Plate 3 | B5 | CCGAGGGGGTACTTTTGCAAAAGAAGTCCCCC |
| Plate 3 | B6 | GCCAACTATATGTAAATGCTGACCCCC |
| Plate 3 | B7 | CCCCCCATCGGCATTTTCGGTCATGGCAGGTAGGG |
| Plate 3 | B8 | GTCATAGTTAGCGTAACATTCCACACCCTCGCTTT |
| Plate 3 | B9 | AAGGGTATCATTCCAAGAACCCCC |
| Plate 3 | B10 | GCAACGTATTGATCAAACCCTCAATTCGCCATAAT |

| | | |
|---|---|---|
| Plate 3 | B11 | TTTACATACGGCAGAGGCATTTTATAATCGCTGAA |
| Plate 3 | B12 | ATATATGTGCACGGGAGAAACAACAAGGATAAAAA |
| Plate 3 | C1 | ATGTTTCCATCGCGCTTTTGCGGGATCCTAAAACATT |
| Plate 3 | C2 | GGCGCGTACTGTGTCCAGGTAAAGGCACTAACAAC |
| Plate 3 | C3 | AAACCGCCAGCAGCGATGCTGATTGCCGTTCCCCC |
| Plate 3 | C4 | CCCCCTGGCAGCCTCCGGAGTAACCTTTCATCAACAGCATG |
| Plate 3 | C5 | GCTCGAGGTGAATTTCTCAT |
| Plate 3 | C6 | CCATTGAGGGAATTTACCAGCGC |
| Plate 3 | C7 | TTCCTGATTATCAGATCAGATGAGATTGCTGGAGA |
| Plate 3 | C8 | CCCCCCTTTGACCCCCAGCGATTAAGGCTGGCCGGAT |
| Plate 3 | C9 | TAACGATTTTAATCACGCAAATTAAATTGGCAATA |
| Plate 3 | C10 | ATACCTCAGAGCCACCACCCCCCC |
| Plate 3 | C11 | CCCCCGTAAAGATTCAACCATCAATACCCCC |
| Plate 3 | C12 | GGGATACATGAGAGCCAGGAACCGCATAAATCAAAA |
| Plate 3 | D1 | AGGATCCCTTTGCATCACGAGCTCGAATTCGCGAT |
| Plate 3 | D2 | TCCACCCTTCTGACCGTTTTTGCGGACCCCC |
| Plate 3 | D3 | CAAAATAGAAATCAGTAGCGACCCCCC |
| Plate 3 | D4 | CCTGAAGCATAAAGTGTCCACTACTTTGGAACAA |
| Plate 3 | D5 | TGGTTTGAACGAGCAGAC |
| Plate 3 | D6 | CCCCCGCGGTTGCGGTATTTGAGGATTTAGAAGTATCCCCC |
| Plate 3 | D7 | GATCTCACGGTCTTCTCCGTGGTGAACCCCC |
| Plate 3 | D8 | GTGAATAAGGCGAATTACTGAGATTTCATAACTCG |
| Plate 3 | D9 | CAACTATCGGCGCTGGTTCCACTATAAAACCGTCT |
| Plate 3 | D10 | CAGCTTTAAACAAAAGGAATTACGAATGCAGATGA |
| Plate 3 | D11 | CAGAGCGCAGTCTCTGAACCCGTATAGCGGGGTTT |
| Plate 3 | D12 | CCCCCGGGAGAATTAACTGAACTAACCAGAACCCAAAAGA |
| Plate 3 | E1 | GGTAGCTTAAACGACCACATACTTTA |
| Plate 3 | E2 | GCGTATGGGATTTTGCTCAATAGGTGACAGGTCAT |
| Plate 3 | E3 | CCCCCGTAATCTTGACAAGAACTGACCTTCATCAAGACCCCC |

**Table S4.** The list of modified staples for Trident.

| Name | Sequence (5' to 3') | Replace |
|---|---|---|
| 3' biotin | ATAAAGGTGGAATAAGTTTAT-Biotin | Plate 3, F1 |
| 3' biotin | TGAGAGTCTGTAAAACTA-Biotin | Plate 3, F2 |
| 3' biotin | AAAGTAAGCGAGGAAACG-Biotin | Plate 3, F3 |
| 3' biotin | GACATTCAACCGTTATTCATTAAA-Biotin | Plate 3, F4 |

| | | |
|---|---|---|
| 5' biotin | Biotin-AGGGTAATTGAGCGCTATATCTTACCCGAACAAAG | Plate 3, F5 |
| 3' biotin | GAGAGGGTAGTCATTGCC-Biotin | Plate 3, F6 |
| 3' biotin | TTAATTTCATCTCCGTGTGATAAA-Biotin | Plate 3, F7 |
| 3' biotin | CAATAATAACTCCTTATTACG-Biotin | Plate 3, F8 |
| 3' biotin | TGCAAATCCAATAATATATTTTAG-Biotin | Plate 3, F9 |
| 3' biotin | GGAAAATTGAGGAGCAAGGCCGGA-Biotin | Plate 3, F10 |
| 3' biotin | GCATGTCAACCCAAAAAC-Biotin | Plate 3, F11 |
| 3' biotin | TAAGGCGTTAAAAAAAGCCTGTTT-Biotin | Plate 3, F12 |
| 3' base dye | AAATCGAACCACAGTTTCGTAGTACCGCCACCCTAG-ATTO 542 | Plate 3, G1 |
| NP binding 3' | AATCACAGAGGACGCTCATGGAAATCCTGAG-TAAAATCCGTTCAAAAAAAAAAAAAAAAAAAAAAAAA | Plate 3, H1 |
| NP binding 3' | AGCTGCGGGTGGTTGGTGGTAA-TAACAAAAAAAAAAAAAAAAAAAAAAAAAAA | Plate 3, H2 |
| NP binding 3' | GAGTGTTTGTTTGATTTTCTTTCACCTT-GAAAAAAAAAAAAAAAAAAAAAAAAA | Plate 3, H3 |
| NP binding 3' | GAACCTCAAATGGCGCCAATTAAAAAAAAAAAAAAAAAAAAAAAAAA | Plate 3, H4 |
| NP binding 3' | GAATGGTAAAAAATTGTGTAAAAAAAAAAAAAAAAAAAAAAAAAA | Plate 3, H5 |
| NP binding 3' | GAAACAAAGTACGGTGTACAACGTAACAAA-GCAGAAAAAAAAAAAAAAAAAAAAAAAAAAA | Plate 3, H6 |
| NP binding 3' | TCATAAATATTTAAACAGGGAACGA-GAAAAAAAAAAAAAAAAAAAAAAAAA | Plate 3, H7 |
| NP binding 3' | CGCCTCAGCAGCGAAAGATGCCACTCATCAG-TCTTATGCAAAAAAAAAAAAAAAAAAAAAAAAA | Plate 3, H8 |
| NP binding 3' | GATACCGATAGTGCGGAAC-CTCGTTAAAAAAAAAAAAAAAAAAAAAAAAA | Plate 3, H9 |
| NP binding 3' | CAATGATTAGTTCCGAAACCCGAAAAAAAAAAAAAAAAAAAAAAAAA | Plate 3, H10 |
| NP binding 3' | TCACTTGACCTACACAGCAGAAGATAAATAAAGCATTCAC-CAGAAAAAAAAAAAAAAAAAAAAAAAAAAAA | Plate 3, H11 |
| NP binding 3' | GATTTTATGCTCATAGAG-GACAGATGAACAACAAAAAAAAAAAAAAAAAAAAAAAAA | Plate 3, H12 |
| Hybridization to Triangle | ATGTAGGTGGTAGAGTTCACAAGAATTGAGTGC-TACAATTTTATCCAGAGCC | Plate 2, H1 |
| Hybridization to Triangle | AGCAAGTCCATTACCAAGGATTGGTGAATTATCACCGTCACCGACTGAAA-TATTGAC | Plate 1, D2 |
| Hybridization to Triangle | GCGCGGTGCAGTCTCGTCCTTTAAAAAATCCCGTAAAATGTGTACCATTT-GCAGCG | Plate 2, F11 |

| | | |
|---|---|---|
| Hybridization to Triangle | CTTGCCAGCATTGTAATAGGTTAGGGCGATCGGTAAGGGGGATGTG | Plate 1, A10 |
| Hybridization to Triangle | CACTAAAAGAGTGATGATAATTATGAAAGTATTAAGAGGCTCCGCCAC-GCAAGCCAAA | Plate 2, E1 |
| Hybridization to Triangle | TCATGAGTGCCGAGCTAAGATTGTCACGCTGCGCGTAAC-CAAATCCGCCGGGC | Plate 2, C4 |
| Capture Strands | TGTGCCTGTTTATCAAGTTTAAGCCTCAGAGCATAAGCAAAATGTTTAT | Plate 3, G5 |
| Capture Strands | ATTTACAACATGTTCAGCTAATGTTTTGTGCCTGTTTATCAAG | Plate 3, G3 |
| Capture Strands | CAGAACGCGCCTTAAGCAATATTTTGTGCCTGTTTATCAAG | Plate 3, G2 |
| Capture Strands | TACCATATCAAAGCAAAAGAATTTTGTGCCTGTTTATCAAG | Plate 3, G4 |
| Capture Strands | CGTATTCTGAATAATGGAAGGGTTAGAACCTTTTGTGCCTGTTTATCAAG | Plate 3, G6 |
| Capture Strands | TGTGCCTGTTTATCAAGTTTGATGATGAAACAAACATACCTGAATT | Plate 3, G7 |
| Capture Strands | TGTGCCTGTTTATCAAGTTTACAATATTACCGCGCCTGCAA | Plate 3, G8 |
| Capture Strands | CAGTGCCACGCTGAGATTAACACCCAGCCATTGTTTT-GTGCCTGTTTATCAAG | Plate 3, G9 |
| Capture Strands | TGTGCCTGTTTATCAAGTTTTTTGCCAGATATATTCGGTCG | Plate 3, G10 |
| Capture Strands | AACGCGAGAGGATAGTAAATTTTGTGCCTGTTTATCAAG | Plate 3, G11 |
| PAINT docking | CATACCGGGGCAAGTGTAGCGTTTTAACATTCC | Plate 1, E7 |
| PAINT docking | AGTAGCCAGCAAGCTGATCACTGCCGGGGTGCCTATTAACATTCC | Plate 2, A6 |
| PAINT docking | CAACGTCAAGTGAGCTAACTCTTTTTTTAACATTCC | Plate 2, B5 |
| PAINT docking | CTAGGGCGCTGGGTTTCTGGGCCGTTTTCACGGTTTTTAACATTCC | Plate 1, B8 |
| PAINT docking | CGTCGGGGTCCGCCGCTGGAAGAAAGCGAAACTGTTTAACATTCC | Plate 1, D1 |
| PAINT docking | GAGAGTTGCAGCAAGCCACCGCCTGGCCCTGATTTTTTTAACATTCC | Plate 2, A3 |

**Table S5.** The list of sequence for staples involved in the sandwich assay.

| **Name** | **Sequence (5′ to 3′ end)** |
|---|---|
| Target | TTCGAATACCACCGTCGAGCCAGAAACTGTCTACATTGCCCGAAATGTCCTCATTACCA-TAATCGAAAGCATGTAGCATCTTGCTCATACGTGCCTCGCCAATTTGGCGGGCAAATTCTTGAT AAACAGGCACAACTGAATATTTCATCGC |
| Imager | GCGATGAAATATTCAGT-Alexa Fluor 647 |
| Blocker | ACTTGATAAA |
| Displacer | TGTGCCTGTTTATCAAGAATTTGCC |
| Sacrificial DNA | GTGATGTAGGTGGTA |

### References


1. Close, C. *et al.* Maximizing the Accessibility in DNA Origami Nanoantenna Plasmonic Hotspots. *Adv Mater Interfaces* **9**, 2200255 (2022).

2. Douglas, S. M. *et al.* Rapid prototyping of 3D DNA-origami shapes with caDNAno. *Nucleic Acids Res* **37**, 5001–5006 (2009).

3. Shetty, R. M., Brady, S. R., Rothemund, P. W. K., Hariadi, R. F. & Gopinath, A. Bench-Top Fabrication of Single-Molecule Nanoarrays by DNA Origami Placement. *ACS Nano* **15**, 11441–11450 (2021).

4. Liu, B. & Liu, J. Freezing Directed Construction of Bio/Nano Interfaces: Reagentless Conjugation, Denser Spherical Nucleic Acids, and Better Nanoflares. *J Am Chem Soc* **139**, 9471–9474 (2017).

5. Jing, X. *et al.* Solidifying framework nucleic acids with silica. *Nat Protoc* **14**, 2416–2436 (2019).

6. Wassermann, L. M., Scheckenbach, M., Baptist, A. V., Glembockyte, V. & Heuer-Jungemann, A. Full Site-Specific Addressability in DNA Origami-Templated Silica Nanostructures. *Advanced Materials* **35**, (2023).

7. Nečas, D. & Klapetek, P. Gwyddion: an open-source software for SPM data analysis. *Open Physics* **10**, 181–188 (2012).

8. Schnitzbauer, J., Strauss, M. T., Schlichthaerle, T., Schueder, F. & Jungmann, R. Super-resolution microscopy with DNA-PAINT. *Nat Protoc* **12**, 1198–1228 (2017).

9. Trofymchuk, K. *et al.* Addressable nanoantennas with cleared hotspots for single-molecule detection on a portable smartphone microscope. *Nat Commun* **12**, 950 (2021).
